# Convergent Signals Encode Persistent Memory of Resilient States

**DOI:** 10.64898/2025.12.15.694462

**Authors:** Teresa Femenia, Hannah Prendeville, Na Sun, Cathal Harmon, Kyriakitsa Galani, Kevin Grove, Lydia Lynch, Manolis Kellis, Leandro Z. Agudelo

## Abstract

Caloric restriction reduces metabolic disease and associated comorbidities. Yet, the molecular mechanisms encoding cellular memory of these benefits remain unclear. Here, we use a functional genomics approach to integrate evolutionary cues, single-cell sequencing, and metabolomics, identifying convergent signals that encode epigenetic memory of adaptive states in visceral adipose tissue natural killer (NK) cells. Cross-species analysis shows that genomic hubs linked to human accelerated regions (HARs) are conserved and active during dietary restriction, forming transcriptional compartments that regulate oxidative stress response and DNA repair genes. Targeted screens and multimodal profiling across human data and mouse models reveal convergent signals needed for persistent adaptation: cooperative transcriptional regulators (NRF2, CIRBP, NR4A2) at key HAR-linked genomic hubs, innate immune mediators (IL15-IL2RB), and metabolic cofactors from linoleic acid oxidation. Activity of these convergent signals reduces DNA damage and methylation and enhances epigenetic plasticity and cytotoxic functions, thereby decreasing tissue fibrosis and senescence while maintaining both local and systemic metabolic plasticity. Mechanistically, linoleic acid metabolism supplies acetyl-CoA to sustain H3K27ac epigenetic marks that maintain coordinated repair and cytotoxic programs after programming. IL15-IL2RB signaling links tissue metabolic state to NK cell function for coherent adaptive responses. We show that engineering of NK cells with this multimodal programming leads to long-term preservation of therapeutic phenotypes in metabolic aging models, rescuing systemic dysfunction through damaged cells clearance and leptin sensitivity. Overall, these findings establish that convergent signals encode persistent cellular phenotypes linked to metabolic plasticity, suggesting design principles for engineering therapeutic memory of cell function.

**One-Sentence Summary:** A multimodal approach reveals how dietary and evolutionary cues can be harnessed to engineer resilient phenotype memory.

## Introduction

Caloric restriction extends healthspan and lifespan across species, yet translating these benefits into therapeutic outcomes has remained difficult. CR enhances metabolic plasticity and activates molecular repair programs, yet the molecular mechanisms underlying these adaptations at the cell-level remain unclear. This gap is critical for several reasons but in particular how engineering durable cellular resilience requires identifying not just what CR activates, but how convergent signals encode lasting functional memory.

Millions worldwide are affected by metabolic diseases such as obesity and type 2 diabetes, creating significant socioeconomic challenges, particularly among aging populations. These conditions are marked by reduced metabolic plasticity in nutrient sensing and stress responses. This plasticity is essential for sustaining physiological processes while playing a central role in evolutionary optimization of energy efficiency traits (*1*, *2*). In mammals, metabolic pressures have resulted in traits that are conserved among species with shared molecular mechanisms yet constrained and undergoing molecular optimization. Notably, molecular adjustments affecting metabolic plasticity can influence disease resilience and longevity (*3*, *4*), often through lineage-specific mutations in coding or noncoding genomic regions. While some mutations, such as human accelerated regions (HARs) act as enhancers impacting neural development (*5*, *6*), the roles of other fine-tuned regions in metabolic plasticity and disease remain largely unknown.

Energy conservation is a selective pressure driving physiological and molecular adaptations that shape metabolic diversity and lineage traits such as cognition and endurance in hominids (*7*, *8*). Factors such as larger body size and lower basal metabolic rate have been associated with resilience during food scarcity in mammals (fasting endurance (*9*, *10*)). Molecularly, CR triggers adaptations that appear conserved across mammals and enhance metabolic plasticity (*3*, *4*), including endurance to oxidative stress that protects against molecular damage and genomic instability (*11–14*). Comparative analyses show that the rates of somatic mutations and DNA methylation inversely correlate with lifespan in mammals (*15*, *16*), with longer-lived species showing enhanced genome stability (*17–19*). However, how adaptive strategies to oxidative stress and DNA damage during energy conservation operate at the cell-type level and link to evolutionary adaptations remain unclear.

The plasticity of genome organization during transcription is vital for biological responses to environmental cues, which help define cellular functional identity (*20*). Transcriptional compartments are positioned within distinct chromosome territories, facilitating the interaction between transcription factories and distant chromatin modules for gene coregulation (*21*, *22*). Influenced by multimodal factors such as intrinsically disordered regions (IDRs), phase separation in regulators (*23*, *24*), local environments, and genome structural features (*23*), nuclear compartments regulate specific gene programs and maintain genome plasticity (*21*, *23*, *25–27*). This plasticity influences epigenetic memory, heritable or stable genome modifications by developmental or environmental cues (*28–30*), influencing cell development, transgenerational information, and transcriptional memory (*31*). The latter reactivates previously regulated genes (*32–36*) in response to oxidative and metabolic stress (*32*, *33*), sustaining transcriptional changes for cellular fitness (*31*). These adaptations can be impaired by oxidative DNA damage, genomic instability, mutations, and excessive DNA methylation (*11–14*), contributing to the loss of cell identities in metabolic disease and aging.

In this study, we implement a functional genomics approach to investigate how dietary and evolutionary cues converge to encode genome stability and persistent memory of cell resilience. We use comparative analyses to identify associations between genome stability and metabolic traits in mammals, then validate the molecular mechanisms in disease models. Our results show that CR enhances stress-endurance in visceral adipose tissue (VAT), particularly in natural killers (NK) cells through transcriptional compartment hubs linked to human-specific mutations and genome stability. These hubs, regulated by CR, exhibit conserved activity in murine models, form chromatin contacts for gene coregulation, and influence epigenetic memory of resilient phenotypes. Through single-cell sequencing and metabolomics, we identify critical transcriptional and metabolite mediators of NK cell function post-CR. By engineering NK cells with these multimodal cues, we enhance phenotype robustness, functional activity, and epigenetic memory, thereby improving cell therapy outcomes for metabolic and aged-related disorders. Overall, our study identifies key functional hubs mediating stress resilience and enhancing cell therapy engineering.

## Results

### Relationships between metabolic traits and genome stability in mammals

To identify associations between energy use, energy allocation, and genome stability traits in mammals, we aimed at analyzing published data on metabolic trait parameters (e.g. body weight and body fat composition), somatic mutation rates, and DNA methylation rates. This was complemented by investigating molecular adaptations across murine disease models (**Fig. 1A**). Recent studies have shown an inverse relationship between lifespan and the rates of somatic mutations and CpG-methylation across mammals (*15*, *16*), which suggests that repair mechanisms such as oxidative stress response (OSR) and DNA damage repair (DDR) are conserved across species but fine-tuned throughout evolution. Interestingly, these mechanisms are upregulated after fasting or CR (*11–14*), yet how they lead to durable resilient states remain unclear.

**Fig. 1.**
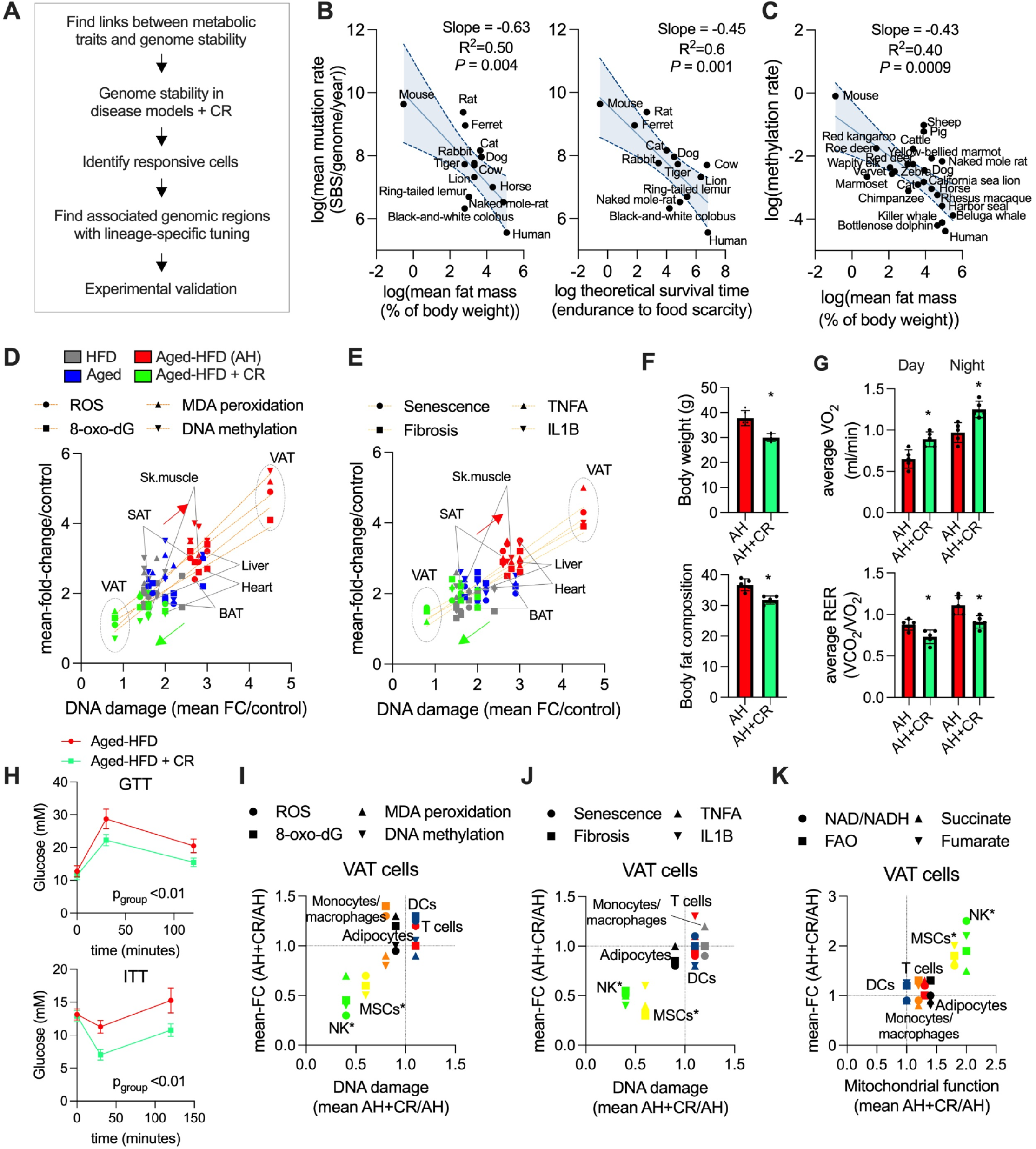
Metabolic traits and genome stability in mammals. **(A)** Illustration of workflow overview. (**B**) Left panel, mean somatic mutation rates across mammals (data from Cagan et al. (*15*)) and body fat composition (BFC) from several curated sources (Supp text and methods), were each transformed using binay logarithmic approach (Methods) and linearized to evaluate dependencies using linear regression (shaded line is 95% CI). Right panel, association with the theoretical survival time scores (Methods) and similar transformation. (**C**) DNA methylation rates (data from Crofts et al. (*16*)) and BFC transformed and analyzed as in B. (**D**) Fold-to-fold change plots for different parameters (y-axis) and DNA damage (x-axis) across tissues. Each mean parameter variation is divided by the mean variation of the control group for that parameter and for each model. Each model is composed of 4-6 mice. Linear regression shows the association between each parameter and DNA damage. (**E**) As in D, fold-to-fold change plots for DNA damage variation and variation of different parameters across tissues. Linear regression for each parameter association. (**F**) Upper panel, body weight (grams), aged-HFD (AH) and AH+CR. Lower panel, BFC, normalized to body weight (n=5) (**G**) Upper panel, average VO_2_ levels (day and night) from 48 hours measurements (at the end of the interventions. Lower panel, average RER levels (n=5). (**H**) Upper panel, glucose tolerance test. Lower panel, insulin tolerance test (n=4-6). (**I**) Fold-to-fold change plots as in D, comparing AH+CR/AH mean molecular variations against DNA damage in cell-types from VAT (n=4-6). (**J**) Fold-to-fold change plots as in I, comparing different parameters against DNA damage variation (n=4-6). (**K**) Fold-to-fold change plots as in I, comparing different metabolic parameters against mitochondrial function variation (n=4-6). Animal experiments were done with n=5-7 mice per group. Data show mean values per group and SEM. Cell extraction experiments were done in each mice per group and mean per groups were compared. Unpaired, two-tailed student’s t-test was used when two groups were compared, and ANOVA followed by fisher’s least significant difference (LSD) test for post hoc comparisons for multiple groups. Two-way ANOVA was used to estimate significance between groups constrained by time measurements, and Tukey test for multiple comparisons. * p-value <0.05.

In our companion study, we found that fat mass strongly correlates with fasting endurance, an estimation of survival time under food scarcity (*9*). Here, we examined the relationships linked to body fat composition (BFC; percentage of body fat relative to average body weight) and genome stability (**fig. S1A**). BFC, unlike the weight composition of other organs such as brain and heart, is inversely correlated with basal metabolic rate per gram, In addition, BFC is increased in hominids and is associated with higher fasting endurance score (**fig. S1B-D,** Table **S1A-D**). In line with previous studies (*8*), this comparative analysis suggests a larger energy budget for metabolic demands.

Assessing the link between metabolic parameters and genome stability traits showed that BFC and the fasting endurance score are inversely associated with somatic mutation rates (**Fig. 1B**), while BFC is also inversely associated with methylation rates in mammals (**Fig. 1C**). Additionally, both BFC and the fasting endurance score are directly correlated with lifespan across mammals (**fig. S1E, F**). These findings highlight conserved links between metabolic plasticity, genome stability, and repair mechanisms in mammals. The association with lifespan suggests that these processes are evolutionarily constrained yet subject to molecular optimization. While further investigation is needed to study in more detail these trait associations, we hypothesized that examining conserved genomic regions with lineage-specific adaptations linked to key repair pathways could uncover functional hubs with broad therapeutic potential.

### Oxidative stress and DNA damage in adipose tissue are reduced under caloric restriction

Oxidative stress plays a key role in mutations, DNA methylation, metabolic regulation, and overall healthspan. Its deleterious consequences (e.g., genomic instability, senescence, fibrosis, cancer) are influenced by oxidative stress endurance, which is modulated by energy conservation (*37–40*). To further explore this, we examined the links between metabolic plasticity, oxidative stress endurance, and genome stability in model organisms under environmental stress. We studied normal (8 weeks) and aged (16-18 months) wild-type mice subjected to a high-fat diet (HFD; 8 weeks) or a combination of HFD (6 weeks) with 2 weeks of CR.

Molecular screening in several metabolic tissues revealed that aging combined with HFD exacerbates oxidative DNA damage, oxidative stress levels, DNA methylation, senescence, fibrosis, and inflammation, particularly in visceral adipose tissue (VAT; **Fig. 1D, E, fig. S2A**). CR improved these molecular parameters in most tissues, especially in VAT (**Fig. 1D, E**). Physiologically, CR reduced body weight gain, BFC, improved metabolic cage parameters (VO_2_, respiratory exchange ratio or RER), and enhanced glucose and insulin tolerance (**Fig. 1F-H**). Plasma levels of triglycerides (TAGs), cholesterol, insulin, and leptin were also reduced (**fig. S2B**), supporting enhanced metabolic plasticity. These results indicate worsening of several parameters in metabolic aging models with these effects being improved by CR, in particular in VAT.

To evaluate cellular responses to CR, we then isolated representative cell types from VAT, including adipocytes, mesenchymal stem cells (MSCs), NK cells, monocytes, macrophages, naïve T cells, and dendritic cells (DCs). Notably, MSCs and NK cells were the cell types that showed the most significant reductions in oxidative DNA damage, DNA methylation, malondialdehyde (MDA) peroxidation, oxidative stress, senescence, fibrosis, and inflammation (**Fig. 1I, J**). These cells also exhibited increased mitochondrial activity with elevated levels of associated outcomes such fatty acid oxidation (FAO), tricarboxylic acid (TCA) cycle metabolites (particularly fumarate), and NAD+ cofactors (**Fig. 1K**). These results indicate that in metabolic aging models, MSCs and NK cells remain responsive to CR, displaying enhanced genomic and metabolic plasticity.

Under stress conditions, MSCs can become profibrotic, senescent, and damaged, triggering immune responses where NK cells play a key role in modulating MSCs proliferation and clearance (*41*, *42*). We therefore focused on the NK-MSCs interplay in our models. This showed that CR reduced the expression of profibrotic and senescent associated secretory phenotype (SASP) gene programs in MSCs from the aged-HFD model (**fig. S2C-E**). CR substantially enhanced colony forming ability of MSCs (**fig. S2F**), indicating improved self-renewal capacity and enhanced clearing of damaged cells.

We next evaluated NK cytotoxic function after CR, which showed increased IFNG production and expression of cytotoxic-associated genes, along with enhanced cytotoxic activity against profibrotic and aged-HFD-derived MSCs (**fig. S2G-I**). Together, these results indicate that CR in NK cells not only enhances resilience to oxidative stress and DNA damage but also improved cytotoxic activity and clearing of damaged MSCs (**fig. S2J**). This suggests that NK repair and cytotoxic functions are enhanced by metabolic plasticity, particularly processes linked to mitochondrial activity and associated cofactors (e.g., NAD+, fumarate), which help maintain genome plasticity (*43*). These experimental observations led us to investigate whether evolutionary tuned genomic regions and transcriptional compartments coordinate these responses in NK cells, potentially mediating organismal resilience.

### Transcriptional hubs with human-specific mutations are linked to OSR and DDR programs in NK cells

Transcriptionally-driven compartments are dynamic organizations that include genomic regions with transcriptional activity, chromatin interactions, cooperative transcriptional regulators, and epigenetic modifications. The integrity of these organizations is influenced by oxidative DNA damage, mutations, methylation, and the activation of repair systems such as OSR and DDR programs (*37–39*). Both of these repair pathways share molecular regulators that are linked to epigenetic plasticity, metabolic disease, and aging (*40*).

This led us to investigate whether these repair pathways share transcriptional, functional, and evolutionary dependencies, providing insights into their broader therapeutic applications. We recently used a functional genomics approach to examine relationships in genomic regions harboring genes linked to human-specific mutations (*44*) (**Fig. 2A**). In that study, we identified “genomic hubs’’ using positional gene enrichments (*45*) (PGEs; clustered genes within chromosomes; **Fig. 2A**), and defined transcriptional compartment hubs when they shared functional enrichments, transcriptional regulation, genome structural features, disease association, and murine conservation (*44*) (**Fig. 2A, fig. S3A, B**). This approach revealed that HAR-associated genes cluster in specific genomic hubs located in interacting chromatin regions, related to metabolic function, and regulated by fasting in adipose tissue.

**Fig. 2.**
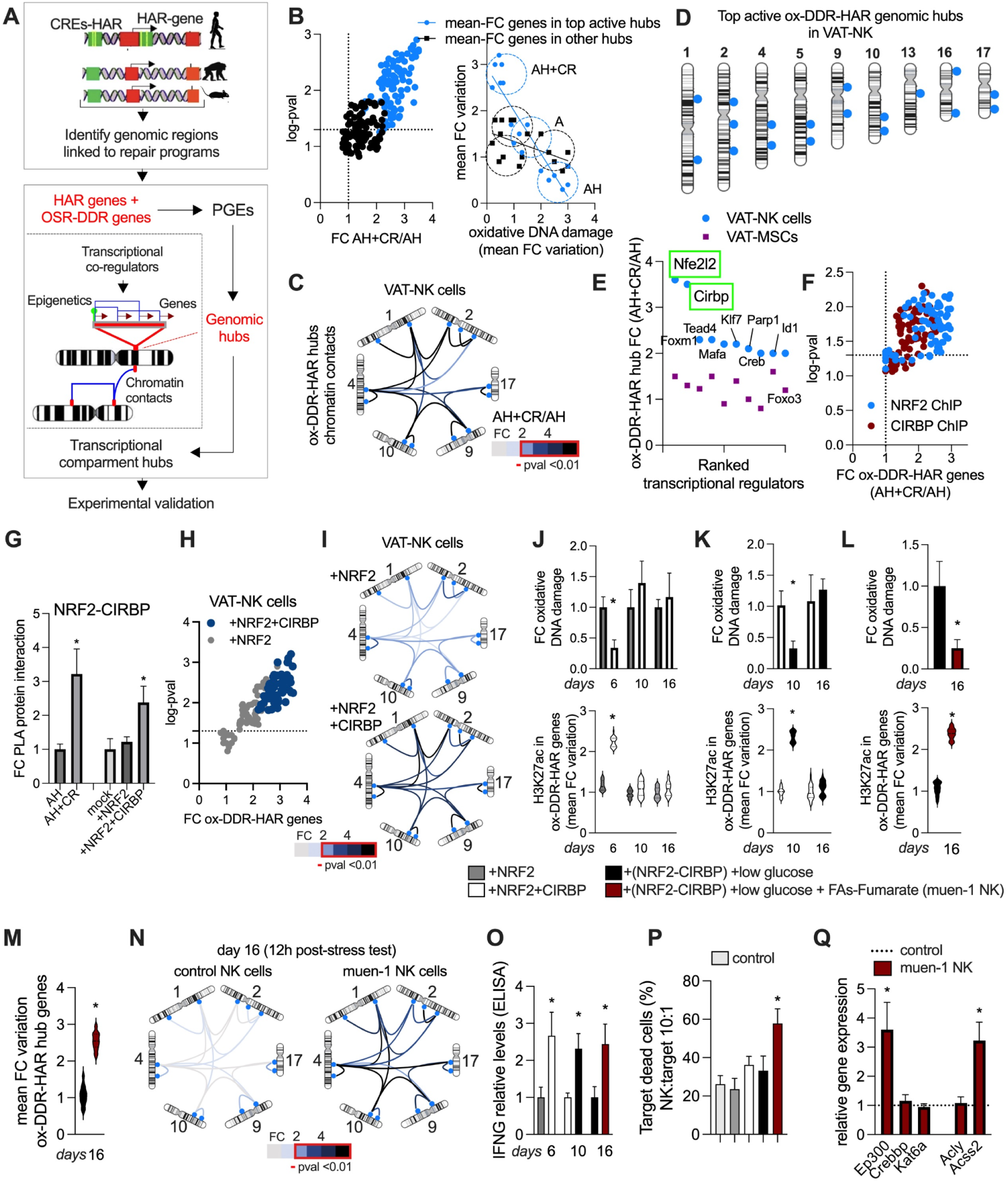
Transcriptional hubs with human-specific mutations linked to OSR and DDR programs in NK cells. (**A**) Illustration of workflow overview. (**B**) Left panel, expression levels for genomic hub genes (top hubs in blue, ranked by PGE-pval, other hubs in black) from aged-HFD(AH)+CR over the AH group. Right panel, fold-to-fold change plots from extracted VAT-NK cells from indicated mice models. Y-axes show mean fold-change variation genes in top active hubs and genes in other hubs (n=20 genes per hub group, hubs ranked by PGE p-value, number of genes in hub, and contextual activity) in each mouse per model group. X-axes show phenotypic o variations for each mouse. Lines represent linear regressions. (**C**) *in situ* ChIP-loop chromosome conformation assays for chromatin contacts between top active ox-DDR-HAR genomic hubs in extracted VAT-NK cells from mice models as in B. Targeted contact variations are represented as fold-change in AH+CR over the AH group. Contacts with FC > 2 have p-value <0.01. (**D**) Illustration of selected ox-DDR-HAR hubs (positioned blue dots in chromosomes). (**E**) oxDDR-HAR hub genes that are transcriptional regulators and regulated by CR, ranked by fold-change. (**F**) ChIP-qPCR binding fold-change for NRF2 and CIRBP to top active ox-DDR-HAR hub promoter regions. (**G**) Proximity ligation assays (PLA), for NRF2-CIRBP relative protein interactions in VAT-NK cells extracted from the indicated mice models and from VAT-NK cells transfected (12 hours) with NRF2 and CIRBP individually or combined. (**H**) In cells as in G, ChIP-qPCR binding fold-change for NRF2 and CIRBP to top active ox-DDR-HAR hub promoter regions. (**I**) In cells as in G, *in situ* ChIP-loop chromosome conformation assays for chromatin contacts between top active ox-DDR-HAR genomic hubs. Targeted contact variations are represented as fold-change in cells with combined transfection over cells with individual transfection. Contacts with FC > 2 have p-value <0.01. (**J**) Top panel, relative DNA damage variation in indicated VAT-NK cells, 12 hours post oxidative stress test (hydrogen peroxide, 250 µM) in indicated days. Lower panel, module evaluation of fold-change variation in H3K27ac levels (by ChIP) across ox-DDR-HAR hub promoter genes (n=15-20 genes) in cells as top panel. (**K**) In cells as in J and in cells with low-glucose (control 5.5 mM, low glucose is 2 mM). Top panel, relative DNA damage. Lower panel, module evaluation of fold-change variation in H3K27ac levels across ox-DDR-HAR hub promoter genes (n=15-20 genes). (**L**) In cells as in K and in cells treated with fatty acids (palmitate, 50 μM) and fumarate (50 μM). Top panel, relative DNA damage. Lower panel, module evaluation of fold-change variation in H3K27ac levels across ox-DDR-HAR hub promoter genes (n=15-20 genes). (**M**) Module evaluation of fold-change variation across ox-DDR-HAR hub genes (n=15 genes) in VAT-NK cells treated as in L. (**N**) In cells as in L, *in situ* ChIP-loop chromosome conformation assays for chromatin contacts between top active ox-DDR-HAR genomic hubs. Targeted contact variations are represented as fold-change in muen-1 cells over control cells. Contacts with FC > 2 have p-value <0.01. (**O**) In VAT-NK cells as indicated, relative IFNG levels by ELISA post-stress test. (**P**) Dead target cell cytotoxic assay. VAT-NKs from indicated models were co-incubated with VAT-MSCs from the aged-HFD model. Percentage of dead target cells stained with propidium iodide (Pi). (**Q**) Relative gene expression for epigenetic enzymes and enzymes producing acetyl-coa. Animal experiments were done with n=4-6 mice per group. Cell experiments were done with 3 independent replicates. Data show mean values per group and SEM. Cell extraction experiments were done in each mice per group and mean per groups were compared when indicated. Unpaired, two-tailed student’s t-test was used when two groups were compared, and ANOVA followed by fisher’s least significant difference (LSD) test for post hoc comparisons for multiple groups. * p-value <0.05.

Since some metabolic HAR (mHARs) regions were associated with repair pathways, we evaluate here whether mHAR genes and OSR-DDR genes colocalize in distinct genomic hubs, potentially displaying chromatin interactions and context-dependent regulation (**Fig. 2A**, **fig. S3A, B,** Table **S2**). Given our observations, we focused on evaluating this in VAT-NK cells (**fig. S3B**). By using PGE, we first identified specific genomic hubs where mHAR and OSR-DDR genes are colocalized, which was followed by experimentally testing their transcriptional activity in NK cells after CR (**Fig. 2B**, Table **S1A-D**). This revealed top active hubs, referred to as ox-DNA damage response (DDR) HAR hubs, which were selected based on gene expression, phenotype associations (oxidative DNA damage), and chromatin interactions (contacts between promoter regions using a targeted 3C approach, **Fig. 2B-D, fig. S3C-D**).

Overall, These findings suggest that functional transcriptional hubs associated with mHARs may play a key role in mediating our previously observed phenotypes in metabolic aging models and CR response. Since ox-DDR-HAR hubs are the functional units, we evaluated the transcriptional machinery driving their regulation.

### Transcriptional mediators of OSR and DDR programs in NK cells

To identify transcriptional regulators, we focused on ox-DDR-HAR hub genes that are encoding nuclear regulators. Ranking these genes by expression after CR in aged-HFD models showed top regulators in NK cells, including Nfe2l2 and Cirbp genes (**Fig. 2E**). NRF2 (gene Nfe2l2) is a regulator of oxidative stress response genes, protecting against oxidative damage (*46*). CIRBP, a cold-induced RNA-binding protein, stabilizes mRNAs under stress conditions, facilitates DNA repair mechanisms, and modulates redox signaling (*47*). NRF2 response is conserved across mammals (*48*, *49*), while the CIRBP has been shown to be fine-tuned in long-lived species, potentially enhancing DNA repair efficiency (*50*). Recent studies showed that both NRF2 and CIRBP can interact with DNA repair machinery (DRM) like PARP1, enhancing the response to DNA damage (*51–53*). In line with this, we also tested the expression of Parp1 in our models, which revealed that it mirrors the expression of Nrf2 and Cirbp, exclusively regulated in NK cells after CR (**Fig. 2E**).

To evaluate these effects in perturbation conditions, we performed RNA-interference screens in VAT-NK cells under low-glucose and oxidative stress. This revealed that perturbation of Nrf2 and Cirbp (single and in combination) significantly blunts the activity of ox-DDR-HAR hubs (**fig. S3E**). ChIP experiments confirmed a substantial increase in NRF2 binding to hub promoters in aged-HFD NK cells after CR (**Fig. 2F**). In line with previous observations on CIRBP association with transcription factors (*47*, *51*), ChIP experiments confirmed shared binding locations for CIRBP and NRF2 on hub promoters post-CR (**Fig. 2F**).

Proximity ligation assays (PLAs) on nuclear extracts confirmed increased NRF2-CIRBP interactions under CR states (**Fig. 2G**). Next, to confirm the involvement of PARP1 in these processes, we used ChIP and activity quantification. CR increased PARP1 binding to ox-DDR-HAR hubs along with increased PARP1 activity (**fig. S3F, G**). These results suggest that CR reverses the impairment of NK signatures seen in metabolic aging models via the cooperative regulation of ox-DDR-HAR hubs by NRF2 and CIRBP.

### Transcriptional and metabolic programming of CR-like NK cells

To explore whether these phenotypic adaptations can be engineered in VAT-NKs *ex vivo*, we used gain-of-function experiments via plasmid vector nucleofection in combination with key metabolic modulators (**fig. S4A**). One day post-co-transfection with Nrf2 and Cirbp vectors increased their protein interactions, enhanced ox-DDR-HAR hub gene expression, and increased chromatin contacts between key genomic hubs (**Fig. 2G-I**). To assess the duration of phenotypes, we monitored the response to oxidative stress tests and analyzed functional, transcriptional, and epigenetic changes at different time points (day 6, 10, and 16; **fig. S4A**). Transcriptional programming only provided acute resilience (6 days) to oxidative stress, DNA damage, and DNA methylation, with functional changes closely matching transcriptional activity and H3K27ac epigenetic marks in ox-DDR-HAR hubs (**Fig. 2J, fig. S4B, C**).

Recognizing the need to extend the longevity of these programmed phenotypes, we tested metabolic conditions similar to CR such as in low-glucose media. This approached prolonged time response (10 days), with resilience to oxidative stress and transcriptional-epigenetic changes (**Fig. 2K, fig. S4D, E**). In line with our *in vivo* observations, low-glucose media also increased mitochondrial activity, FAO, NAD+ levels, and fumarate accumulation (**fig. S4F, G**), which were all blunted by the FAO inhibitor etomoxir (**fig. S4F-H**), indicating the need of mitochondrial lipid oxidation for lasting transcriptional programming by metabolic factors.

Given that fumarate is known to increase NRF2 activity, while NAD+ is required for the proper DDR activity (*54–56*), we tested their supplementation. Their addition further enhanced NK phenotypic longevity (16 days), improved stress response, and enhanced transcriptional, epigenetic, and genome structural adaptations within ox-DDR-HAR hubs (referred to as multimodal engineered NK cells or muen-1 NK cells; **Fig. 2L-N, fig. S4I**). This was associated with enhanced IFNG production, cytotoxic activity, increased glutathione (GSH), NRF2 and PARP1 activity (**Fig. 2O, P, fig. S4J, K**). We observed that gamma-H2AX phosphorylation (a repair-marker linked to double strand breaks (DSBs)) was increased post-stress (12 h) but reduced substantially after 24 h, similar to apoptosis measurements (**fig. S4L, M**), suggesting improved DDR and stress response.

We next assessed the temporal gene activity of Nrf2 and Cirbp, observing acute increases in post-transfection and post-stress tests, with metabolic programming further amplifying their expression and stress-induced reactivation (**fig. S5A-C**). Notably, elevated H3K27ac levels at Nrf2 and Cirbp promoters were observed after metabolic programming and further induced post-stress tests (**fig. S5D**). In agreement, we found increased expression of the H3K27ac enzyme Ep300 and the acetyl-coa-producing enzyme Acss2 (**Fig. 2Q**).

In line with increased H3K27ac and associated enzymes, we found elevated nuclear acetyl-coa levels, which were dependent on mitochondrial FAO (**fig. S5E**). Interestingly, the Acss2 gene is also located within a top ox-DDR-HAR hub, highlighting the contextual interplay between genome structure and function (**fig. S5F**). These results suggest that metabolic mediators enhance the epigenetic memory of phenotypes in programmed cells, prolonging adaptations and enhancing NK cellular states linked to disease resilience (**fig. S5F**). To confirm these regulators are needed for CR adaptations in vivo, we tested their role in genetic deletion models.

### CR-like NK cells restore impairments from Nrf2 or Cirbp deletion

A major challenge in cell therapies is the reduced persistence of programmed phenotypes, particularly under chronic conditions where sustained stress can disrupt cell function (*57*). To evaluate this, we examined the *in vivo* response of programmed NK cells to persistent insults, using Nrf2 and Cirbp knockout (KO) mice exposed to metabolic stressors. This allowed us to assess their role as CR-mediators and their response to muen-1 NK cell allograft.

Under unchallenged conditions, KO models did not show major metabolic changes, including body weight, fat mass, GTT and ITT, or metabolic cages parameters (**fig. S6A-E**). Likewise, after 6 weeks of HFD, only minor variations in glucose tolerance were observed (**fig. S6F-K**), leading us to evaluate CR or NK cell allograft post-HFD in both control and KO strains (**fig. S7A**). KO-HFD mice did not respond to CR, exhibiting worsened metabolic parameters compared to control HFD mice after CR, with increased body weight, white adipose tissue mass, reduced VO_2_ and RER levels, altered GTT and ITT (**Fig. 3A-D, fig. S7B-E**). KO-HFD also displayed elevated plasma levels of TAGs, cholesterol, insulin, and leptin (**fig. S7F, G**). Remarkably, a single muen-1 NK cells allograft into VAT of KO-HFD mice restored physiological parameters similar to CR, unlike control NK allograft (**Fig. 3A-D, fig. S7B-G**).

**Fig. 3.**
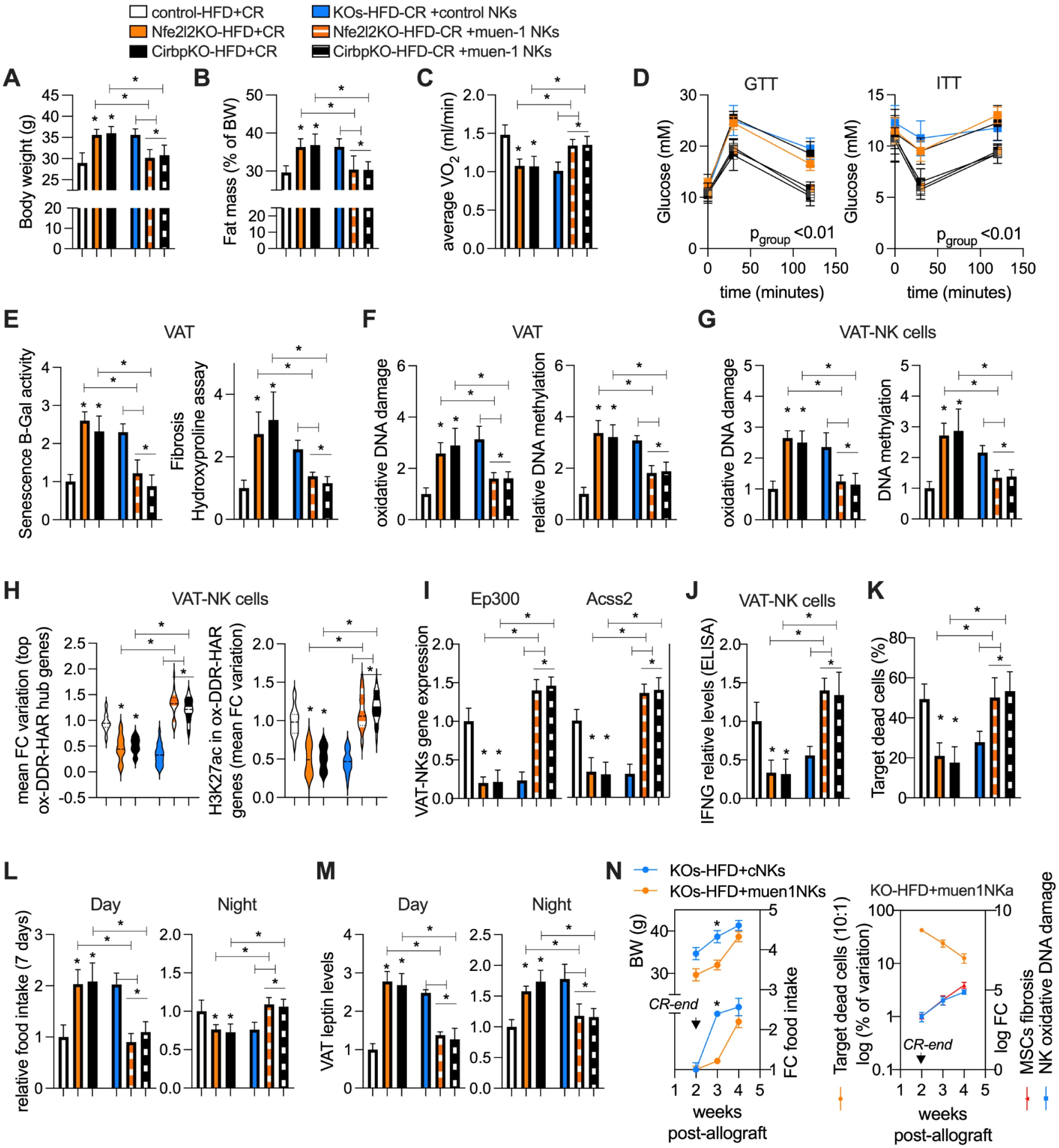
CR-like NK cells restore impairments from Nrf2 or Cirbp deletion. (**A**) Body weight (BW, grams). (**B**) Fat mass composition (% of BW). (**C**) Average VO_2_ levels (night) from 48 hours measurements (at the end of the interventions). (**D**) Left panel, glucose tolerance test (GTT). Right panel, insulin tolerance test (ITT). (**E**) Left panel, relative senescence B-gal assay activity in visceral adipose tissue (VAT). Right panel, relative hydroxyproline assay levels. (**F**) Left panel, relative oxidative DNA damage assay in VAT. Right panel, relative DNA methylation assay. (**G**) Left panel, relative oxidative DNA damage assay in VAT-NK cells. Right panel, relative DNA methylation assay. (**H**) Left panel, module evaluation of fold-change (FC) variation across ox-DDR-HAR hub genes (n=15 genes) in VAT-NK cells. Right panel, module evaluation of fold-change variation in H3K27ac levels (by ChIP) across ox-DDR-HAR hub promoter genes (n=15-20 genes). (**I**) Relative gene expression. (**J**) Relative IFNG levels by ELISA. (**K**) Dead target cell cytotoxic assay. VAT-NKs from indicated models were co-incubated with VAT-MSCs from the aged-HFD model. Percentage of dead target cells stained with propidium iodide (Pi). (**L**) Relative food intake last week of interventions. (**M**) Relative VAT leptin levels by ELISA. (**N**) Left panel, weekly follow-up of BW (left y-axis) and FC food intake (right y-axis) in mice models post-CR and reintroduced to HFD. Right panel, weekly follow-up of dead target cell assay as in K (left y-axis) and FC for VAT-MSCs fibrosis assays as in E and VAT-NK oxidative DNA damage. Animal experiments were done with n=4-6 mice per group. Cell experiments were done with 3 independent replicates. Data show mean values per group and SEM. Cell extraction experiments were done in each mice per group and mean per groups were compared when indicated. Unpaired, two-tailed student’s t-test was used when two groups were compared, and ANOVA followed by fisher’s least significant difference (LSD) test for post hoc comparisons for multiple groups. Two-way ANOVA was used to estimate significance between groups constrained by time measurements, and Tukey test for multiple comparisons * p-value <0.05.

At the molecular level, Nrf2 and Cirbp deletion impaired CR-adaptations in VAT and VAT-MSCs, leading to increased senescence, fibrosis, inflammation (**Fig. 3E, fig. S7H-M**), as well as increased VAT levels of MDA peroxidation, oxidative stress, oxidative DNA damage and DNA methylation (**Fig. 3F, fig. S7N-O**). VAT-NK cells from KO-models exhibited higher DNA damage, DNA methylation, and oxidative stress, along with reduced mitochondrial function, FAO, NAD+, and nuclear acetyl-coa levels (**Fig. 3G, fig. S8A-E**). suggesting that Nrf2 and Cirbp deletion disrupt CR-mediated genomic and metabolic plasticity. Transcriptional ox-DDR-HAR hubs adaptations seen with CR in HFD mice were absent in KO-HFD models, which showed decreased gene expression, reduced H3K27ac marks at promoters, as well as reduced expression of enzymes linked to epigenetic function (**Fig. 3H, I**). Functional assays revealed lower IFNG production, NK cytotoxic activity, glutathione and PARP1 activity, and increased DSBs markers in VAT-NKs from KO-HFD post-CR (**Fig. 3J, K, fig. S8F-H**). Notably, muen-1 NK cell allograft into VAT restored molecular adaptations in KO-HFD mice similar to CR (**Fig. 3E-K, fig. S7H-O, fig. S8A-H**). These observations indicate that Nrf2 and Cirbp are mediators of CR adaptations and that engineered muen-1 NK cells restore them, even in the presence of genetic deletions and chronic metabolic stress.

### CR-like NK cells temporarily enhance leptin sensitivity under chronic stress

Given the elevated leptin levels in our disease models and the established link between inflammation and leptin resistance (*58–60*), we hypothesized that NK-allograft weight loss is associated with leptin sensitivity. In KO-HFD models post-CR, increased food intake during the resting phase was significantly reduced by muen1 NK cell allograft, leading to night-specific time-restricted feeding patterns (**Fig. 3L**). In agreement, the elevated VAT leptin levels in KO models were reduced by muen-1 NK therapy (**Fig. 3M**). Accordingly, muen-1 NK therapy increased norepinephrine (NE) levels and thermogenesis-related module gene expression in both VAT and SAT (**fig. S8I, J**). Microdissection of the hypothalamic area revealed that leptin-responsive genes, downregulated in KO models, were increased after muen-1 NK therapy (**fig. S8K**). However, while NK therapy improved these phenotypes at early stages, some benefits diminished after 2 weeks post-allograft (**Fig. 3N, fig. S7A**).

One week post-CR, body-weight and food intake were higher in KO-HFD mice receiving control NK cells compared to mice treated with muen-1 NK cells (**Fig. 3N**). However, in VAT of HFD-KO mice treated with muen-1 NK cells, post-allograft NK cells after 3 weeks displayed reduced activity of programmed functions, leading to reduced DDR and cytotoxic activity, and increased fibrosis in MSCs (**Fig. 3N**). Overall, these results show that CR-like NK cells can mitigate metabolic impairments caused by genetic and environmental stressors, highlighting the essential role of ox-DDR-HAR hubs (**fig. S8L**). Despite the benefits of multimodal NK cell programming, the persistence of these effects during prolonged stress remains a challenge, highlighting the need for identifying additional modulators sustaining long-term cell-specific responses. To identify these modulators, we next combined experimental screens with computational analysis of CR-induced molecular signatures.

### The IL15-IL2RB pair modulated by CR links DDR to innateness signatures

To identify cytokine-receptor modulators that link DDR with NK hormesis, we used a combination of targeted experimental screens, computational interrogation of associated signatures, and high-throughput screen validation (HTS; **Fig. 4A**). First we found that CR in aged-HFD mice reduced the expression of pro-inflammatory cytokines such as Il1b, Il6, Il17, and Tnfa in both VAT-MSCs and VAT-NK cells, while upregulating anti-inflammatory cytokines including Il4 and Il10 (**Fig. 4B, C**). Notably, the interleukin-receptor pair Il15-Il2rb showed the most significant expression post-CR, with Il15 in VAT-MSCs and Il12rb in VAT-NK cells (**Fig. 4B, C**). Additionally, ELISA further confirmed increased IL2RB in muen-1 NK cells and reduced IL2RB levels in Nrf2 or Cirbp KO mice post-CR (**Fig. 4D, fig. S9A, B**). These observations suggest that CR enhances MSCs-NK association, linking CR adaptations to known drivers of NK innate function and transcriptional drivers of OSR-DDR programs.

**Fig. 4.**
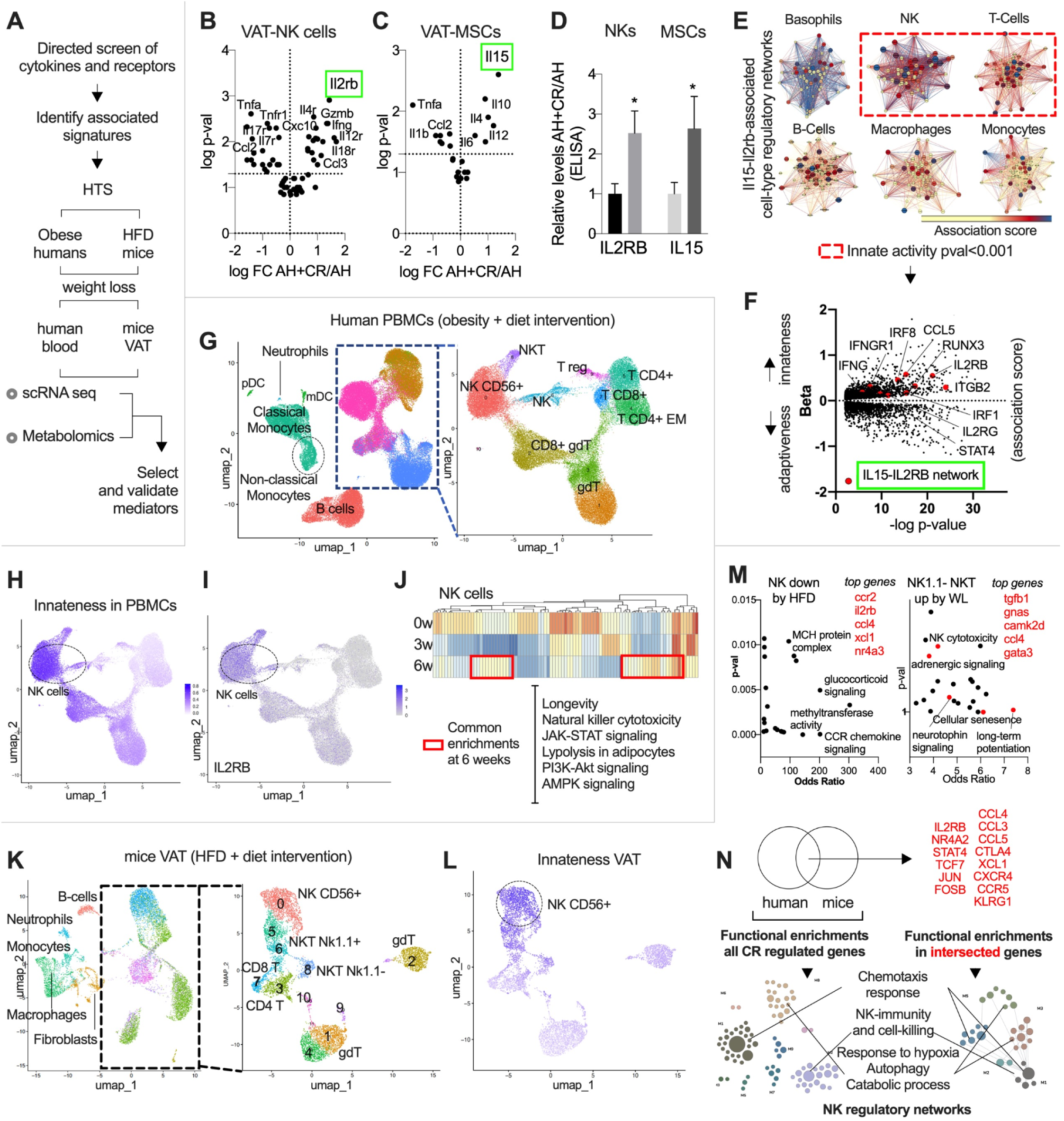
DR regulates IL15-IL2RB and innateness signatures. (**A**) Illustration of workflow overview (high-throughput screen; HTS). (**B**) Volcano-plot gene expression of cytokine-receptor pairs by qPCR in VAT-NK cells from aged-HFD+CR over aged-HFD mice (AH, n=4-6 mice). (**C**) Volcano-plot gene expression of cytokines by qPCR in VAT-MSCs cells from AH+CR over AH mice (n=4-6 mice). (**D**) Relative IL2RB and IL15 levels by ELISA in VAT-NKs and VAT-MSCs respectively. Same models as in B. (**E**) Gene regulatory networks (repository from Greene et al (*62*)) associated with IL2RB andIL15 across immune cells, and functional annotation (shaded red box). **(F**) Innateness and adaptive gradients across lymphocyte human population from scRNA-seq (Data from Gutierrez-Arcelus et al. (*63*)). Association beta score was downloaded and plotted, showing IL2RB-IL15 functional associated gene signatures from E (in red). (**G**) Umap clustering of scRNA-seq data from circulating cells from obese subjects under dietary intervention. Right panel, reclustering of lymphocyte populations. (**H**) From clusters as in G, enrichment of innateness signatures (from F) on scRNA clusters. (**I**) Il2rb gene expression in scRNA umap clusters. (**J**) Functional annotation of differentially expressed genes in blood NK cells every week during weekly dietary intervention in obese subjects. (**K**) Umap clustering of scRNA-seq data from VAT cells from HFD mice subjected to dietary weight loss intervention. Right panel, reclustering of lymphocyte populations. (**L**) From clusters as in K, enrichment of innateness signatures (from F) on scRNA clusters. (**M**) Functional biological enrichments using enrichR tools from differentially expressed genes across distinct lymphocyte populations. (**N**) Top panel, venn-diagram compiling regulated genes across both human and mice post-dietary interventions (in red upregulated genes). Lower panel, functional annotation of weight loss regulated genes in NK-specific regulatory networks (data from (*63*)), Animal experiments were done with n=4-6 mice per group. Data show mean values per group and SEM. Cell extraction experiments were done in each mice per group and mean per groups were compared. Unpaired, two-tailed student’s t-test was used when two groups were compared, and ANOVA followed by fisher’s least significant difference (LSD) test for post hoc comparisons for multiple groups. * p-value <0.05.

To further explore these findings, we evaluated public single-cell data from the tabula muris project (*61*), focusing on adipose tissue. Il15 was found mainly expressed in fibroblasts, preadipocytes, and MSCs, while Il2rb transcript was predominantly expressed in NK cells (**fig. S9C**). To identify functionally associated genes to Il15 and Il2rb, we used published Bayesian networks, which capture tissue- and cell-enriched signatures from experimental data (*62*). This revealed signatures associated with the Il15-Il2rb pair, particularly in adipose tissue and blood, with enriched innate immune response (**fig. S9D**). At the cellular level, basophils, NK cells, and T cells exhibited the highest enrichment with the Il15-Il2rb network, which were linked to chemotaxis, innate and cytotoxic function (**Fig. 4E**).

Building on a previous study that mapped innateness and adaptive gradients in lymphocytes (*63*), we further examined whether the Il15-Il2rb network aligns more closely with innate or adaptive immune responses using their published data (*63*). Innateness signatures are associated with specific programs such as cytotoxic activity and oxidative stress, all important for NK function (*63*). Consistent with this, the Il15-Il2rb regulatory network was predominantly associated with innateness signatures, particularly in NK cells where Il2rb was a top receptor correlated with innate traits (**Fig. 4F, fig. S9E-H**).

To validate these findings *in vivo*, we administered IL15 injections and observed a substantial increase in NK cells (more than in NKT, gdT, and iNKT) in VAT (more than in spleen and muscle), along with upregulated expression of key innateness genes, such as Gzmb, Ifng, and Il2rb (**fig. S10A-E**). IFNG and IL2RB levels were also confirmed by ELISA in VAT-NK cells (**fig. S10F**). Altogether, these findings highlight the link between molecular repair and innateness traits, emphasizing the significance of uncovering additional regulators via high-throughput screening to enhance immune programming. To translate these findings and identify conserved modulators across species, we analyzed immune cell responses to dietary restriction in humans and mice.

### Weight loss induces innateness signatures in blood from obese subjects

While dietary restriction (DR) and CR are both effective strategies for weight loss, they differ significantly in their approach. CR involves a strict reduction in overall caloric intake reduction, primarily focusing on reducing energy consumption. DR, on the other hand, is a more sustainable and nuanced approach that may incorporate modest caloric reduction along with changes in dietary patterns, composition, or meal timing. DR offers broader and attainable metabolic benefits, providing insights into how weight loss impacts long-term cellular and immune functions, as well as metabolic plasticity (*64*). To further investigate specific regulators that could enhance engineered NK cell function, we integrated multimodal data from dietary restriction (DR) interventions in both humans and mice (**Fig. 4A**).

Interestingly, previous studies have shown that obesity impairs the cytotoxic activity of circulating NK cells via lipid accumulation (*65*). In this study, we evaluated whether a 6-week DR intervention in obese human subjects could induce distinct transcriptional adaptations in circulating immune cells, with a focus on innateness signatures. Obese subjects undergoing this weight loss intervention showed reductions in body weight, fasting blood glucose, and HbA1c levels (**Fig. 4A, fig. S11A**). Blood samples collected at week 1, 3, and 6, underwent targeted metabolomics (from plasma) and single-cell sequencing of CD45+ immune cells. scRNA-seq analysis revealed different clusters, annotated using seurat and manual marker annotation (**Fig. 4G, fig. S11B**).

Our goal was to assess innateness profile variations in T cells, innate lymphocytes, and NK cells, while also exploring the effects on other circulating cells to provide a resource for further investigation. DR elicited specific transcriptional responses (at week 6) across various immune cells: monocytes (both classical and non-classical) showed enhanced chemokine signaling, IFNG, and FAO; B lymphocytes (memory and naive) upregulated TNFa, IFNG, and IL2 signaling; and both myeloid and plasmacytoid DCs showed increased chemotaxis, IFNG, and TNFa signaling (**Fig. 4G, Table S2**). These signature variations highlight the impact of dietary interventions on immune function, through pathways linked to chemokines, inflammation, and innateness mediators such as IFNG signaling.

To further evaluate the innate-adaptive gradients, we refined and annotated clusters associated with NK to T cells, including NKT cells, gamma-delta T cells, T regs, and both CD4+ and CD8+ T cells (**Fig. 4G**). We observed that innateness signatures were predominantly enriched in NK clusters, where the Il2rb gene is expressed, and linked to biological pathways involving IFNG response (**Fig. 4H, I, fig. S11C-E**). In addition, the innateness signature was also enriched in differentially expressed genes, with NK clusters showing strong enrichments (**fig. S11F**). Common upregulated gene-ontology programs include transcriptional processes, homeostasis, IFNG, cAMP signaling, and inflammation (**fig. S11G**). Pathway analysis in DR-induced genes within NK clusters revealed significant enrichment for longevity, AMPK signaling, NK cytotoxicity, and JAK-STAT signaling (**Fig. 4J**). Further analysis of CR-induced genes linked to cytotoxic function showed upregulation of Ccl4, Ccl5, Cd69, and Cxcr4 in NK clusters (**fig. S11G, Table S3.**). Overall, these findings revealed adaptations to dietary restriction in blood from obese subjects, such as transcriptional programs related to innate activity, particularly enriched in NK cells.

### Weight loss induces innateness signatures in VAT from HFD-mice

To assess if molecular adaptations linked to innate function observed in DR are also present in HFD-mice, we performed a study with a 6-week HFD followed by a 2-week reintroduction to a standard diet (**Fig. 4A, fig. S12A**). VAT from three groups–standard diet (SD), HFD, and HFD+DR mice–was analyzed using scRNA-seq (CD45+ cells) and metabolomics (tissue-level). As expected, the HFD+DR group showed body weight loss and reduction of adipose depots mass, reaching comparable levels to standard diet (**fig. S12A, B**). After phenotypic evaluation, VAT scRNA seq was performed and was followed by evaluation of key molecular signatures.

Single-cell data from VAT showed several clusters with marks representing different cell types such as B cells, neutrophils, monocytes, macrophages, fibroblasts, NK cells, and T cells (**Fig. 4K, fig. S12C**). Given our earlier observations, we here focused on specific signatures induced by DR such as innateness gradients, while offering other gene program changes in other cells as a dataset for exploration (**Fig. 4K**). For example, we found that differentially expressed genes during weight loss were linked to: IFNG, lysosome, and transcriptional activity in B cells; Il2/STAT5 and TNFa signaling, IFNG, and chemokine activity in DCs; lysosome, PPAR signaling, cholesterol metabolism, and chemotaxis in macrophages; TNFa and IFNG signaling, acetylation, PPARG, and TGFB signaling in monocytes; and TNFa, oxidative phosphorylation, autophagy, and transcriptional activity in neutrophils (**fig. 4K, Table S3**).

By evaluating innateness-adaptiveness gradients in specific clusters, we found enriched innateness signatures in NK and NKT clusters compared to T cells clusters, with pathways related to cytokine signaling, NK cell chemotaxis, and IFNG production (**Fig. 4K, L, fig. S12D-H**). Adaptiveness on the other hand was linked to pathways such as endoplasmic reticulum protein targeting, peptide biosynthesis, and ribosome biogenesis (**fig. S12H**). In addition, downregulated genes by HFD in innate clusters were associated with chemokine signaling, corticoid receptor, transcriptional and methyltransferase activity, with genes including Il2rb, Cxcr4, Ccr2, Ccr5, and Ccl4 (**Fig. 4M, fig. S12I, J**). In contrast, upregulated genes by HFD in these clusters were linked to insulin resistance, TLR-signaling, and TNFa signaling (**fig. S12K**), while DR upregulated genes across innate populations included Tgfb1, gnas, and camk2d, Il2rb, Igkc, Tnfaip3, and Nfatc with pathways linked to NK cytotoxicity, adrenergic signaling, senescence, and neurotrophin signaling (**Fig. 4M, fig. S12L**).

To integrate our findings, we compiled all regulated genes associated with innateness from both human and mice dietary interventions, including those downregulated by HFD or obese states and DR upregulated genes (**Fig. 4N**). We evaluated functional modules using cell-specific networks as previously indicated (*62*), identifying innate-related signatures such as chemotaxis, NK immunity, autophagy, and catabolic processes (**Fig. 4N**). Overall, our results show that DR increases innateness transcriptional programs in NK cells across both humans and mice. This regulation involves key genes associated with chemokines (Ccl3, Ccl5, Cxcr4, and Ccr5), transcription factors (Nr4a2, Stat4, and Tcf7), and the receptor gene Il2rb.

### DR-induced metabolomic signatures and multi-omic integration

To identify metabolic signatures, we analyzed targeted metabolomics in both human and mouse models. Human plasma samples, collected at week 1, week 3, and week 6 of diet restriction revealed several induced metabolites, including GABA, 4-pyridoxate, acetyl-carnitine, and allantoin (**fig. S13A, B**). These are associated with glutamate metabolism, urea cycle, glutathione, and FAO, aligning well with the increased mitochondrial activity observed in mice post-CR (**fig. S13A, B**). Similarly, plasma metabolites in mice showed that a 2-week DR post-HFD reversed variations linked to fat diet, increasing carnitines derivatives while reducing octadecanoyl-carnitine, a metabolite linked to incomplete FAO (**Fig. 5A, fig. S13C**). Weight loss was also associated with increased glycolytic metabolites pyruvate and lactate, increased TCA cycle intermediates and redox metabolites; it was further linked to reduced proline and branched chain amino acids (BCAAs; **Fig. 5A**). Metabolite enrichment analysis was associated with glutamate metabolism, FAO, glutathione metabolism, and TCA cycle (**Fig. 5A, fig. S13D**), suggesting that dietary restriction elicits similar metabolic signatures in blood across humans and mice.

**Fig. 5.**
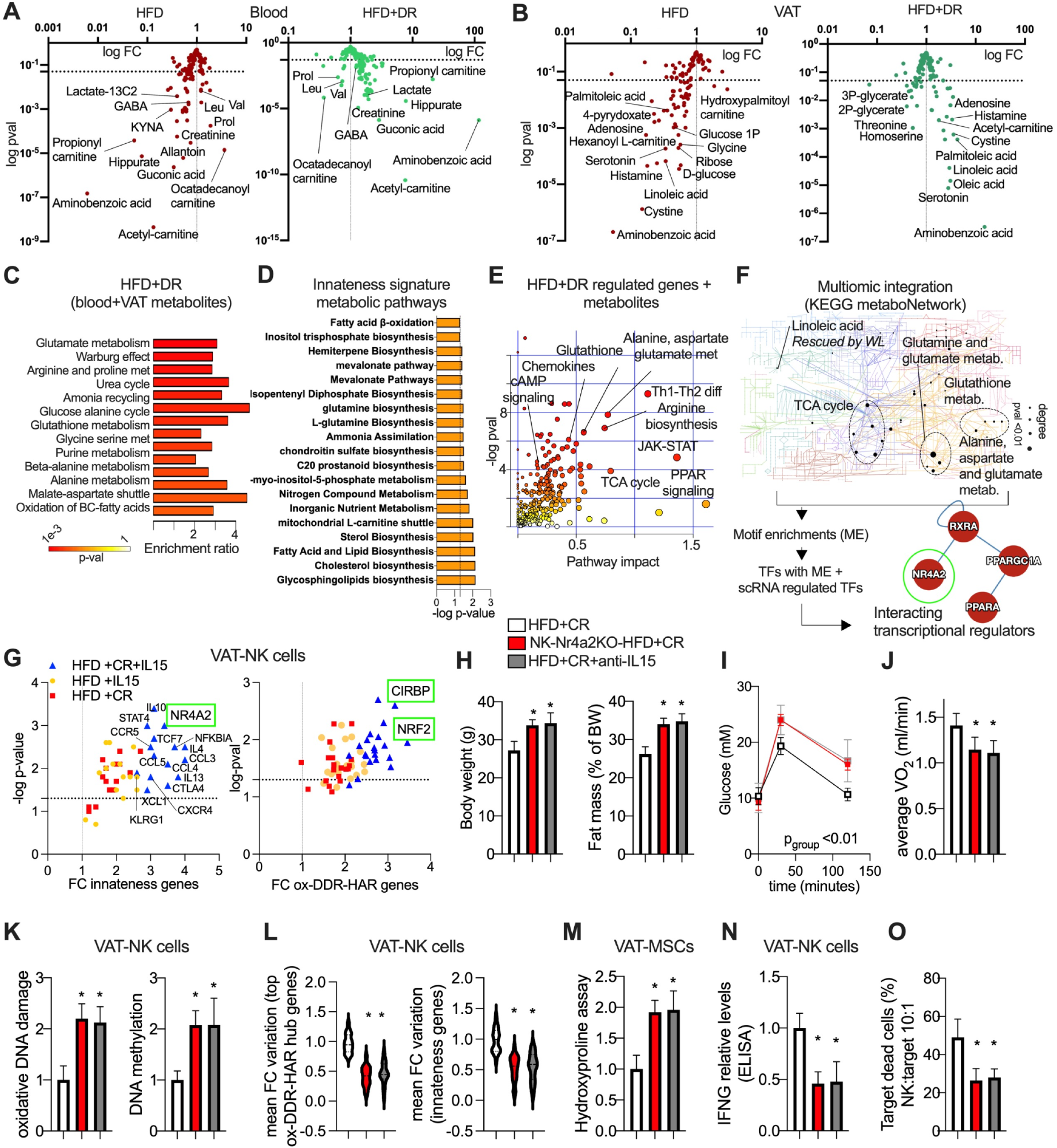
Metabolite CR-induced signatures, multi-omic integration, and validation of CR-mediators. (**A**) Volcano plots for metabolomic variation in blood plasma from mice subjected to HFD and HFD followed by dietary restriction (DR) intervention. (**B**) Volcano plots for metabolomic variation in VAT from mice as in A. (**C**) Metabolic enrichment analysis (using the metaboanalyst tool) form murine metabolites associated with DR. (**D**) Pathway enrichment for metabolic signatures (using the humancyc tool) for top-associated innateness signature genes. (**E**) Multi-omic pathway signature enrichment (using the metaboanalyst tool) for genes (Fig. 4) and metabolites associated with DR. (**F**) Top panel, multi-omic KEGG metaboNetwork analysis (using the metaboanalyst tool) for genes (Fig. 4) and metabolites associated with DR. Lower panel, motif enrichment analysis for predicted signatures (metaboanalyst), followed by inclusion of DR-induced transcription factors and evaluation of protein interactions (HuRi repository). (**G**) Left panel, volcano plot gene expression analysis for innateness genes in VAT-NK cells from mice treated with IL15 i.p. Injections alone or in combination with CR (2 weeks). Right panel, volcano plot for ox-DDR-HAR genomic hub genes in models as in left panel. (**H**) Body weight (grams; g) and fat mass percentage from indicated mice models. (**I**) Glucose tolerance test from mice as in H. (**J**) Average VO_2_ levels (night) from 48 hours measurements (at the end of the interventions). (**K**) Relative oxidative DNA damage (left) and DNA methylation (right) in VAT-NK cells from mice as in H. (**L**) Left panel, module evaluation of fold-change (FC) variation across ox-DDR-HAR hub genes (n=15 genes) in VAT-NK cells from mice as in H. Right panel, module evaluation of fold-change (FC) variation across innateness genes (n=15 genes). (**M**) Relative hydroxyproline assay activity in VAT-MSCs. (**N**) Relative IFNG levels by ELISA. (**O**) Dead target cell cytotoxic assay. VAT-NKs from indicated models were co-incubated with VAT-MSCs from the aged-HFD model. Percentage of dead target cells stained with propidium iodide (Pi). Animal experiments were done with n=4-6 mice per group. Cell experiments were done with 3 independent replicates. Data show mean values per group and SEM. Cell extraction experiments were done in each mice per group and mean per groups were compared when indicated. Unpaired, two-tailed student’s t-test was used when two groups were compared, and ANOVA followed by fisher’s least significant difference (LSD) test for post hoc comparisons for multiple groups. Two-way ANOVA was used to estimate significance between groups constrained by time measurements, and Tukey test for multiple comparisons * p-value <0.05.

In VAT, some metabolites were both reduced after HFD and increased after DR such as aminobenzoic acid, linoleic acid, cystine, serotonin, palmitoleic acid, adenosine, and acetyl-carnitine (**Fig. 5B**). Metabolite enrichment analysis showed signatures such as purine metabolism, histidine metabolism, linoleic acid metabolism, as well as glycine, serine and threonine metabolism (**fig. S13E**). Increased metabolites after DR from both blood and VAT were linked to catabolic signatures such as glutamate, glutathione, urea cycle, malate-aspartate shuttle, and FAO (**Fig. 5C**).

To complement these metabolic enrichments, we also evaluated metabolic signatures linked to DR-induced genes and innateness genes, which were linked to lipid metabolism and FAO (**Fig. 5D**). Combining DR-induced transcriptomics and metabolomics data using multi-omic pathway enrichment analysis (metaboanalyst) revealed coregulated hubs including alanine, aspartate, and glutamate metabolism, TCA cycle, glutathione, and PPAR signaling (**Fig. 5E**). KEGG-metabolic network enrichment further highlighted several key modules such as linoleic acid and the TCA (**Fig. 5F**).

To assess potential transcriptional regulators behind these signatures, we first performed transcription factor motif-enrichments. This approach revealed motifs linked to inflammatory regulators such as NFKB and STATs, along with PPAR motifs (**Fig. 5F, fig. S13F**). To the list of regulators derived from motif enrichments, we also included DR-induced transcriptional regulators from our single-cell data, and evaluated protein-protein interactions using data repositories. This approach inferred potential cooperative regulators involved in metabolic functions such as PPARA, PGC1A, RXRA, along with the DR-induced regulator, the NR4A2 transcription factor (**Fig. 5F**). NR4A2 (also known as Nurr1) is linked to neurodevelopment, inflammation, antioxidant processes, and metabolism (*66*, *67*). In addition, NR4A2 was shown to facilitate DNA repair and interact with PARP1 (*68*), in line with our observations on NRF2-CIRBP cooperative behavior. Overall, these findings underscore the multifaceted impact of DR, linking energy use, innate function, and molecular repair programs. This also highlights potential hubs mediating NK innateness including IL15-IL2RB signaling, NR4A2, and linoleic acid.

### NK-specific Nr4a2-KO and IL15-blocking antibodies impair CR-adaptations in VAT

To assess the CR role of the identified functional hubs linked to innateness in NK cells, we first focused on IL15 as a driver and NR4A2 as mediator of adaptive responses. We first evaluated the effect of IL15 i.p. injections alone or during CR in mice post-HFD. This revealed increased regulation of innateness genes and ox-DDR-HAR programs, including key genes Nr4a2, Nrf2, and Cirbp (**Fig. 5G**). To assess CR-adaptations when perturbing NR4A2 and IL15, we used NK-specific deletion of Nr4a2 and IL15 blocking antibodies. In unchallenged conditions and in HFD (6 weeks), the NK-Nr4a2-/- model did not show further variations in body weight, fat mass, and glucose tolerance (**fig. S14A-C**). However, following the HFD (6 weeks) + CR (2 weeks) intervention, both mice models showed impaired CR response, exhibiting increased body weight and fat accumulation, worsened glucose tolerance, and reduced VO_2_ (**Fig. 5H-J**). These results suggest that IL15 signaling and NR4A2 in NK cells are needed for the metabolic adaptations induced by CR.

Molecular evaluation of VAT and VAT-NKs from these models showed that both Nr4a2 deletion in NK cells and IL15-blockade led to increased oxidative DNA damage, DNA methylation, oxidative stress, MDA peroxidation, inflammation, senescence, and fibrosis (**Fig. 5K, fig. S14D-J**). CR failed to regulate ox-DDR-HAR hub gene expression and innateness genes in NK cells, including key regulators Nrf2, Cirbp, and Parp1 (**Fig. 5L**). Metabolically, these cells exhibited reduced functional parameters such as mitochondrial activity, FAO, along with reduced NAD+ levels (**fig. S14K, L**). Furthermore, these models exhibited increased fibrosis in VAT-MSCs, accompanied by reduced NK cytotoxic activity and MSCs clearing post-CR (**Fig. 5M-O**). Overall, these findings suggest that NR4A2 and IL15 signaling play an essential role in mediating CR-adaptations in VAT, linking metabolic responses, NK cytotoxic function, and molecular repair via ox-DDR-HAR hubs. The combinatorial modulation of NK phenotypes further indicates that CR-driven multimodal adjustments might be orchestrated to sustain long-term effects via epigenetic plasticity.

### Metabolic cofactors regulate epigenetic memory of resilient-states in NK cells

The relationship between 3D genome organization, gene co-regulation, cooperative signals, and transcriptional regulators shapes the epigenetic memory of cellular function, linking the duration of chromatin modifications to phenotype longevity (*31*, *39*, *69–74*). In addition, DDR mechanisms have been implicated in regulating lasting phenotypic adaptations and memory formation (*75*, *76*). Our findings show that cooperative modulators of CR in VAT-NK cells enhance transcriptional compartment regulation and epigenetic marks associated with adaptive states. In contrast, metabolic disorders and aging disrupt these processes altering the preservation of cell identities.

Given the importance of maintaining cell-state memory for homeostasis and therapeutic potential, we evaluated the duration of adaptations and epigenetic marks post-CR and after reintroducing a HFD (**fig. S15A**). In HFD mice post-CR, adaptations such as reduced body weight, VAT mass, glucose levels, VAT-TAG levels, oxidative stress, and inflammation were relatively preserved for 2 weeks after reintroduction of HFD, whereas these effects were decreased in HFD-KO-mice models (NK-Nr4a2-/-, Nrf2-/-; **fig. S15B-F**). After 4 weeks of HFD-reintroduction, most of these adaptations were lost in both control and KO mice (**fig. S15B-F**).

Molecularly, 2 weeks post-CR, both H3K27ac marks at promoter regions and gene activity within ox-DDR-HAR hub genes and innate genes remained elevated in VAT NK cells from control-HFD mice, but were significantly reduced in KO models (**Fig. 6A, fig. S15G**). After 4 weeks post-CR, these changes declined further in both groups, with the KO exhibiting a complete loss of adaptations (**Fig. 6A, fig. S15G**). Notably, oxidative-DNA damage and DNA methylation followed an inverse pattern, suggesting that DDR may play a role in sustaining epigenetic tags that support phenotype preservation (**Fig. 6B, fig. S15H, I**).

**Fig. 6.**
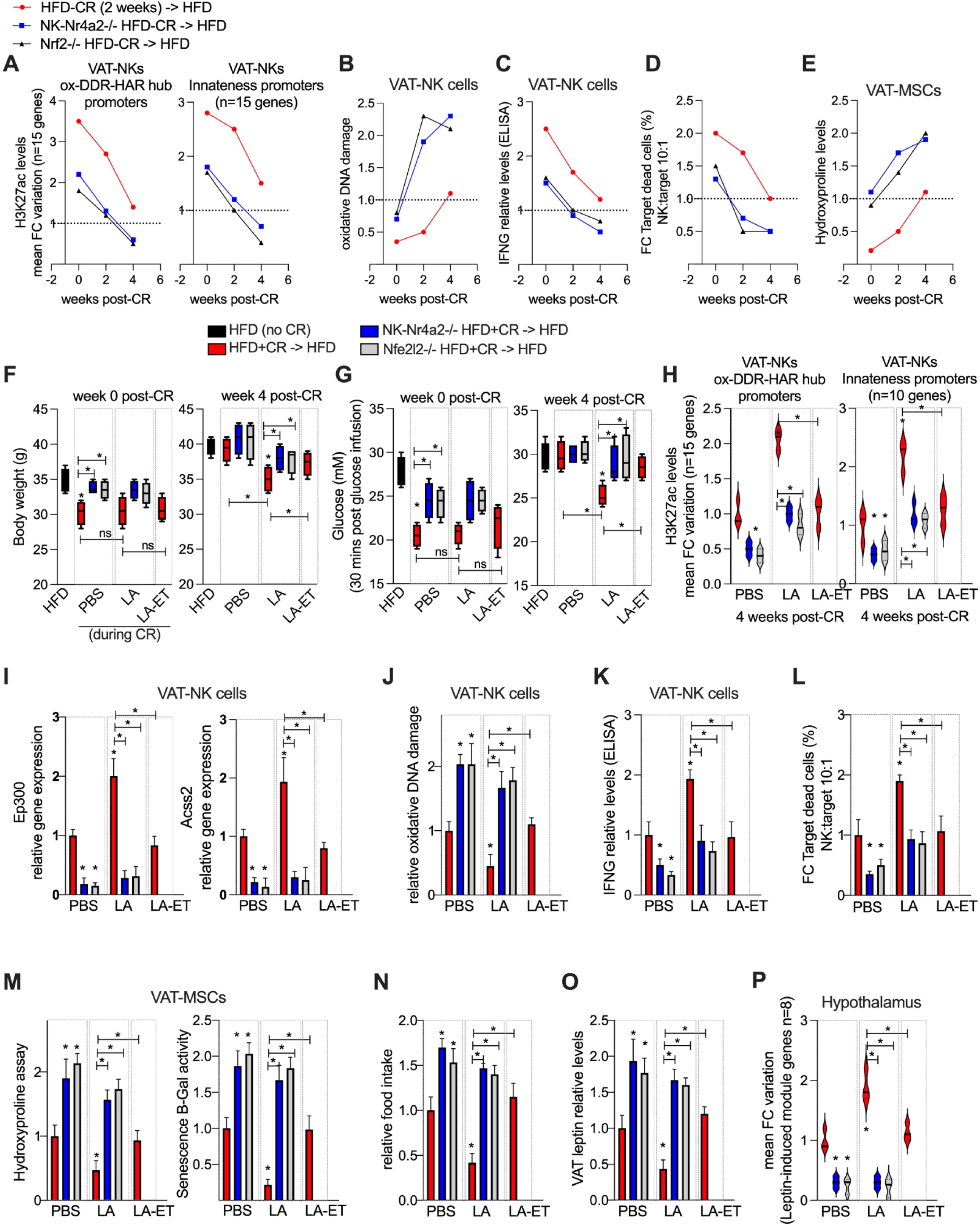
Dietary and evolutionary cofactors regulate epigenetic memory of resilience-enhancing NK cells. (**A**) Module evaluation of fold-change variation in H3K27ac levels (by ChIP) across ox-DDR-HAR hub (left panel) and innateness (right panel) promoter genes (n=15-20 genes) in VAT-NK cells from indicated mice models. (**B**) Relative oxidative DNA damage in NK cells as in A. (**C**) Relative IFNG levels by ELISA as in A. (**D**) Dead target cell cytotoxic assay. VAT-NKs from indicated models were co-incubated with VAT-MSCs from the aged-HFD model. Percentage of dead target cells stained with propidium iodide (Pi). (**E**) Relative hydroxyproline assay levels in VAT-MSCs from models as in A. (**F**) Body weight (grams; g) in indicated mice models at week 0 (right after CR intervention, left panel) and after 4 weeks of HFD reintroduction (right panel). During CR, mice were treated with PBS control, LA (10 mg/kg of body weight), and LA-etomoxir co-treatment (etomoxir 10 mg/kg). (**G**) Glucose tolerance test 30 minutes post glucose infusion, in mice as F, for week 0 (left panel) and week 4 (right panel) post-intervention. (**H**) In mice models as in F (4 weeks post-intervention), module evaluation of fold-change variation in H3K27ac levels (by ChIP) across ox-DDR-HAR hub (left panel, n=15 genes) and innateness (right panel, n=15 genes) promoter genes in VAT-NK cells. (**I**) Ep300 and Acss2 relative gene expression in VAT-NK cells as in H. (**J**) Relative oxidative DNA damage in VAT-NK cells as in H. (**K**) Relative IFNG levels by ELISA in VAT-NK cells as in H. (**L**) Dead target cell cytotoxic assay as in D. VAT-NKs from indicated models were co-incubated with VAT-MSCs from the aged-HFD model. (**M**) Left panel, relative hydroxyproline assay levels in VAT-MSCs from mice models as in H. Right panel, relative senescence B-gal assay activity. (**N**) Relative food intake last week of interventions from mice models as in H. (**O**) Relative VAT leptin levels by ELISA from mice models as in H. (**P**) Module evaluation of fold-change (FC) variation across leptin-responsive genes (n=10 genes) from mice as in H. Animal experiments were done with n=4-6 mice per group. Data show mean values per group and SEM. Cell extraction experiments were done in each mice per group and mean per groups were compared when indicated. Unpaired, two-tailed student’s t-test was used when two groups were compared, and ANOVA followed by fisher’s least significant difference (LSD) test for post hoc comparisons for multiple groups. Two-way ANOVA was used to estimate significance between groups constrained by time measurements, and Tukey test for multiple comparisons * p-value <0.05.

Cellular metabolic states influence epigenetic regulation with mitochondrial function, NAD+ levels, and TCA metabolites associated with gene activity through chromatin modifications, while reduced metabolic function is associated with CpG methylation and repressive markers (*77*, *78*). In line with this, we observed early decline in mitochondrial FAO and NAD+ levels post-CR in KO models compared to control mice (**fig. S15J, K**). Similarly, GSH and PARP1 activity in VAT-NK cells decreased post-CR, accompanied by reductions in cytotoxic activity, and increases in VAT and VAT-MSCs fibrosis and senescence (**Fig. 6C-E, fig. S15L-N**). These findings suggest that adipose tissue plasticity, FAO and related regulators sustain CR molecular processes such as molecular repair and epigenetic plasticity.

Our metabolomic analysis showed mobilization of FAs after DR, with a substantial increase in linoleic acid (LA). LA is known to modulate inflammation, immune responses, vascular function and its levels rise during fasting and physical exercise (*79–84*). Additionally, LA regulates epigenetic activity through acetyl-coa supply, enhancing memory-like function in effector T cells via mitochondrial FAO (*85*). To further investigate this, we administered LA i.p. injections during CR in control models and KOs and reintroduced them to a HFD during 4 weeks (**fig. S16A**). We tested whether LA-effects were dependent on FAO by inhibiting it with etomoxir cotreatment (**fig. S16A**). Interestingly, LA did not induce immediate changes in body weight or glucose tolerance post-CR, although these were improved by week 4 post-CR, except in KO models and in LA-etomoxir treated mice (**Fig. 6F, G**).

At week 4, LA treatment reduced VAT mass, and lowered plasma levels of TAG, cholesterol, and insulin levels (**fig. S16B-E**). In addition, LA reduced VAT TNFA levels, TAG content, and lipid peroxidation, though these modifications were absent in KO models or in the LA-etomoxir group (**fig. S16F-H**). In VAT-NK cells, H3K27ac tags in promoter regions and gene activity within ox-DDR-HAR hub genes and innate genes were still elevated 4 weeks post-CR LA-treated mice during CR, with these changes being absent in KO models and in mice receiving LA-etomoxir (**Fig. 6H, fig. S16I**). Similarly, key enzymes linked to epigenetic marks, such as Ep300 and Acss2, displayed the same variations (**Fig. 6I**). VAT-NK cells from LA treated mice showed reduced oxidative DNA damage, reduced DNA methylation, and reduced oxidative stress markers (**Fig. 6J, fig. S17A-D**). They also showed increased FAO, NAD+ levels, and nuclear acetyl-coa levels, further supporting enhanced epigenetic plasticity (**fig. S17E-G**). In agreement, these cells also exhibited elevated activity in NRF2, glutathione, and PARP1 (**fig. S17H, I**). Moreover, VAT-NK cells from LA-treated mice displayed enhanced cytotoxic activity and clearing of damaged MSCs, which was reflected in reduced VAT and VAT-MSCs fibrosis and senescence parameters (**Fig. 6K-M, fig. S17J**). Notably, these long-term LA-induced variations were absent in KO models and LA-etomoxir treated mice (**Fig. 6I-M, fig. S18A-J**).

Given these results and our previous observations on NK function and leptin sensitivity, we further evaluated associated parameters. LA-treated control mice exhibited reduced food intake, and lower plasma and VAT leptin levels (**Fig. 6N, O, fig. S17K**). This was accompanied by increased NE levels and upregulated expression of thermogenesis module genes in VAT and leptin-induced module genes in the hypothalamus (**Fig. 6P, fig. S17L, M**). Notably, the parameters associated with leptin sensitivity were blunted in KO-mice and LA-etomoxir treated mice (**Fig. 6N-P, fig. S17K-M**). These findings suggest that LA is a key mediator of CR adaptations, influencing long-term NK metabolic programming and function. This regulation maintains epigenetic memory of DDR and innate activity, contributing to overall metabolic plasticity (**fig. S17N**).

### Convergent factors enhance epigenetic memory of NK cells driving systemic resilience

We next explored whether additional coregulators (e.g. NR4A2, IlL5, and LA) could enhance cell engineering and therapeutic responses by promoting phenotype longevity. By comparing muen-1 NK cells and muen-2 NK cells (programmed with NRF2, CIRBP, NR4A2, low-glucose, IL15, and LA), we observed that, at week 6 after programming and 2 days post-oxidative stress test, only muen-2 NK cells exhibited increased gene activity and elevated epigenetic H3K27ac marks in promoter regions of ox-DDR-HAR hub genes and innateness genes (**Fig. 7A, B, fig. S18A**), compared to muen-1 NK cells that were responsive to stress-tests no longer than after 2 weeks (see **Fig. 2**).

**Fig. 7.**
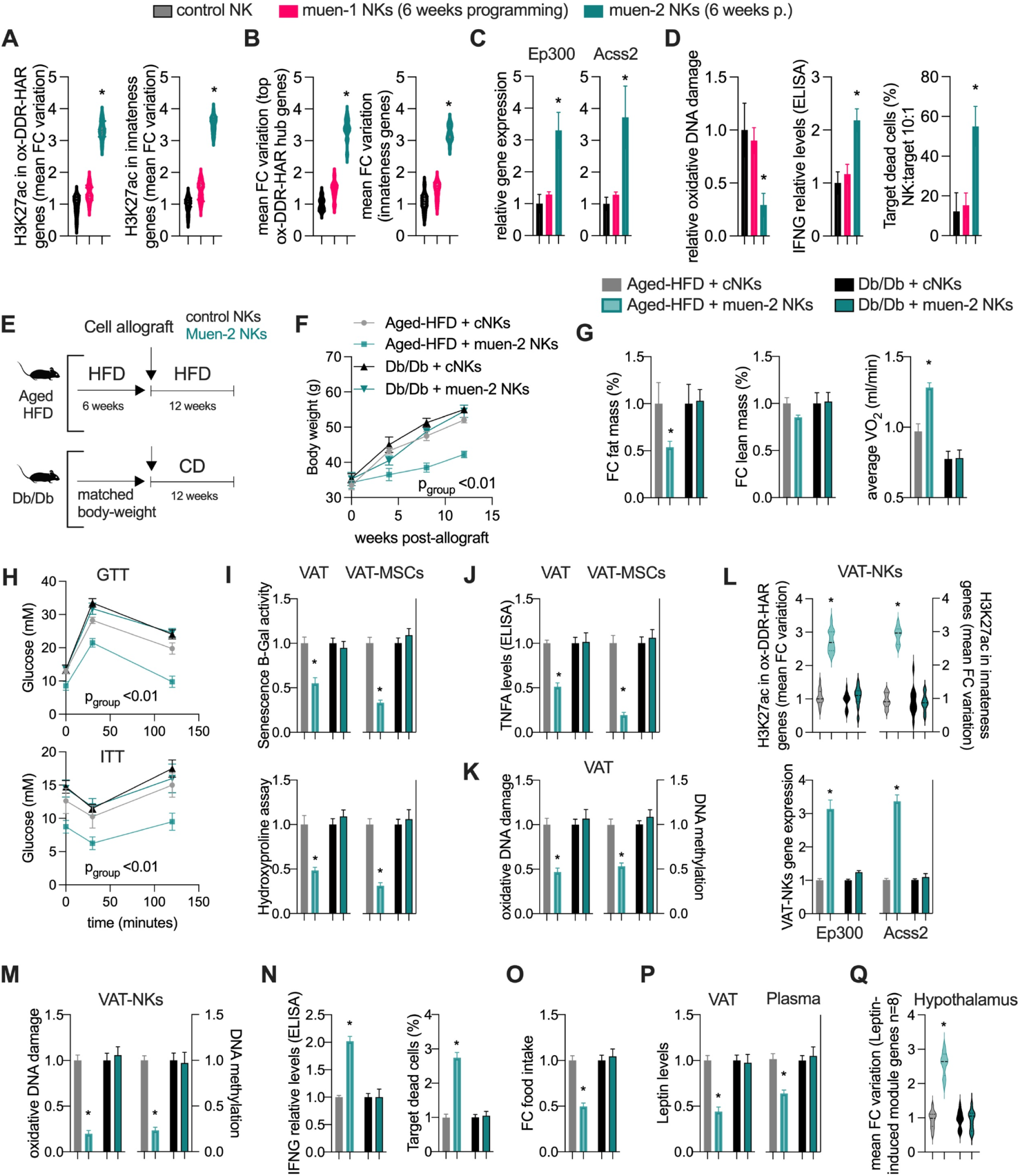
resilience-enhancing NK cells enhance long term cell therapy responses against metabolic-aging. (**A**) Module evaluation of fold-change variation in H3K27ac levels (by ChIP) across ox-DDR-HAR hub (left panel, n=15 genes) and innateness (right panel, n=15 genes) promoter genes in control, muen-1, and muen-2 VAT-NK cells at week 6, and 24 hours post-oxidative stress test (hydrogen peroxide, 250 µM). (**B**) Left panel, module evaluation of fold-change (FC) variation across ox-DDR-HAR hub genes (n=15 genes) in VAT-NK cells as in A. Right panel, module evaluation of fold-change (FC) variation across innateness genes (n=15 genes). (**C**) Relative Ep300 and Acss2 gene expression in VAT-NK cells as in A. (**D**) In cells as in A, relative oxidative DNA damage (left panel), IFNG levels by ELISA (middle panel), and dead target cell cytotoxic assay (right panel), VAT-NKs from indicated groups were co-incubated with VAT-MSCs from the aged-HFD model. Percentage of dead target cells stained with propidium iodide (Pi). (**E**) Illustration of workflow overview. (**F**) Body weight (gram; g) monitoring every 4 weeks in models as in E. (**G**) Fat mass percentage variation (left panel) and lean mass variation (middle panel) in models as in E at the end of the intervention. Average VO_2_ levels (night; right panel) from 48 hours measurements (at the end of the interventions). (**H**) Top panel, glucose tolerance test (GTT) from mice as in G at the end of the intervention. Lower panel, insulin tolerance test (ITT). (**I**) Top panel, relative senescence B-gal assay levels in VAT and VAT-MSCs from mice models as in G. Lower panel, relative hydroxyproline assay levels. (**J**) Relative TNFA levels by ELISA in VAT and VAT-MSCs from mice models as in G. (**K**) Relative oxidative DNA damage (left panel) and relative DNA methylation in VAT from mice models as in G. (**L**) Top panel, module evaluation of fold-change variation in H3K27ac levels (by ChIP) across ox-DDR-HAR hub (left panel, n=15 genes) and innateness (right panel, n=15 genes) promoter genes in VAT-NK cells from mice models as in G. Lower panel, relative Ep300 and Acss2 gene expression. (**M**) Relative oxidative DNA damage (left panel) and relative DNA methylation in VAT-NK cells from mice models as in G. (**N**) In VAT-NK cells as in G, relative IFNG levels by ELISA (left panel) and dead target cell cytotoxic assay (right panel), VAT-NKs from indicated groups were co-incubated with VAT-MSCs from the aged-HFD model. Percentage of dead target cells stained with propidium iodide (Pi). (**O**) Relative food intake last week of interventions from mice models as in G. (**P**) Relative VAT (left panel) and plasma (right panel) leptin levels by ELISA from mice models as in G. (**Q**) Module evaluation of fold-change (FC) variation across leptin-responsive genes (n=10 genes) from mice as in G. Animal experiments were done with n=4-6 mice per group. Cell experiments were done with 3 independent replicates. Data show mean values per group and SEM. Cell extraction experiments were done in each mice per group and mean per groups were compared when indicated. Unpaired, two-tailed student’s t-test was used when two groups were compared, and ANOVA followed by fisher’s least significant difference (LSD) test for post hoc comparisons for multiple groups. Two-way ANOVA was used to estimate significance between groups constrained by time measurements, and Tukey test for multiple comparisons * p-value <0.05.

To assess whether cofactor engineering leads to early transcriptional priming and subsequent enhanced gene activity, we evaluated the behavior of key regulators. This revealed a more pronounced early gene activity and H3K27ac promoter marks in key transcriptional regulators with a higher response post-stress test (**fig. S18A-C**). In agreement, muen-2 NK cells displayed increased expression of enzymes linked to H3K27ac and increased chromatin contacts between ox-DDR-HAR hubs post-stress test (**Fig. 7C, fig. S18D**). We also observed that NR4A2 binding to promoters of ox-DDR-HAR and innateness genes was increased substantially post-stress (**fig. S18E**), suggesting transcriptional memory through cooperative coregulation with early priming of key mediators.

Molecularly, these changes were associated with reduced oxidative DNA damage and methylation, reduced oxidative stress and inflammation, and increased mitochondrial activity, FAO, NAD+, and nuclear acetyl-coa levels (**Fig. 7D, fig. S18F-K**). Functionally, this led to increased NRF2, glutathione, and PARP activity, reduced phospho-H2AX and apoptosis, along with enhanced cytotoxic activity (**Fig. 7D, fig. S18L-O)**. Overall, these results suggest that transcriptional and metabolic engineering of cooperative signals regulating DDR and innate activity can significantly enhance epigenetic memory and phenotype longevity in NK cells.

To further assess the long-term therapeutic potential and association with leptin signaling, we conducted a single allograft of muen-2 NK cells in aged-HFD mice and leptin receptor-deficient db/db mice (**Fig. 7D**). The cell allograft (control and muen-2 NK cells) was performed into VAT at week 6 of an 18-week HFD protocol in aged mice and body-weight matched Db/Db mice (10-13 weeks of age), and we monitored body weight gain every month and examined molecular adaptations after 12 weeks (**Fig. 7D**). Muen-2 NK cell therapy significantly improved metabolic outcomes in the aged-HFD model, not in the db/db mice, with changes including reduced body weight gain, decreased white fat mass, increased VO_2_ and reduced RER, as well as improved GTT and ITT (**Fig. 7E-I, fig. S19A, B**).

In addition, aged-HFD mice treated with muen-2 NK showed reductions in plasma levels of TAGs, cholesterol, and insulin, reductions in VAT levels of TAG, MDA, and ROS (**fig. S19C-F**). This was accompanied with reductions in VAT and VAT-MSCs levels of senescence, fibrosis, and inflammation (**Fig. 7J, H**). These benefits, on the other hand, were not observed in the db/db mice. Muen-2 NK benefits were also associated with reduced VAT oxidative DNA damage and methylation (**Fig. 7I**).

VAT-NK cells from aged-HFD mice treated with muen-2 NKs displayed increased H3K27ac marks in promoters with increased ox-DDR-HAR and innateness genes, along with elevated expression levels of key enzymes Ep300 and Acss2 (**Fig. 7J, fig. S19G**). Again, these changes were absent in db/db mice. Additionally, VAT-NKs cells from muen-2 NK treated mice showed reduced oxidative DNA damage, DNA methylation, oxidative stress markers, while showing increased mitochondrial function, FAO, NAD+, and nuclear acetyl-coa levels (**Fig. 7K, fig. S19H-J**). These adaptations were again absent in db/db mice models. Functionally, VAT-NKs from muen-2 treated aged-HFD mice exhibited elevated cytotoxic activity, NRF2, GSH, and PARP1 activity, with reduced phospho-H2AX (**Fig. 7L, fig. S19K, L**). In addition, evaluation of other organs, such as in liver, hypothalamus, heart, and skeletal muscle, revealed significant reductions in both inflammation and fibrosis in muen-2 NK treated aged-HFD mice (**fig. S20A, B**).

Since our results indicated that the enhanced longevity of programmed NK function and organismal homeostasis are influenced by leptin signaling and sensitivity, we further evaluated associated parameters. This revealed reduced food intake along with decreased plasma and VAT leptin levels in muen-2 NK treated mice, with no changes in db/db mice (**Fig. 7M, N**). This was associated with increased NE levels, elevated expression of thermogenesis module genes in white adipose tissue, as well as elevated expression of leptin-induced module genes in hypothalamus (**fig. S20C, D**). Altogether, these findings show that multimodal modulators enhance programming and memory-like functions of resilience-promoting NK cells. This also suggests that optimizing epigenetic memory of specific NK cell function holds promise for cell therapy approaches targeting chronic conditions such as obesity, metabolic disease, and aging (**fig. S20E**).

## Discussion

Metabolic plasticity is critical for molecular and evolutionary adaptation, and its decline, seen in metabolic disease, increases the risk of comorbidities accelerating aging. Conversely, energy-conservation states such as CR enhance metabolic plasticity and overall health. This plasticity depends on the mobilization of resources for molecular adaptations, where adipose tissue plays a central role in sustaining physiological adaptations. Aging and metabolism are tightly interconnected; while aging can be driven by damage accumulation, metabolic processes provide energy and resources for molecular repair (*40*).

In line with this, genome maintenance appears to be more robust in longer-lived species (*17–19*) while CR enhances these repair mechanisms (*11–14*). Building on previous studies (*15*, *16*), we observe an inverse association between genome instability and metabolic traits such as fat mass composition and fasting endurance score in mammals. This suggests that these traits are not only associated but also fine-tuned for phenotypic adaptations, which led us to investigate the molecular links between evolution and metabolic plasticity, as well as identifying functional hubs linked to resilient phenotypes.

Convergent signals are responsible for persistent memory of cellular function. We show this by engineering NK cells with sustained therapeutic function under chronic metabolic stress. While CR turns on molecular repair programs in many different cells, maintenance of these adaptations requires the integration of cell specific signals supporting epigenetics memory. Our work identifies convergence of the OSR-DDR with innate immune signals through evolutionary-tuned genomic hubs, enabling durable functional specialization.

### Operational principles linking energy conservation to cellular memory

Our findings show that there are three integrated mechanisms of action of CR in VAT-NK cells. First, ox-DDR-HAR hubs serve as functional scaffolds for the coordination of stress response and cytotoxic programs by cooperative regulators (NRF2, CIRBP, NR4A2). Second, intermediates from FAO (especially linoleic acid conversion to acetyl coenzyme A (AcCoA)) produce cofactors that maintain H3K27ac epigenetic marks at these hubs. Third, IL15-IL2RB signaling provides a link between metabolic state and cell-specific function, coupling repair cycles to innate activity. This convergence allows for epigenetic memory, where NK cells retain functional adaptations for weeks after initial programming even in the presence of metabolic challenges.

Conservation of these mechanisms across different species, from comparative trait analysis in mammals, to murine models and human dietary intervention, indicate fundamental design principles. Body fat composition is inversely related to somatic mutation rates and DNA methylation in mammals, relationships that are seen mechanistically in the way VAT coordinates systemic responses. When adipose tissue plasticity is sustained (CR, or linoleic acid supplementation, muen-NK allograft) NK cells preserve repair capacity and cytotoxic function. When this is disturbed (by HFD, genetic deletion of NRF2/CIRBP/NR4A2 or by interference in leptin signaling), molecular damage accumulates and cells lose molecular identity.

### Evolutionary tuning of metabolic-immune integration

The activity of ox-DDR-HAR hubs in NK cells is suggestive of evolutionary optimization of metabolic-immune crosstalk. HAR regions with a neural developmental connection have been described before (*5*, *6*); however, our identification of metabolic HARs connecting repair programs to innate function shows additional axes of human-specific adaptation. The cooperative regulation by NRF2 (conserved stress response), CIRBP (fine-tuned in long-lived species) and NR4A2 (linking metabolism and inflammation) are examples of how layered evolutionary modifications can tune adaptability and systemic resilience. Body fat composition in hominids may have been selected not only for energy storage, when food is scarce, but because it helped coordinate tissue-resident immune function by lipid-derived signals such as linoleic acid.

### Implications for memory engineering in adaptive systems

A major obstacle to cell therapy methods is the persistence of phenotypes in chronic pathologic conditions. Our muen-2 NK cells retain therapeutic function for several weeks post-transfer, rescuing metabolic parameters in disease models through sustained clearance of damaged MSCs and leptin sensitivity. This durability relies on epigenetic memory as H3K27ac marks at ox-DDR-HAR hubs are maintained even after the expression of transgenes ceased, allowing for the reactivation of protective programs by stress. The reliance on leptin signaling identifies system level needs as engineered cells are not able to overcome complete disruption of nutrient sensing (db/db mice), but can re-establish leptin sensitivity when core machinery remains intact.

The multimodal programming strategy has practical implications. Transcriptional programming alone has only acute benefits while addition of metabolic conditioning and cell type signals prolonged phenotype persistence. This progressive extension reflects a more stable epigenetic encoding, ranging from transcription factor binding, metabolic cofactor availability and integration with developmental identity programs. This is in line with recent studies showing cell-specific responses despite similar cytokine and receptor activity (*86*) and with findings on the influence of 3D genome organization on persistent epigenetic adaptations (*72*).

More broadly, our findings reveal several principles that can be generalized for engineering persistent memory in adaptive systems: (i) functional memory emerges from signal convergence, not signal strength: multiple inputs convert to a shared substrate for stable encoding of adaptive states. Computationally, using parallel multimodal activation into a common, grounding yet different substrate (from inputs) instead of threshold-crossing in one input channel might offer better encoding. (ii) Substrate persistence decoupled from its writing signals enable memory of functional states without continuous input. (iii) Multi-timescale signal integration enables plasticity and stability: adaptive responses occur at different time-scales filtering transient noise for substrate consolidation. Temporal hierarchies (from fast patterns, integration over longer windows, and consolidation over recurring features) could help achieve adaptive learning and memory. Altogether, this suggests that robust long-term memory requires design rules that are identified in cell systems yet are generalizable, opening new avenues for their implementation through heterogeneous memory systems with write and decay rates operating in parallel.

### Limitations and future directions

Several aspects need further investigation. First, our focus on VAT-NK cells leaves open several questions for example whether similar principles apply to other tissue-resident immune populations or whether different genomic hubs are operational in different contexts. Second, although we show that linoleic acid prolongs epigenetic memory, the entire metabolic intermediate network that controls associated chromatin modifications is yet to be described. Third, the data on human dietary restriction, although consistent with mouse data, is from a 6-week intervention in a limited number of people. Longer-term studies with larger numbers are needed to validate our findings. Fourth, knockout models tested the need of individual factors, but combinatorial deletion would be a better way of defining redundancy and epistasis in a systematic way.

Finally, although muen-2 NK cells exhibit phenotype persistence in mice, showing similar durability in human cells and safety in larger animal models will be needed. We show that engineered NK cells can be a cell therapy platform, yet clinical validation in metabolic disease patients is needed. The therapeutic effects were found to be leptin dependent, which implies that patient stratification by leptin receptor status might be important. In addition, testing other models (obesity-associated cancer, autoimmune diseases) would help demonstrate how general these approaches are.

### Concluding perspective

Our findings define the molecular capability of convergent signals (transcriptional regulators that are sensitive to changes in energy state, metabolic cofactors from tissue remodeling, and cell-specific differentiation cues) to integrate and encode long-term memory of adaptive cellular states. This offers both mechanistic insights into how CR effects can be sustained for longer time than the restriction period and an engineering framework for cell therapies requiring preserved function in challenging microenvironments. More generally, the idea that repair cycles (driven by energy-conservation) can be coupled to epigenetic memory via genomic hubs is a design principle with applications beyond NK cells. This can have implications for approaches aiming at maintaining cellular identity in aging, regenerative medicine and synthetic biology. The evolutionary coupling of metabolic plasticity with immune surveillance raises the possibility that therapies aimed at this axis could exploit deeply conserved mechanisms for organismal resilience.

## Acknowledgments

We thank all the members from the various labs who have contributed to this project. We also thank Amy Grayson for her help reviewing and editing the manuscript.

## Funding

LZA and MK are supported by the Novo Nordisk foundation, Novo Nordisk research center in Seattle and Boston. TF and AM are supported by Ramon y Cajal grants by the Spanish state research agency, and the “Severo Ochoa ’’ programme for Centres of Excellence in R&D (SEV-2017-0723). KV was supported by Novo Nordisk.

## Author contributions

Project design and Conceptualization: LZA, LL, TF, MK. Computational Methodology: LZA with feedback from LL, NS, HP, MK, TF, KG. Experimental design and validation: LZA, HP, CH, KG, LL, TF, KG. Supervision of the work: LZA, LL, MK, TF. Funding acquisition: LZA, LL, MK, TF. Writing of original draft: LZA. Writing & editing: LZA, TF, LL, MK. All authors reviewed the manuscript.

## Competing interests

Authors declare non competing interests.

## Data and materials availability

Data is available in the supplementary materials.

## Supplementary Materials

Figures S1-20

Tables S1-S12

Materials and methods

## Notes

### Competing Interest Statement

The authors have declared no competing interest.

## References and Notes

1. C. R. White, L. A. Alton, C. L. Bywater, E. J. Lombardi, D. J. Marshall, Metabolic scaling is the product of life-history optimization. Science 377, 834–839 (2022).

2. C. R. White, D. J. Marshall, L. A. Alton, P. A. Arnold, J. E. Beaman, C. L. Bywater, C. Condon, T. S. Crispin, A. Janetzki, E. Pirtle, H. S. Winwood-Smith, M. J. Angilletta Jr, S. F. Chenoweth, C. E. Franklin, L. G. Halsey, M. R. Kearney, S. J. Portugal, D. Ortiz-Barrientos, The origin and maintenance of metabolic allometry in animals. Nat Ecol Evol 3, 598–603 (2019).

3. J. Guo, X. Huang, L. Dou, M. Yan, T. Shen, W. Tang, J. Li, Aging and aging-related diseases: from molecular mechanisms to interventions and treatments. Signal Transduct Target Ther 7, 391 (2022).

4. V. D. Longo, M. P. Mattson, Fasting: molecular mechanisms and clinical applications. Cell Metab. 19, 181–192 (2014).

5. K. S. Pollard, S. R. Salama, N. Lambert, M.-A. Lambot, S. Coppens, J. S. Pedersen, S. Katzman, B. King, C. Onodera, A. Siepel, A. D. Kern, C. Dehay, H. Igel, M. Ares Jr, P. Vanderhaeghen, D. Haussler, An RNA gene expressed during cortical development evolved rapidly in humans. Nature 443, 167–172 (2006).

6. J. A. Capra, G. D. Erwin, G. McKinsey, J. L. R. Rubenstein, K. S. Pollard, Many human accelerated regions are developmental enhancers. Philos. Trans. R. Soc. Lond. B Biol. Sci. 368, 20130025 (2013).

7. D. E. Lieberman, D. M. Bramble, The evolution of marathon running : capabilities in humans. Sports Med. 37, 288–290 (2007).

8. H. Pontzer, M. H. Brown, D. A. Raichlen, H. Dunsworth, B. Hare, K. Walker, A. Luke, L. R. Dugas, R. Durazo-Arvizu, D. Schoeller, J. Plange-Rhule, P. Bovet, T. E. Forrester, E. V. Lambert, M. E. Thompson, R. W. Shumaker, S. R. Ross, Metabolic acceleration and the evolution of human brain size and life history. Nature 533, 390–392 (2016).

9. S. L. Lindstedt, M. S. Boyce, Seasonality, Fasting Endurance, and Body Size in Mammals. [Preprint] (1985). 10.1086/284385.

10. C. C. Lindsey, BODY SIZES OF POIKILOTHERM VERTEBRATES AT DIFFERENT LATITUDES. Evolution 20, 456–465 (1966).

11. A. R. Heydari, A. Unnikrishnan, L. V. Lucente, A. Richardson, Caloric restriction and genomic stability. Nucleic Acids Res. 35, 7485–7496 (2007).

12. W. P. Vermeij, M. E. T. Dollé, E. Reiling, D. Jaarsma, C. Payan-Gomez, C. R. Bombardieri, H. Wu, A. J. M. Roks, S. M. Botter, B. C. van der Eerden, S. A. Youssef, R. V. Kuiper, B. Nagarajah, C. T. van Oostrom, R. M. C. Brandt, S. Barnhoorn, S. Imholz, J. L. A. Pennings, A. de Bruin, Á. Gyenis, J. Pothof, J. Vijg, H. van Steeg, J. H. J. Hoeijmakers, Restricted diet delays accelerated ageing and genomic stress in DNA-repair-deficient mice. Nature 537, 427–431 (2016).

13. A. V. Everitt, S. I. S. Rattan, D. G. Couteur, R. de Cabo, Calorie Restriction, Aging and Longevity (Springer Science & Business Media, 2010).

14. G. M. Lim, N. Maharajan, G.-W. Cho, How calorie restriction slows aging: an epigenetic perspective. J. Mol. Med., doi: 10.1007/s00109-024-02430-y (2024).

15. A. Cagan, A. Baez-Ortega, N. Brzozowska, F. Abascal, T. H. H. Coorens, M. A. Sanders, A. R. J. Lawson, L. M. R. Harvey, S. Bhosle, D. Jones, R. E. Alcantara, T. M. Butler, Y. Hooks, K. Roberts, E. Anderson, S. Lunn, E. Flach, S. Spiro, I. Januszczak, E. Wrigglesworth, H. Jenkins, T. Dallas, N. Masters, M. W. Perkins, R. Deaville, M. Druce, R. Bogeska, M. D. Milsom, B. Neumann, F. Gorman, F. Constantino-Casas, L. Peachey, D. Bochynska, E. S. J. Smith, M. Gerstung, P. J. Campbell, E. P. Murchison, M. R. Stratton, I. Martincorena, Somatic mutation rates scale with lifespan across mammals. Nature 604, 517–524 (2022).

16. S. J. C. Crofts, E. Latorre-Crespo, T. Chandra, DNA methylation rates scale with maximum lifespan across mammals. Nat Aging 4, 27–32 (2024).

17. S. L. MacRae, M. M. Croken, R. B. Calder, A. Aliper, B. Milholland, R. R. White, A. Zhavoronkov, V. N. Gladyshev, A. Seluanov, V. Gorbunova, Z. D. Zhang, J. Vijg, DNA repair in species with extreme lifespan differences. Aging 7, 1171–1184 (2015).

18. D. Toren, A. Kulaga, M. Jethva, E. Rubin, A. V. Snezhkina, A. V. Kudryavtseva, D. Nowicki, R. Tacutu, A. A. Moskalev, V. E. Fraifeld, Gray whale transcriptome reveals longevity adaptations associated with DNA repair and ubiquitination. Aging Cell 19, e13158 (2020).

19. L. Zhang, X. Dong, X. Tian, M. Lee, J. Ablaeva, D. Firsanov, S.-G. Lee, A. Y. Maslov, V. N. Gladyshev, A. Seluanov, V. Gorbunova, J. Vijg, Maintenance of genome sequence integrity in long- and short-lived rodent species. Sci Adv 7, eabj3284 (2021).

20. E. H. Finn, T. Misteli, Molecular basis and biological function of variability in spatial genome organization. Science 365 (2019).

21. T. Misteli, The Self-Organizing Genome: Principles of Genome Architecture and Function. Cell 183, 28–45 (2020).

22. C. H. Eskiw, N. F. Cope, I. Clay, S. Schoenfelder, T. Nagano, P. Fraser, Transcription factories and nuclear organization of the genome. Cold Spring Harb. Symp. Quant. Biol. 75, 501–506 (2010).

23. D. Hnisz, K. Shrinivas, R. A. Young, A. K. Chakraborty, P. A. Sharp, A Phase Separation Model for Transcriptional Control. Cell 169, 13–23 (2017).

24. B. R. Sabari, A. Dall’Agnese, A. Boija, I. A. Klein, E. L. Coffey, K. Shrinivas, B. J. Abraham, N. M. Hannett, A. V. Zamudio, J. C. Manteiga, C. H. Li, Y. E. Guo, D. S. Day, J. Schuijers, E. Vasile, S. Malik, D. Hnisz, T. I. Lee, I. I. Cisse, R. G. Roeder, P. A. Sharp, A. K. Chakraborty, R. A. Young, Coactivator condensation at super-enhancers links phase separation and gene control. Science 361 (2018).

25. A. R. Strom, A. V. Emelyanov, M. Mir, D. V. Fyodorov, X. Darzacq, G. H. Karpen, Phase separation drives heterochromatin domain formation. Nature 547, 241–245 (2017).

26. J. A. Riback, L. Zhu, M. C. Ferrolino, M. Tolbert, D. M. Mitrea, D. W. Sanders, M.-T. Wei, R. W. Kriwacki, C. P. Brangwynne, Composition-dependent thermodynamics of intracellular phase separation. Nature 581, 209–214 (2020).

27. A. Klosin, F. Oltsch, T. Harmon, A. Honigmann, F. Jülicher, A. A. Hyman, C. Zechner, Phase separation provides a mechanism to reduce noise in cells. Science 367, 464–468 (2020).

28. T. Jenuwein, C. D. Allis, Translating the histone code. Science 293, 1074–1080 (2001).

29. A. J. S. Klar, Propagating epigenetic states through meiosis: where Mendel’s gene is more than a DNA moiety. [Preprint] (1998). 10.1016/s0168-9525(98)01535-2.

30. J. C. Rice, C. D. Allis, Histone methylation versus histone acetylation: new insights into epigenetic regulation. Curr. Opin. Cell Biol. 13, 263–273 (2001).

31. A. D’Urso, J. H. Brickner, Mechanisms of epigenetic memory. Trends Genet. 30, 230–236 (2014).

32. W. H. Light, J. Freaney, V. Sood, A. Thompson, A. D’Urso, C. M. Horvath, J. H. Brickner, A conserved role for human Nup98 in altering chromatin structure and promoting epigenetic transcriptional memory. PLoS Biol. 11, e1001524 (2013).

33. B. Sump, D. G. Brickner, A. D’Urso, S. H. Kim, J. H. Brickner, Mitotically heritable, RNA polymerase II-independent H3K4 dimethylation stimulates transcriptional memory. Elife 11 (2022).

34. W. Siwek, S. S. H. Tehrani, J. F. Mata, L. E. T. Jansen, Activation of Clustered IFNγ Target Genes Drives Cohesin-Controlled Transcriptional Memory. Mol. Cell 80, 396–409.e6 (2020).

35. V. Sood, J. H. Brickner, Genetic and Epigenetic Strategies Potentiate Gal4 Activation to Enhance Fitness in Recently Diverged Yeast Species. Curr. Biol. 27, 3591–3602.e3 (2017).

36. P. Pascual-Garcia, B. Debo, J. R. Aleman, J. A. Talamas, Y. Lan, N. H. Nguyen, K. J. Won, M. Capelson, Metazoan Nuclear Pores Provide a Scaffold for Poised Genes and Mediate Induced Enhancer-Promoter Contacts. Mol. Cell 66, 63–76.e6 (2017).

37. Z. D. Smith, A. Meissner, DNA methylation: roles in mammalian development. Nat. Rev. Genet. 14, 204–220 (2013).

38. A. Barzilai, K.-I. Yamamoto, DNA damage responses to oxidative stress. DNA Repair 3, 1109–1115 (2004).

39. R. Scully, A. Panday, R. Elango, N. A. Willis, DNA double-strand break repair-pathway choice in somatic mammalian cells. Nat. Rev. Mol. Cell Biol. 20, 698–714 (2019).

40. C. López-Otín, M. A. Blasco, L. Partridge, M. Serrano, G. Kroemer, Hallmarks of aging: An expanding universe. Cell 186, 243–278 (2023).

41. D. Muñoz-Espín, M. Serrano, Cellular senescence: from physiology to pathology. Nat. Rev. Mol. Cell Biol. 15, 482–496 (2014).

42. B. G. Childs, M. Durik, D. J. Baker, J. M. van Deursen, Cellular senescence in aging and age-related disease: from mechanisms to therapy. Nat. Med. 21, 1424–1435 (2015).

43. E. Verdin, NAD^+^ in aging, metabolism, and neurodegeneration. Science 350, 1208–1213 (2015).

44. L. Z. Agudelo, R. Tuyeras, C. Llinares, A. Morcuende, Y. Park, N. Sun, S. Linna-Kuosmanen, N. Atabaki-Pasdar, L.-L. Ho, K. Galani, P. W. Franks, B. Kutlu, K. Grove, T. Femenia, M. Kellis, Metabolic resilience is encoded in genome plasticity, bioRxiv (2021)p. 2021.06.25.449953.

45. K. De Preter, R. Barriot, F. Speleman, J. Vandesompele, Y. Moreau, Positional gene enrichment analysis of gene sets for high-resolution identification of overrepresented chromosomal regions. Nucleic Acids Res. 36, e43 (2008).

46. B. Harder, T. Jiang, T. Wu, S. Tao, M. Rojo de la Vega, W. Tian, E. Chapman, D. D. Zhang, Molecular mechanisms of Nrf2 regulation and how these influence chemical modulation for disease intervention. Biochem. Soc. Trans. 43, 680–686 (2015).

47. S. Rana, M. K. Jogi, S. Choudhary, R. Thakur, G. C. Sahoo, V. Joshi, Unraveling the intricacies of cold-inducible RNA-binding protein: A comprehensive review. Cell Stress Chaperones 29, 615–625 (2024).

48. G. P. Sykiotis, D. Bohmann, Stress-activated cap’n’collar transcription factors in aging and human disease. Sci. Signal. 3, re3 (2010).

49. H. Motohashi, M. Yamamoto, Nrf2–Keap1 defines a physiologically important stress response mechanism. Trends Mol. Med. 10, 549–557 (2004).

50. D. Firsanov, M. Zacher, X. Tian, Y. Zhao, J. C. George, T. L. Sformo, G. Tombline, S. A. Biashad, A. Gilman, N. Hamilton, A. Patel, M. Straight, M. Lee, J. Yuyang Lu, E. Haseljic, A. Williams, N. Miller, V. N. Gladyshev, Z. Zhang, J. Vijg, A. Seluanov, V. Gorbunova, DNA repair and anti-cancer mechanisms in the longest-living mammal: the bowhead whale, bioRxiv (2023)p. 2023.05.07.539748.

51. J.-K. Chen, W.-L. Lin, Z. Chen, H.-W. Liu, PARP-1–dependent recruitment of cold-inducible RNA-binding protein promotes double-strand break repair and genome stability. Proceedings of the National Academy of Sciences 115, E1759–E1768 (2018).

52. T. Wu, X.-J. Wang, W. Tian, M. C. Jaramillo, A. Lau, D. D. Zhang, Poly(ADP-ribose) polymerase-1 modulates Nrf2-dependent transcription. Free Radic. Biol. Med. 67, 69–80 (2014).

53. X. Sun, Y. Wang, K. Ji, Y. Liu, Y. Kong, S. Nie, N. Li, J. Hao, Y. Xie, C. Xu, L. Du, Q. Liu, NRF2 preserves genomic integrity by facilitating ATR activation and G2 cell cycle arrest. Nucleic Acids Res. 48, 9109–9123 (2020).

54. G. Hayashi, M. Jasoliya, S. Sahdeo, F. Saccà, C. Pane, A. Filla, A. Marsili, G. Puorro, R. Lanzillo, V. Brescia Morra, G. Cortopassi, Dimethyl fumarate mediates Nrf2-dependent mitochondrial biogenesis in mice and humans. Hum. Mol. Genet. 26, 2864–2873 (2017).

55. M. S. Brennan, M. F. Matos, B. Li, X. Hronowski, B. Gao, P. Juhasz, K. J. Rhodes, R. H. Scannevin, Dimethyl fumarate and monoethyl fumarate exhibit differential effects on KEAP1, NRF2 activation, and glutathione depletion in vitro. PLoS One 10, e0120254 (2015).

56. C. Cantó, K. J. Menzies, J. Auwerx, NAD+ Metabolism and the Control of Energy Homeostasis: A Balancing Act between Mitochondria and the Nucleus. Cell Metab. 22, 31–53 (2015).

57. J. W. Leavenworth, L. Z. Shi, X. Wang, H. Wei, Immune Cell Lineage Reprogramming in Cancer (Frontiers Media SA, 2022).

58. N. Iikuni, Q. L. K. Lam, L. Lu, G. Matarese, A. La Cava, Leptin and Inflammation. Curr. Immunol. Rev. 4, 70–79 (2008).

59. J. M. Friedman, Leptin and the endocrine control of energy balance. Nat Metab 1, 754–764 (2019).

60. H. Cui, M. López, K. Rahmouni, The cellular and molecular bases of leptin and ghrelin resistance in obesity. Nat. Rev. Endocrinol. 13, 338–351 (2017).

61. Tabula Muris Consortium, Overall coordination, Logistical coordination, Organ collection and processing, Library preparation and sequencing, Computational data analysis, Cell type annotation, Writing group, Supplemental text writing group, Principal investigators, Single-cell transcriptomics of 20 mouse organs creates a Tabula Muris. Nature 562, 367–372 (2018).

62. C. S. Greene, A. Krishnan, A. K. Wong, E. Ricciotti, R. A. Zelaya, D. S. Himmelstein, R. Zhang, B. M. Hartmann, E. Zaslavsky, S. C. Sealfon, D. I. Chasman, G. A. FitzGerald, K. Dolinski, T. Grosser, O. G. Troyanskaya, Understanding multicellular function and disease with human tissue-specific networks. Nat. Genet. 47, 569–576 (2015).

63. M. Gutierrez-Arcelus, N. Teslovich, A. R. Mola, R. B. Polidoro, A. Nathan, H. Kim, S. Hannes, K. Slowikowski, G. F. M. Watts, I. Korsunsky, M. B. Brenner, S. Raychaudhuri, P. J. Brennan, Lymphocyte innateness defined by transcriptional states reflects a balance between proliferation and effector functions. Nat. Commun. 10, 687 (2019).

64. L. Fontana, L. Partridge, Promoting health and longevity through diet: From model organisms to humans. Cell 161, 106–118 (2015).

65. X. Michelet, L. Dyck, A. Hogan, R. M. Loftus, D. Duquette, K. Wei, S. Beyaz, A. Tavakkoli, C. Foley, R. Donnelly, C. O’Farrelly, M. Raverdeau, A. Vernon, W. Pettee, D. O’Shea, B. S. Nikolajczyk, K. H. G. Mills, M. B. Brenner, D. Finlay, L. Lynch, Metabolic reprogramming of natural killer cells in obesity limits antitumor responses. Nat. Immunol. 19, 1330–1340 (2018).

66. M. A. Maxwell, G. E. O. Muscat, The NR4A subgroup: immediate early response genes with pleiotropic physiological roles. Nucl. Recept. Signal. 4, e002 (2006).

67. M. A. Pearen, G. E. O. Muscat, Minireview: Nuclear hormone receptor 4A signaling: implications for metabolic disease. Mol. Endocrinol. 24, 1891–1903 (2010).

68. E. Noro, A. Yokoyama, M. Kobayashi, H. Shimada, S. Suzuki, M. Hosokawa, T. Takehara, R. Parvin, H. Shima, K. Igarashi, A. Sugawara, Endogenous Purification of NR4A2 (Nurr1) Identified Poly(ADP-Ribose) Polymerase 1 as a Prime Coregulator in Human Adrenocortical H295R Cells. Int. J. Mol. Sci. 19 (2018).

69. J. Dabin, A. Fortuny, S. E. Polo, Epigenome maintenance in response to DNA damage. Mol. Cell 62, 712–727 (2016).

70. G. Cavalli, E. Heard, Advances in epigenetics link genetics to the environment and disease. Nature 571, 489–499 (2019).

71. J. R. Dixon, D. U. Gorkin, B. Ren, Chromatin domains: The unit of chromosome organization. Mol. Cell 62, 668–680 (2016).

72. J. A. Owen, D. Osmanović, L. Mirny, Design principles of 3D epigenetic memory systems. Science 382, eadg3053 (2023).

73. M. Kim, J. Costello, DNA methylation: an epigenetic mark of cellular memory. Exp. Mol. Med. 49, e322–e322 (2017).

74. H.-G. Lee, J. M. Rone, Z. Li, C. F. Akl, S. W. Shin, J.-H. Lee, L. E. Flausino, F. Pernin, C.-C. Chao, K. L. Kleemann, L. Srun, T. Illouz, F. Giovannoni, M. Charabati, L. M. Sanmarco, J. E. Kenison, G. Piester, S. E. J. Zandee, J. P. Antel, V. Rothhammer, M. A. Wheeler, A. Prat, I. C. Clark, F. J. Quintana, Disease-associated astrocyte epigenetic memory promotes CNS pathology. Nature 627, 865–872 (2024).

75. V. Jovasevic, E. M. Wood, A. Cicvaric, H. Zhang, Z. Petrovic, A. Carboncino, K. K. Parker, T. E. Bassett, M. Moltesen, N. Yamawaki, H. Login, J. Kalucka, F. Sananbenesi, X. Zhang, A. Fischer, J. Radulovic, Formation of memory assemblies through the DNA-sensing TLR9 pathway. Nature 628, 145–153 (2024).

76. A. Flemming, Neuronal TLR9 signalling crucial for memory formation. Nat. Rev. Immunol. 24, 306 (2024).

77. W. G. Kaelin Jr, S. L. McKnight, Influence of metabolism on epigenetics and disease. Cell 153, 56–69 (2013).

78. I. Martínez-Reyes, N. S. Chandel, Mitochondrial TCA cycle metabolites control physiology and disease. Nat. Commun. 11, 102 (2020).

79. L. Zhou, A. Nilsson, Fasting increases tissue uptake and interconversion of plasma unesterified linoleic acid in guinea pigs. Biochim. Biophys. Acta 1436, 499–508 (1999).

80. Z. Y. Chen, S. C. Cunnane, Preferential retention of linoleic acid-enriched triacylglycerols in liver and serum during fasting. Am. J. Physiol. 263, R233–9 (1992).

81. R. Groscolas, G. R. Herzberg, Fasting-induced selective mobilization of brown adipose tissue fatty acids. J. Lipid Res. 38, 228–238 (1997).

82. A. Mika, F. Macaluso, R. Barone, V. Di Felice, T. Sledzinski, Effect of exercise on fatty acid metabolism and adipokine secretion in adipose tissue. Front. Physiol. 10, 26 (2019).

83. M. A. Hidalgo, M. D. Carretta, R. A. Burgos, Long chain fatty acids as modulators of immune cells function: Contribution of FFA1 and FFA4 receptors. Front. Physiol. 12, 668330 (2021).

84. F. Marangoni, C. Agostoni, C. Borghi, A. L. Catapano, H. Cena, A. Ghiselli, C. La Vecchia, G. Lercker, E. Manzato, A. Pirillo, G. Riccardi, P. Risé, F. Visioli, A. Poli, Dietary linoleic acid and human health: Focus on cardiovascular and cardiometabolic effects. Atherosclerosis 292, 90–98 (2020).

85. C. B. Nava Lauson, S. Tiberti, P. A. Corsetto, F. Conte, P. Tyagi, M. Machwirth, S. Ebert, A. Loffreda, L. Scheller, D. Sheta, Z. Mokhtari, T. Peters, A. T. Raman, F. Greco, A. M. Rizzo, A. Beilhack, G. Signore, N. Tumino, P. Vacca, L. A. McDonnell, A. Raimondi, P. D. Greenberg, J. B. Huppa, S. Cardaci, I. Caruana, S. Rodighiero, L. Nezi, T. Manzo, Linoleic acid potentiates CD8+ T cell metabolic fitness and antitumor immunity. Cell Metab. 35, 633–650.e9 (2023).

86. A. Cui, T. Huang, S. Li, A. Ma, J. L. Pérez, C. Sander, D. B. Keskin, C. J. Wu, E. Fraenkel, N. Hacohen, Dictionary of immune responses to cytokines at single-cell resolution. Nature 625, 377–384 (2024).

87. A. Navarrete, C. P. van Schaik, K. Isler, Energetics and the evolution of human brain size. Nature 480, 91–93 (2011).

88. H. Pontzer, D. A. Raichlen, A. D. Gordon, K. K. Schroepfer-Walker, B. Hare, M. C. O’Neill, K. M. Muldoon, H. M. Dunsworth, B. M. Wood, K. Isler, J. Burkart, M. Irwin, R. W. Shumaker, E. V. Lonsdorf, S. R. Ross, Primate energy expenditure and life history. Proc. Natl. Acad. Sci. U. S. A. 111, 1433–1437 (2014).

89. T. Masuda, K. Itoh, H. Higashitsuji, H. Higashitsuji, N. Nakazawa, T. Sakurai, Y. Liu, H. Tokuchi, T. Fujita, Y. Zhao, H. Nishiyama, T. Tanaka, M. Fukumoto, M. Ikawa, M. Okabe, J. Fujita, Cold-inducible RNA-binding protein (Cirp) interacts with Dyrk1b/Mirk and promotes proliferation of immature male germ cells in mice. Proc. Natl. Acad. Sci. U. S. A. 109, 10885–10890 (2012).

90. E. Eckelhart, W. Warsch, E. Zebedin, O. Simma, D. Stoiber, T. Kolbe, T. Rülicke, M. Mueller, E. Casanova, V. Sexl, A novel Ncr1-Cre mouse reveals the essential role of STAT5 for NK-cell survival and development. Blood 117, 1565–1573 (2011).

91. B. Kadkhodaei, T. Ito, E. Joodmardi, B. Mattsson, C. Rouillard, M. Carta, S.-I. Muramatsu, C. Sumi-Ichinose, T. Nomura, D. Metzger, P. Chambon, E. Lindqvist, N.-G. Larsson, L. Olson, A. Björklund, H. Ichinose, T. Perlmann, Nurr1 is required for maintenance of maturing and adult midbrain dopamine neurons. J. Neurosci. 29, 15923–15932 (2009).

92. S. A. Hirota, A. Ueno, S. E. Tulk, H. M. Becker, L. P. Schenck, M. S. Potentier, Y. Li, S. Ghosh, D. A. Muruve, J. A. MacDonald, P. L. Beck, Exaggerated IL-15 and Altered Expression of foxp3+ Cell-Derived Cytokines Contribute to Enhanced Colitis in Nlrp3-/- Mice. Mediators Inflamm. 2016, 5637685 (2016).

93. S. S. P. Rao, M. H. Huntley, N. C. Durand, E. K. Stamenova, I. D. Bochkov, J. T. Robinson, A. L. Sanborn, I. Machol, A. D. Omer, E. S. Lander, E. L. Aiden, A 3D map of the human genome at kilobase resolution reveals principles of chromatin looping. Cell 159, 1665–1680 (2014).

94. H. Kanno, M. Nose, J. Itoh, Y. Taniguchi, M. Kyogoku, Spontaneous development of pancreatitis in the MRL/Mp strain of mice in autoimmune mechanism. Clin. Exp. Immunol. 89, 68–73 (1992).

95. G. Paxinos, C. Watson, The Rat Brain in Stereotaxic Coordinates (Academic Press, 2013).

96. M. Palkovits, Punch sampling biopsy technique. Methods Enzymol. 103, 368–376 (1983).

97. M. P. Murray, I. Engel, G. Seumois, S. Herrera-De la Mata, S. L. Rosales, A. Sethi, A. Logandha Ramamoorthy Premlal, G.-Y. Seo, J. Greenbaum, P. Vijayanand, J. P. Scott-Browne, M. Kronenberg, Transcriptome and chromatin landscape of iNKT cells are shaped by subset differentiation and antigen exposure. Nat. Commun. 12, 1446 (2021).

98. N. M. LaMarche, H. Kane, A. C. Kohlgruber, H. Dong, L. Lynch, M. B. Brenner, Distinct iNKT cell populations use IFNγ or ER stress-induced IL-10 to control adipose tissue homeostasis. Cell Metab. 32, 243–258.e6 (2020).

99. H. Kane, N. M. LaMarche, Á. Ní Scannail, A. E. Garza, H.-F. Koay, A. I. Azad, B. Kunkemoeller, B. Stevens, M. B. Brenner, L. Lynch, Longitudinal analysis of invariant natural killer T cell activation reveals a cMAF-associated transcriptional state of NKT10 cells. Elife 11, e76586 (2022).

