## Supplementary materials for "Convergent Signals Encode Persistent Memory of Resilient States"

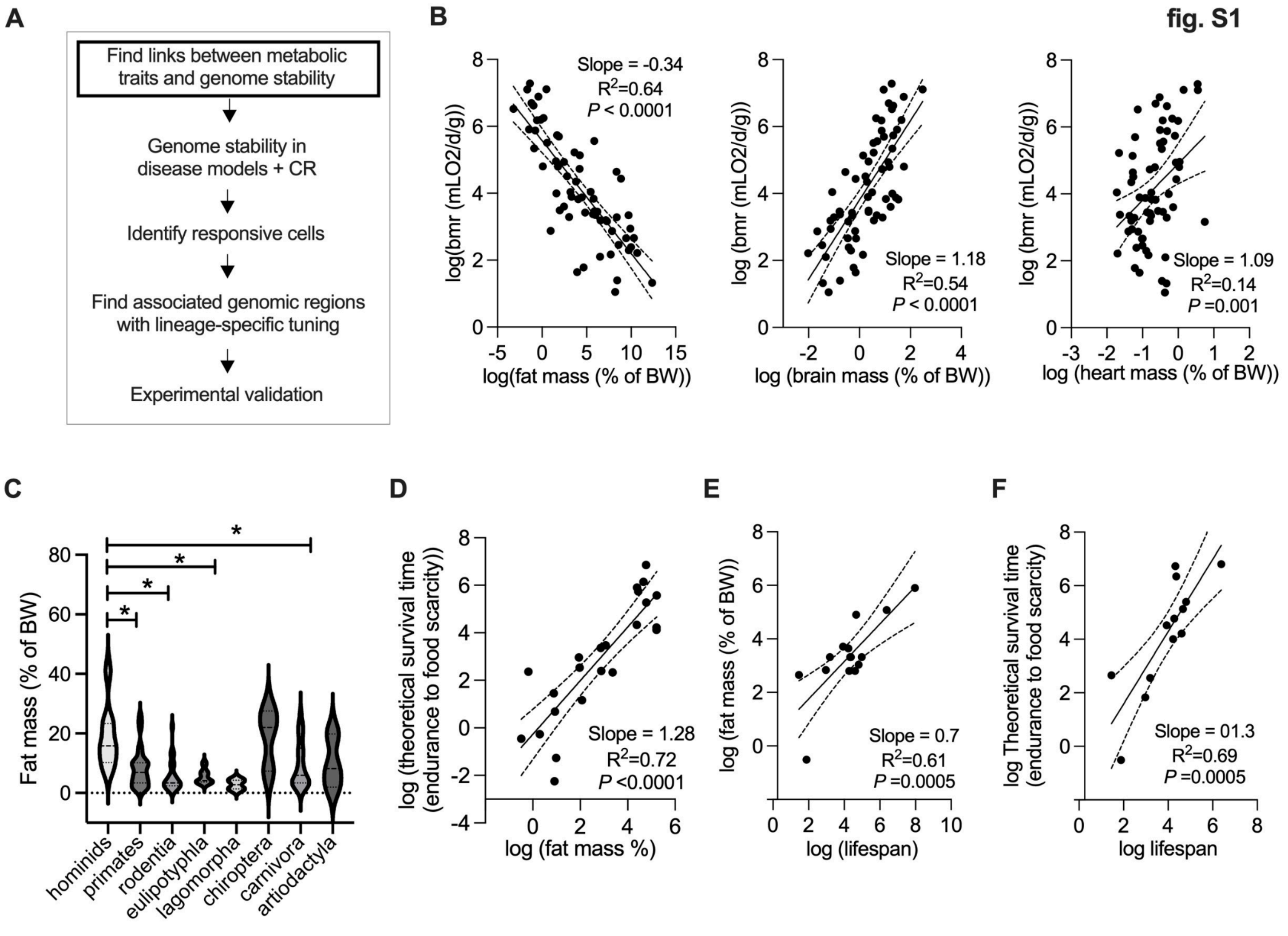


**Fig. S1 Relationships between metabolic traits and genome stability in mammals.** *(related to Fig. 1)*
(**A**) Illustration of workflow overview.
(**B**) Left panel, normalized basal metabolic rate (BMR) per gram data across mammals and body fat composition (BFC; data from Navarrete et al. (*87*)) data were transformed using binay logarithmic approach (Methods) and linearized to evaluate dependencies using linear regression (shaded line is 95% CI). Middle panel, BMR/g and brain mass percentage. Right panel, BMR/g and heart mass percentage comparison.
(**C**) BFC across taxonomic mammalian groups (data from references (*8*, *87*, *88*)).
(**D**) Theoretical survival time (Methods) and BFC transformed and analyzed as in B
(**E**) BFC from several curated sources (methods) and lifespan (data from references (*15*, *16*)) transformed and analyzed as in B.
(**F**) Theoretical survival time (Methods) and lifespan transformed and analyzed as in B.
Data show mean values per group and SEM. Unpaired, two-tailed student's t-test was used when two groups were compared, and ANOVA followed by fisher's least significant difference (LSD) test for post hoc comparisons for multiple groups. * p-value <0.05.

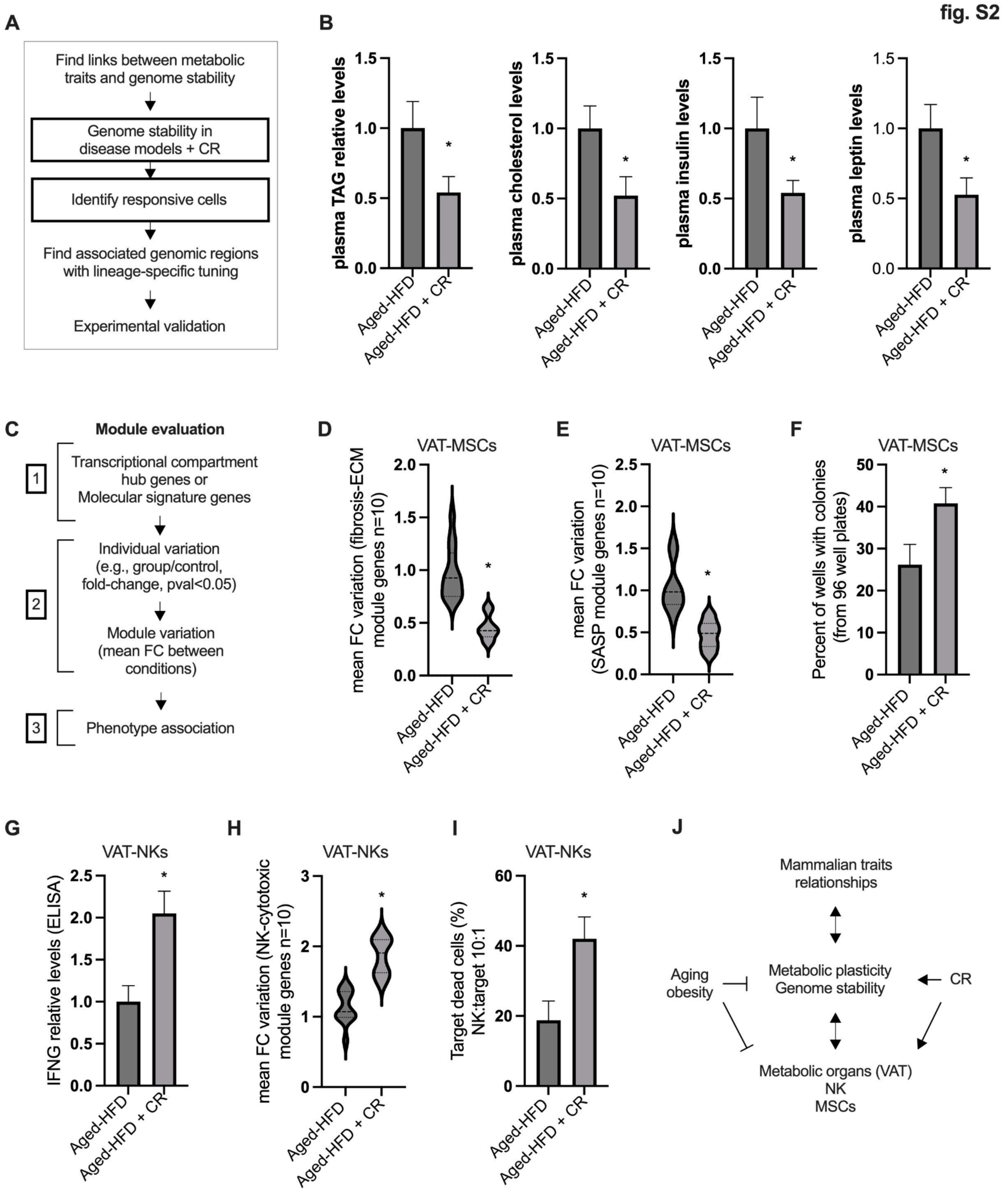


**Fig. S2. Oxidative stress and DNA damage in adipose tissue are reduced under caloric restriction.** *(related to Fig. 1)*
(**A**) Illustration of workflow overview.

(**B**) Panels from left to right, relative plasma levels of triglycerides (TAG), cholesterol, insulin, and leptin (n=4-6)
(**C**) Illustration of workflow overview for module evaluation of gene programs.
(**D**) As in C, fibrosis-ECM gene program activity (10 genes, Table S2, mice n=4-6).
(**E**) As in C, senescence-associated secretory phenotype (SASP) gene program activity (10 genes, Table S2, mice n=4-6).
(**F**) Colony forming assays from extracted VAT-MSCs. Cells were plated in 96 well plates and data is shown as the percent of wells with colonies for time points and conditions.
(**G**) Relative IFNG levels by ELISA form extracted VAT-NK cells (n=4-6).
(**H**) As in C, NK-cytotoxic gene program activity (10 genes, Table S2, mice n=4-6).
(**I**) Dead target cell cytotoxic assay. VAT-NKs from indicated models were co-incubated with VAT-MSCs from the aged-HFD model. Percentage of dead target cells stained with propidium iodide (Pi).
(**J**) Illustration of summary overview.
Animal experiments were done with n=4-6 mice per group. Data show mean values per group and SEM. Cell extraction experiments were done in each mice per group and mean per groups were compared. Unpaired, two-tailed student's t-test was used when two groups were compared, and ANOVA followed by fisher's least significant difference (LSD) test for post hoc comparisons for multiple groups. * p-value <0.05.

fig. S3

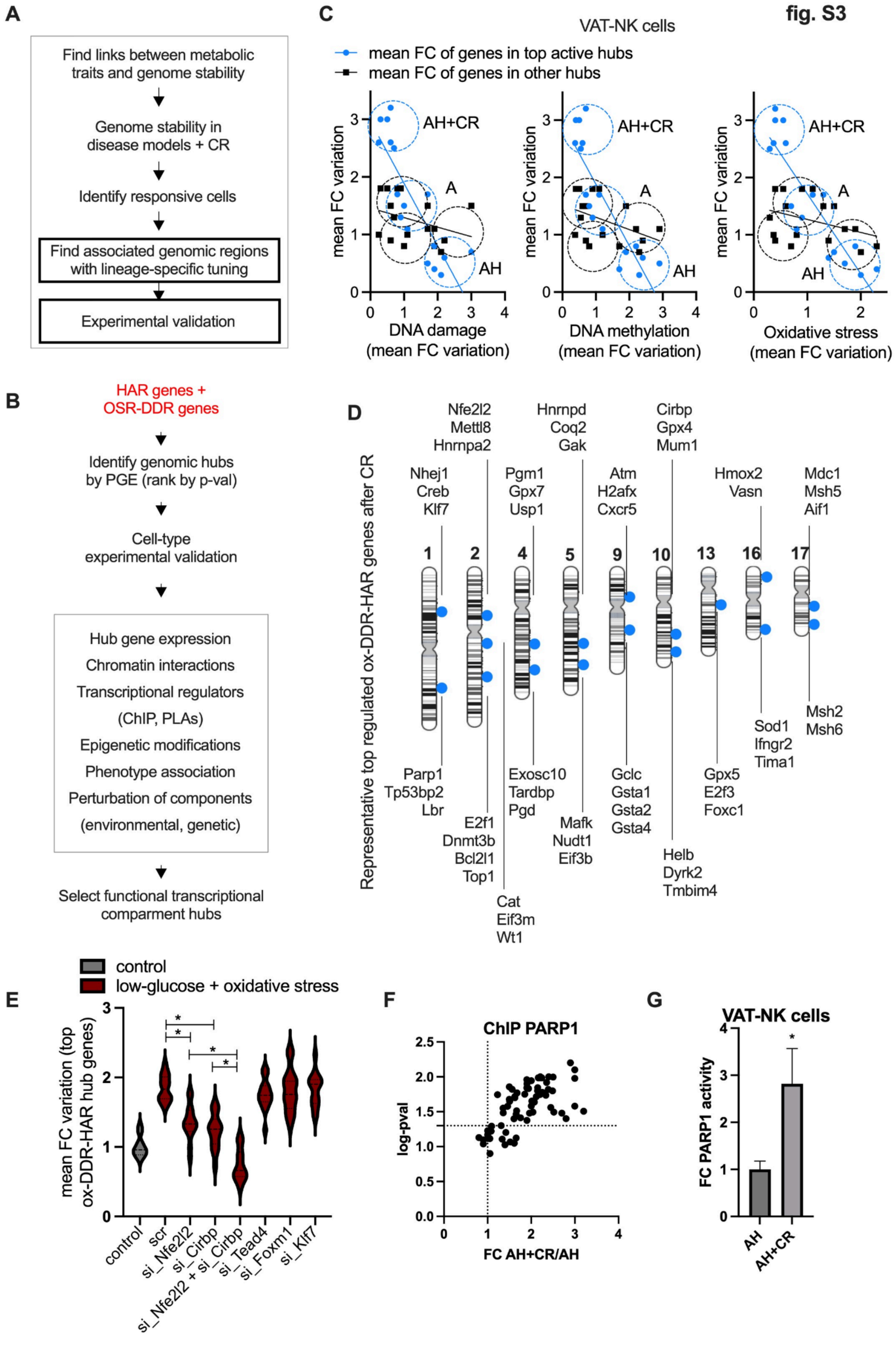

A
Find links between metabolic traits and genome stability
Genome stability in disease models + CR
Identify responsive cells
Find associated genomic regions with lineage-specific tuning
Experimental validation
B
HAR genes + OSR-DDR genes
Identify genomic hubs by PGE (rank by p-val)
Cell-type experimental validation
Hub gene expression
Chromatin interactions
Transcriptional regulators
(ChIP, PLAs)
Epigenetic modifications
Phenotype association
Perturbation of components
(environmental, genetic)
Select functional transcriptional comparment hubs
C
VAT-NK cells
mean FC of genes in top active hubs
mean FC of genes in other hubs
AH+CR
A
AH
mean FC variation
DNA damage (mean FC variation)
DNA methylation (mean FC variation)
Oxidative stress (mean FC variation)
D
Representative top regulated ox-DDR-HAR genes after CR
1 2 4 5 9 10 13 16 17
Nhej1 Creb Klf7
Nfe2l2 Mettl8 Hnrnpa2
Pgm1 Gpx7 Usp1
Hnrnpd Coq2 Gak
Atm H2afx Cxcr5
Cirbp Gpx4 Mum1
Hmox2 Vasn
Mdc1 Msh5 Aif1
Parp1 Tp53bp2 Lbr
E2f1 Dnmt3b Bcl2l1 Top1
Cat Eif3m Wt1
Exosc10 Tardbp Pgd
Mafk Nudt1 Eif3b
Gclc Gsta1 Gsta2 Gsta4
Helb Dyrk2 Tmbim4
Gpx5 E2f3 Foxc1
Sod1 Ifngr2 Tima1
Msh2 Msh6
E
control
low-glucose + oxidative stress
mean FC variation (top ox-DDR-HAR hub genes)
control
scr
si_Nfe2l2
si_Cirbp
si_Nfe2l2 + si_Cirbp
si_Tead4
si_Foxm1
si_Klf7
F
ChIP PARP1
log-pval
FC AH+CR/AH
G
VAT-NK cells
FC PARP1 activity
AH
AH+CR

**Fig. S3. Transcriptional hubs with HARs linked to OSR and DDR programs in VAT-NK cells.** *(related to Fig. 2)*.

(**A**) Illustration of workflow overview.

(**B**) Illustration of workflow overview of highlighted boxes from A.

(**C**) Fold-to-fold change plots from extracted VAT-NK cells from mice models. Y-axes show mean fold-change variation genes in top active hubs and genes in other hubs (n=20 genes per hub group, hubs ranked by PGE p-value, number of genes in hub, and contextual activity) in each mouse per model group. X-axes show phenotypic variations for each mouse. Lines represent linear regressions.

(**D**) Illustration of selected ox-DDR-HAR hubs (positioned blue dots in chromosomes) and highly active genes.

(**E**) Module evaluation for ox-DDR-HAR hub genes (n=15 genes) in VAT-NK cells treated with scramble and specific siRNAs, and comparing the effect of low-glucose (control 5.5 mM, low glucose is 2 mM) and oxidative stress treatment (hydrogen peroxide, 250-500 μM).

(**F**) ChIP-qPCR fold-change binding of PARP1 to ox-DDR-HAR hub promoter regions.

(**G**) PARP1 activity in extracted VAT-NK cells from different models.

Animal experiments were done with n=4-6 mice per group. Data show mean values per group and SEM. Cell extraction experiments were done in each mice per group and mean per groups were compared when indicated. Unpaired, two-tailed student's t-test was used when two groups were compared, and ANOVA followed by fisher's least significant difference (LSD) test for post hoc comparisons for multiple groups. * p-value $<0.05$.

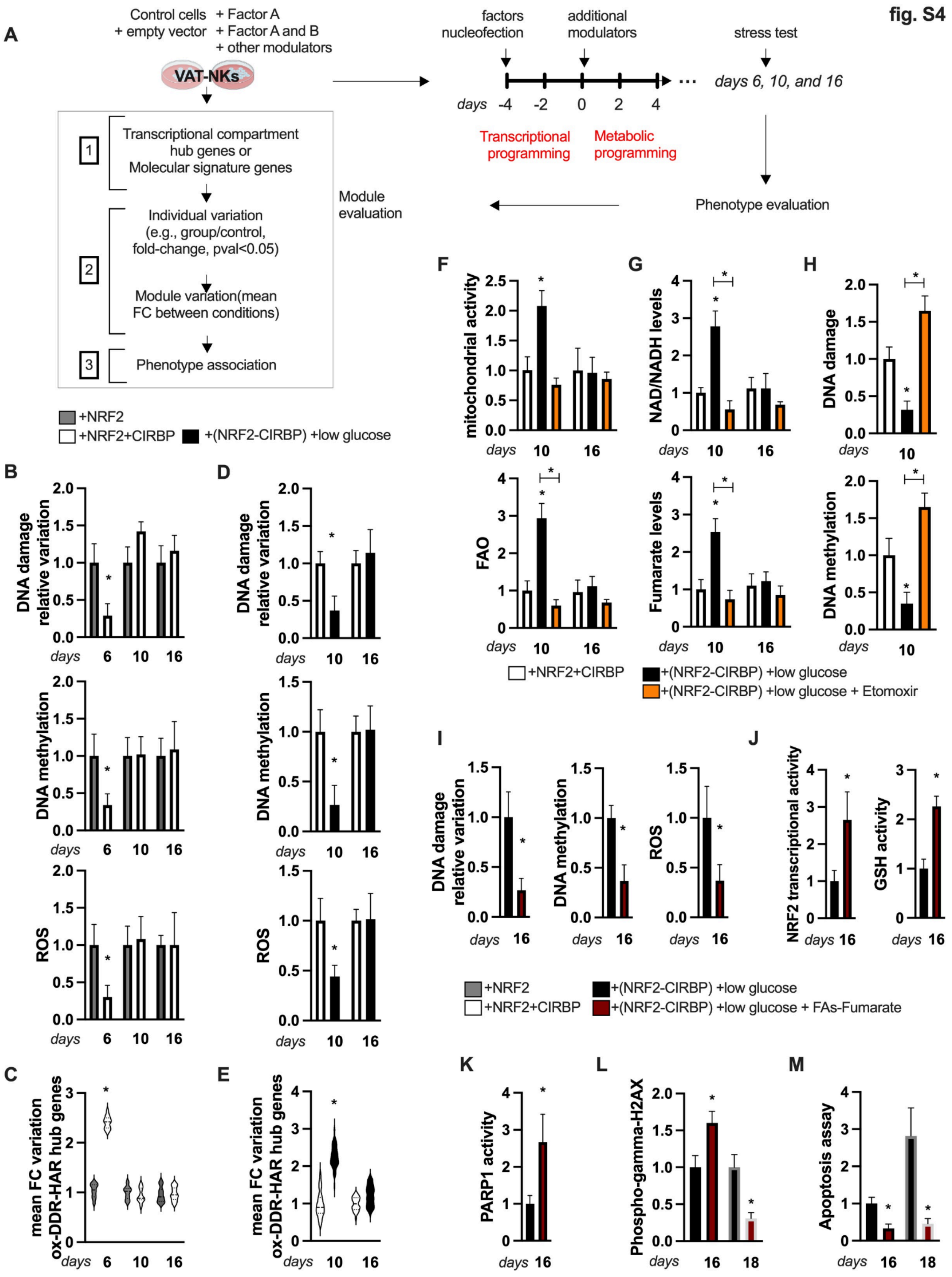
fig. S4
A
Control cells + empty vector
+ Factor A
+ Factor A and B
+ other modulators
VAT-NKs
1 Transcriptional compartment hub genes or Molecular signature genes
2 Individual variation (e.g., group/control, fold-change, pval<0.05)
Module variation(mean FC between conditions)
3 Phenotype association
Module evaluation
factors nucleofection
additional modulators
stress test
days -4 -2 0 2 4
days 6, 10, and 16
Transcriptional programming
Metabolic programming
Phenotype evaluation
+NRF2
+NRF2+CIRBP
+(NRF2-CIRBP) +low glucose
B
D
DNA damage relative variation
DNA methylation
ROS
C
E
mean FC variation ox-DDR-HAR hub genes
F
G
H
mitochondrial activity
NAD/NADH levels
DNA damage
FAO
Fumarate levels
DNA methylation
+(NRF2-CIRBP) +low glucose + Etomoxir
I
J
NRF2 transcriptional activity
GSH activity
+(NRF2-CIRBP) +low glucose + FAs-Fumarate
K
L
M
PARP1 activity
Phospho-gamma-H2AX
Apoptosis assay
days

**Fig. S4. Engineered CR-like NK cells in vitro.** *(related to Fig. 2)*
(**A**) Illustration of workflow overview.
(**B**) Top panel, relative DNA damage variation in indicated VAT-NK cells, 12 hours post oxidative stress test (hydrogen peroxide, 250 µM) in indicated days. Middle panel, relative DNA methylation. Lower panel, reactive oxygen species (ROS) assay levels.
(**C**) Module evaluation for the expression variation of ox-DDR-HAR hub genes (n=15 genes) in VAT-NK cells treated as in B.
(**D**) As in B, in indicated cells, from top to bottom, relative DNA damage, relative DNA methylation, and relative ROS levels (control 5.5 mM, low glucose is 2 mM).
(**E**) As in C, module evaluation for ox-DDR-HAR hub genes.
(**F**) Top panel, relative basal mitochondrial activity of indicated cell groups, and using etomoxir (50µM). Lower panel, relative fatty acid oxidation (FAO) levels (treated with FCCP 1 µM).
(**G**) As in F. Top panel, NAD+ relative levels. Lower panel, relative fumarate levels.
(**H**) As in F. Top panel, relative DNA damage. Lower panel, relative DNA methylation.
(**I**) VAT-NK cells treated as in D, and as indicated, an additional group of VAT-NK cells are treated with fatty acids (palmitate, 50 µM) and fumarate (50 µM). From left to right, relative DNA damage, relative DNA methylation, and relative ROS levels post stress test.
(**J**) As in I. Left panel, relative NRF2 transcriptional activity. Right panel, relative glutathione activity.
(**K**) As in I, relative PARP1 activity.
(**L**) Cells as in I, relative H2AX phosphorylation assay by ELISA 12h post-stress test and 2 days after.
(**M**) As in M, relative apoptosis levels.
Animal experiments were done with n=4-6 mice per group. Cell experiments were done with 3 independent replicates. Data show mean values per group and SEM. Cell extraction experiments were done in each mice per group and mean per groups were compared when indicated. Unpaired, two-tailed student's t-test was used when two groups were compared, and ANOVA followed by fisher's least significant difference (LSD) test for post hoc comparisons for multiple groups. * p-value <0.05.

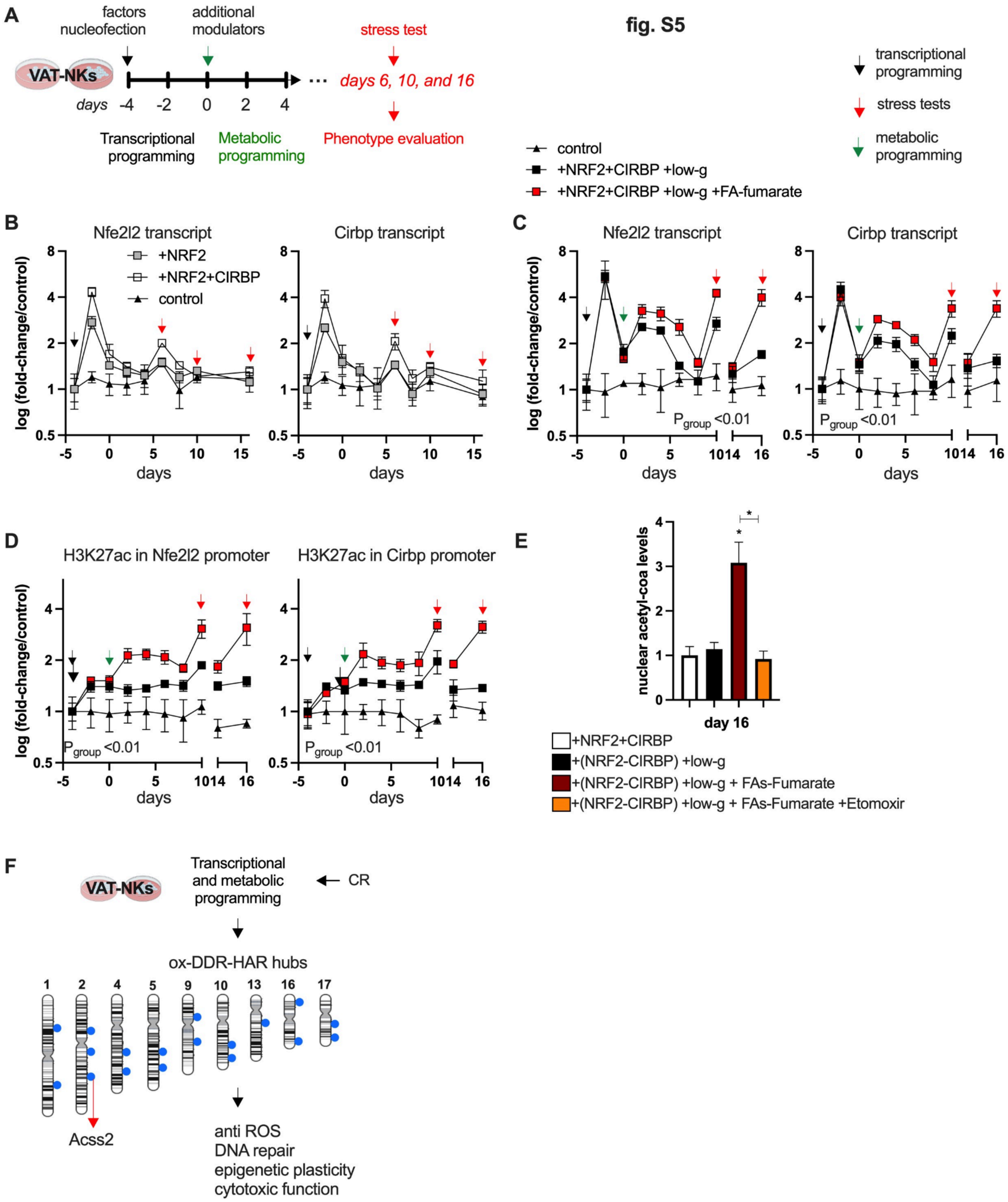
fig. S5
A
factors nucleofection
additional modulators
stress test
VAT-NKs
days -4 -2 0 2 4
days 6, 10, and 16
Transcriptional programming
Metabolic programming
Phenotype evaluation
transcriptional programming
stress tests
metabolic programming
control
+NRF2+CIRBP +low-g
+NRF2+CIRBP +low-g +FA-fumarate
B
Nfe2l2 transcript
Cirbp transcript
+NRF2
+NRF2+CIRBP
control
log (fold-change/control)
days
C
Nfe2l2 transcript
Cirbp transcript
P group <0.01
D
H3K27ac in Nfe2l2 promoter
H3K27ac in Cirbp promoter
E
nuclear acetyl-coa levels
day 16
+NRF2+CIRBP
+(NRF2-CIRBP) +low-g
+(NRF2-CIRBP) +low-g + FAs-Fumarate
+(NRF2-CIRBP) +low-g + FAs-Fumarate +Etomoxir
F
VAT-NKs
Transcriptional and metabolic programming
CR
ox-DDR-HAR hubs
1 2 4 5 9 10 13 16 17
Acss2
anti ROS
DNA repair
epigenetic plasticity
cytotoxic function

**Fig. S5. Engineered CR-like NK cells in vitro.** *(related to Fig. 2)*

(**A**) Illustration of workflow overview.
(**B**) Temporal gene expression in VAT-NK cells treated as indicated, and undergoing independent oxidative stress tests (red arrow, hydrogen peroxide, 250 μM).
(**C**) Temporal gene expression in VAT-NK cells treated as indicated and with stress test as in B. Control glucose media (5.5 mM), low glucose (2 mM), fatty acids (palmitate, 50 μM), and fumarate (50 μM).
(**D**) Temporal H3K27ac levels (by ChIP) in promoters from cells as in C.
(**E**) Relative acetyl-coa levels in nuclear extracts from indicated cells.
(**F**) Summary illustration.
Primary cell experiments were done with 3 independent replicates. Data show mean values per group and SEM. Cell extraction experiments were done in each mice per group and mean per groups were compared when indicated. Unpaired, two-tailed student's t-test was used when two groups were compared, and ANOVA followed by fisher's least significant difference (LSD) test for post hoc comparisons for multiple groups. * $p$-value $<0.05$.

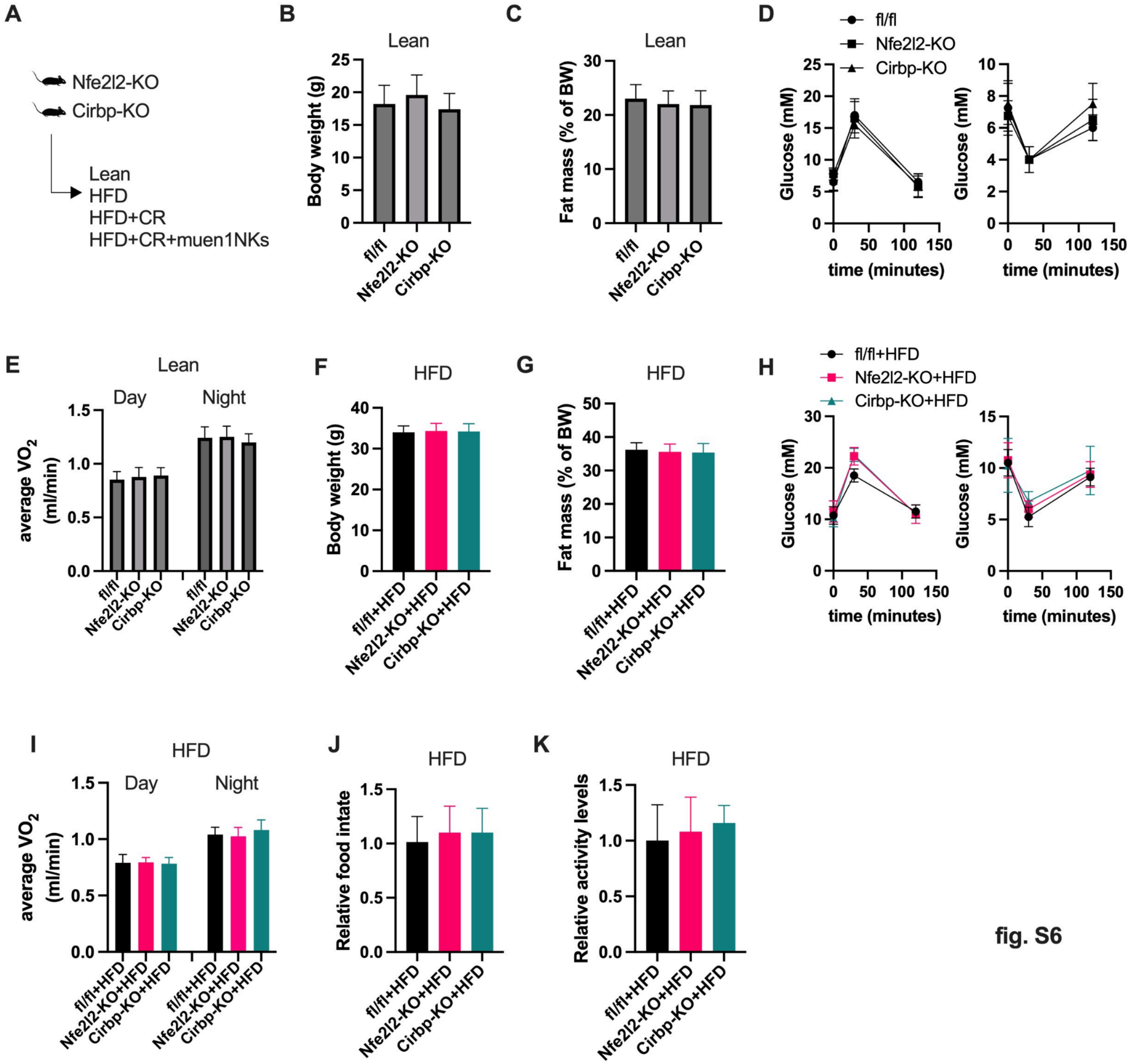


fig. S6

**Fig. S6. CR-mediators KOs.** *(related to Fig. 3)*

(**A**) Illustration of murine models and interventions.

(**B**) Body weight (BW, grams).

(**C**) Fat mass composition (% of BW)

(**D**) Left panel, glucose tolerance test (GTT). Right panel, insulin tolerance test (ITT)

(**E**) Average $VO_2$ levels from 48 hours measurements in metabolic cages.

(**F**) BW (grams).

(**G**) Fat mass composition (% of BW)

(**H**) Left panel, glucose tolerance test (GTT). Right panel, insulin tolerance test (ITT)

(**I**) Average $VO_2$ levels as in E.

(**J**) Relative food intake.

(**K**) Relative activity levels.

Animal experiments were done with n=4-6 mice per group. Data show mean values per group and SEM. Cell extraction experiments were done in each mice per group and mean per groups were compared. Unpaired, two-tailed student's t-test was used when two groups were compared, and ANOVA followed by fisher's least significant difference (LSD) test for post hoc comparisons for multiple groups. Two-way ANOVA was used to estimate significance between groups constrained by time measurements, and Tukey test for multiple comparisons * p-value <0.05.

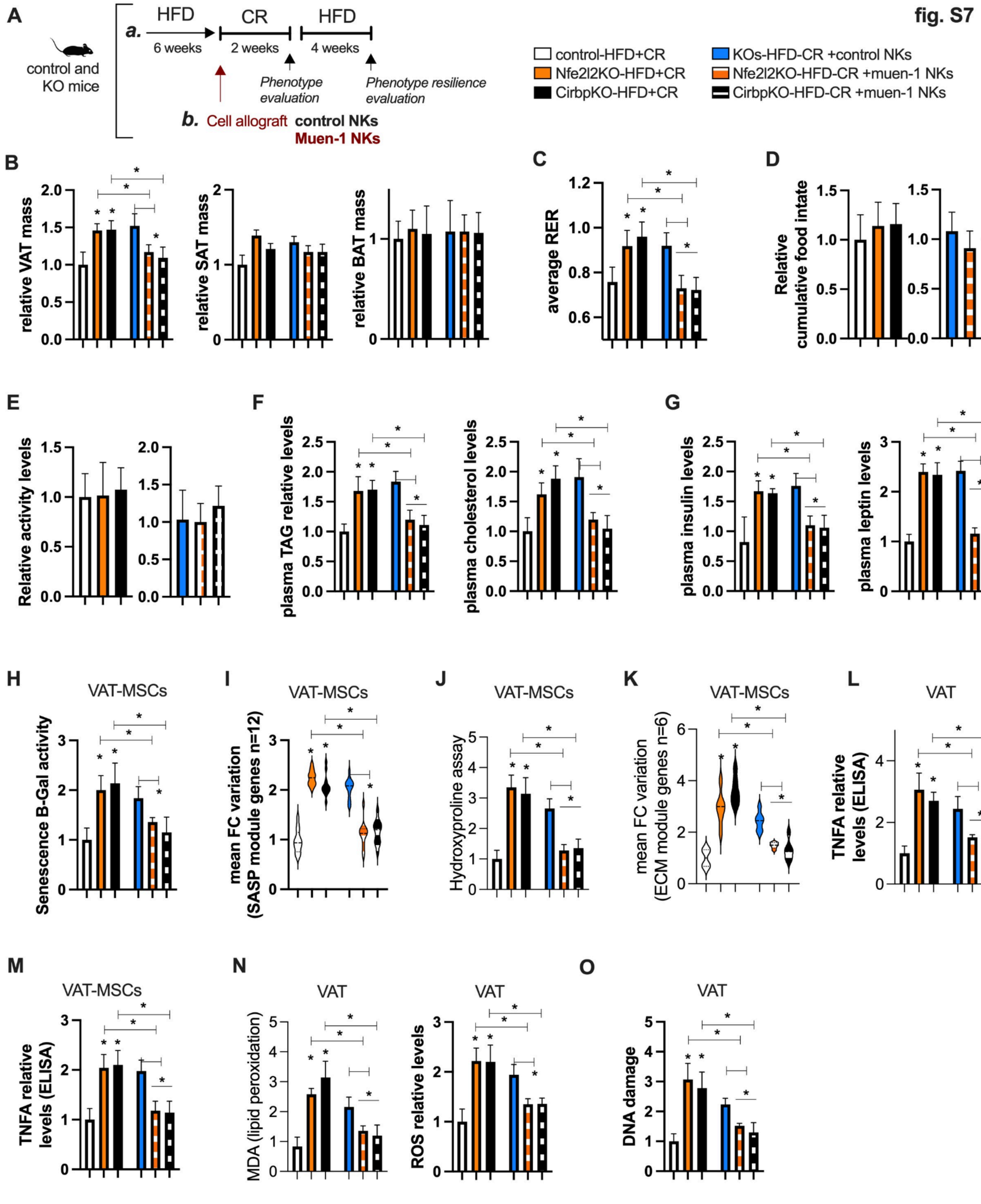
fig. S7
A
control and KO mice
a. HFD 6 weeks
CR 2 weeks
HFD 4 weeks
Phenotype evaluation
Phenotype resilience evaluation
b. Cell allograft control NKs
Muen-1 NKs
control-HFD+CR
Nfe2l2KO-HFD+CR
CirbpKO-HFD+CR
KOs-HFD-CR +control NKs
Nfe2l2KO-HFD-CR +muen-1 NKs
CirbpKO-HFD-CR +muen-1 NKs
B
relative VAT mass
relative SAT mass
relative BAT mass
C
average RER
D
Relative cumulative food intate
E
Relative activity levels
F
plasma TAG relative levels
plasma cholesterol levels
G
plasma insulin levels
plasma leptin levels
H
VAT-MSCs
Senescence B-Gal activity
I
VAT-MSCs
mean FC variation (SASP module genes n=12)
J
VAT-MSCs
Hydroxyproline assay
K
VAT-MSCs
mean FC variation (ECM module genes n=6)
L
VAT
TNFA relative levels (ELISA)
M
VAT-MSCs
TNFA relative levels (ELISA)
N
VAT
MDA (lipid peroxidation)
VAT
ROS relative levels
O
VAT
DNA damage

**Fig. S7. CR-like NK cells restore impairments from Nrf2 or Cirbp deletion** *(related to Fig. 3)*

(**A**) Illustration of workflow overview.
(**B**) Fat mass of different adipose depots relative to body weight (BW).
(**C**) Average $VO_2$ levels (night) from 48 hours measurements in metabolic cages.
(**D**) Relative food intake.
(**E**) Relative activity levels.
(**F**) Relative plasma levels of triacylglycerol (TAGs) and cholesterol.
(**G**) Relative plasma levels of insulin (TAGs) and leptin.
(**H**) Relative senescence B-gal assay activity in VAT-MSCs.
(**I**) Module evaluation of fold-change (FC) variation across SASP genes (n=10 genes) in VAT-MSCs.
(**J**) Relative hydroxyproline assay activity in VAT-MSCs.
(**K**) Module evaluation of fold-change (FC) variation across fibrosis-associated ECM genes (n=10 genes) in VAT-MSCs.
(**L**) Relative TNFA levels by ELISA in VAT and in VAT-MSCs (**M**).
(**N**) Relative MDA (left panel) and ROS levels (right panel) in VAT.
(**O**) Relative DNA damage in VAT.
Animal experiments were done with n=4-6 mice per group. Data show mean values per group and SEM. Cell extraction experiments were done in each mice per group and mean per groups were compared. Unpaired, two-tailed student's t-test was used when two groups were compared, and ANOVA followed by fisher's least significant difference (LSD) test for post hoc comparisons for multiple groups. * p-value <0.05.

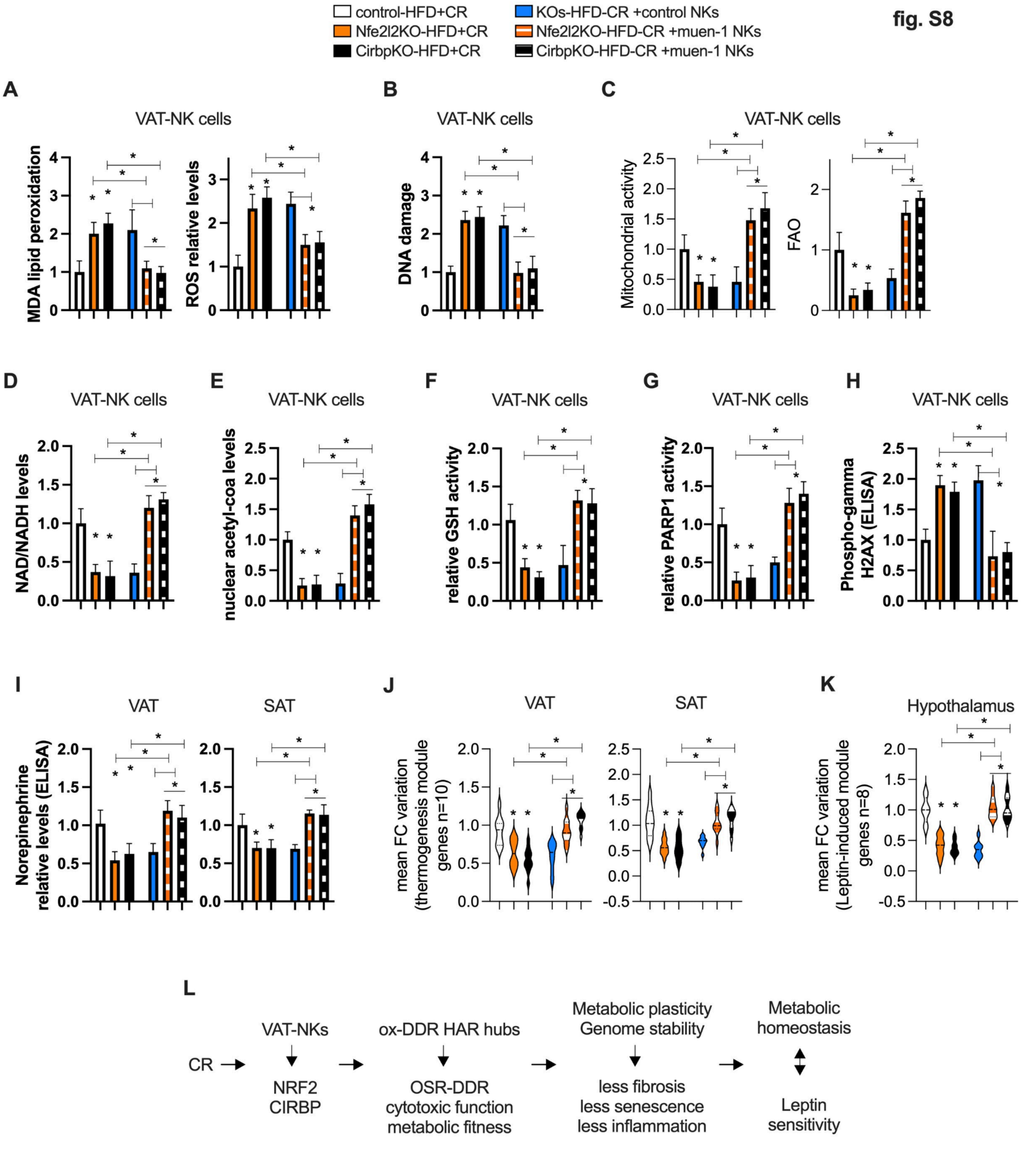


**Fig. S8. CR-like NK cells restore impairments from Nrf2 or Cirbp deletion and enhance leptin sensitivity** *(related to Fig. 3)*

(**A**) Relative MDA (left panel) and ROS levels (right panel) in VAT-NK cells.

(**B**) Relative DNA damage in VAT-NK cells.

(**C**) Relative basal mitochondrial activity (left panel) and FAO (right panel, treated with FCCP 1 μM).

(**D**) Relative NAD+ levels.

(**E**) Relative acetyl-coa levels in nuclear extracts from VAT-NK cells.
(**F**) Relative glutathione activity.
(**G**) Relative PARP1 activity.
(**H**) Relative H2AX phosphorylation assay by ELISA.
(**I**) Relative norepinephrine levels by ELISA.
(**J**) Module evaluation of fold-change (FC) variation across thermogenesis-associated genes (n=10 genes) in white fat depots.
(**K**) Module evaluation of fold-change (FC) variation across leptin-responsive genes (n=10 genes).
(**L**) Summary illustration.
Animal experiments were done with n=4-6 mice per group. Data show mean values per group and SEM. Cell extraction experiments were done in each mice per group and mean per groups were compared. Unpaired, two-tailed student's t-test was used when two groups were compared, and ANOVA followed by fisher's least significant difference (LSD) test for post hoc comparisons for multiple groups. * p-value <0.05.

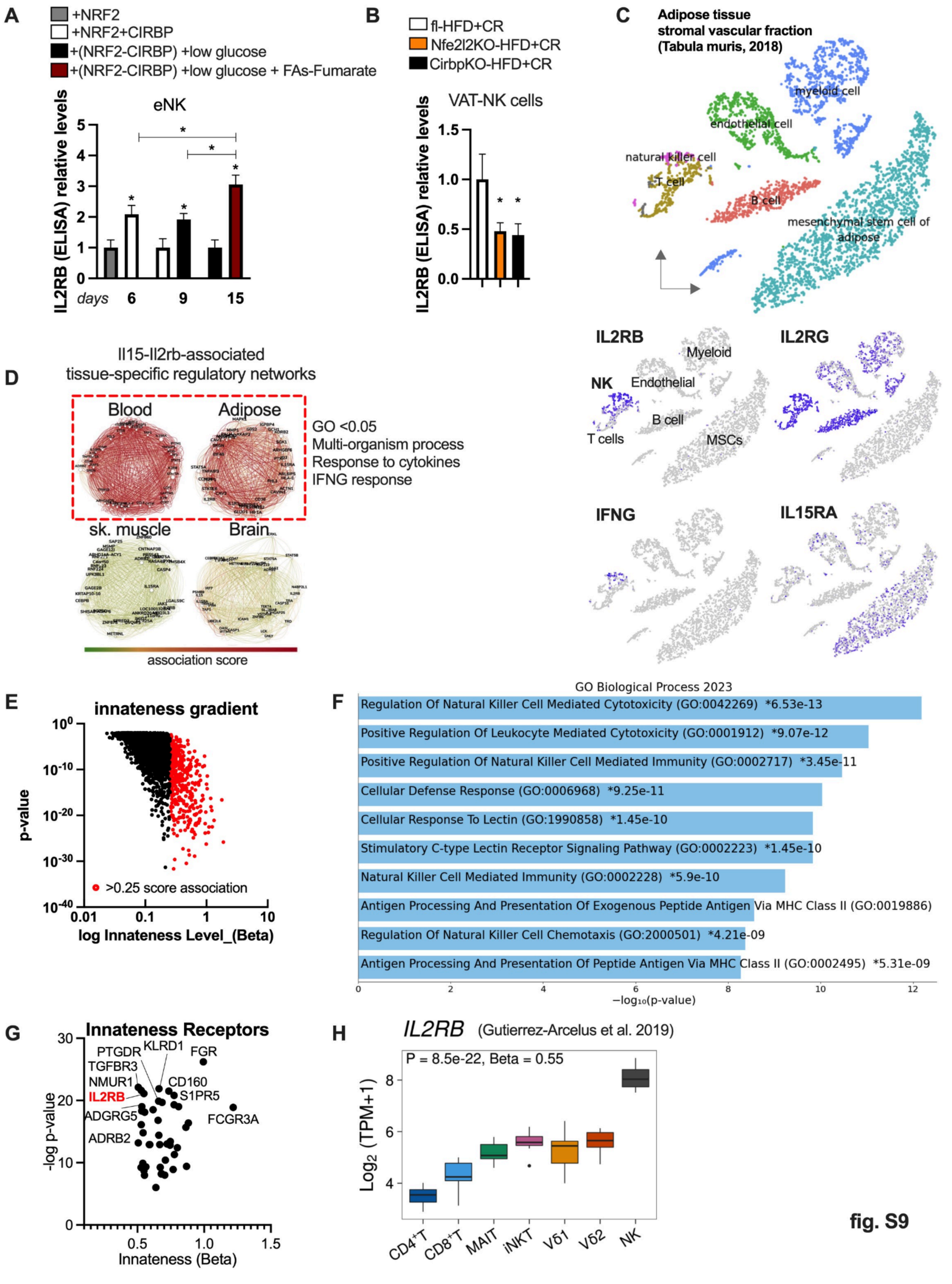
A
+NRF2
+NRF2+CIRBP
+(NRF2-CIRBP) +low glucose
+(NRF2-CIRBP) +low glucose + FAs-Fumarate
eNK
IL2RB (ELISA) relative levels
days 6 9 15
B
fl-HFD+CR
Nfe2l2KO-HFD+CR
CirbpKO-HFD+CR
VAT-NK cells
IL2RB (ELISA) relative levels
C
Adipose tissue stromal vascular fraction (Tabula muris, 2018)
myeloid cell
endothelial cell
natural killer cell
T cell
B cell
mesenchymal stem cell of adipose
IL2RB
IL2RG
IFNG
IL15RA
Myeloid
NK
Endothelial
B cell
T cells
MSCs
D
Il15-Il2rb-associated tissue-specific regulatory networks
Blood
Adipose
sk. muscle
Brain
GO <0.05
Multi-organism process
Response to cytokines
IFNG response
association score
E
innateness gradient
p-value
>0.25 score association
log Innateness Level_(Beta)
F
GO Biological Process 2023
Regulation Of Natural Killer Cell Mediated Cytotoxicity (GO:0042269) *6.53e-13
Positive Regulation Of Leukocyte Mediated Cytotoxicity (GO:0001912) *9.07e-12
Positive Regulation Of Natural Killer Cell Mediated Immunity (GO:0002717) *3.45e-11
Cellular Defense Response (GO:0006968) *9.25e-11
Cellular Response To Lectin (GO:1990858) *1.45e-10
Stimulatory C-type Lectin Receptor Signaling Pathway (GO:0002223) *1.45e-10
Natural Killer Cell Mediated Immunity (GO:0002228) *5.9e-10
Antigen Processing And Presentation Of Exogenous Peptide Antigen Via MHC Class II (GO:0019886)
Regulation Of Natural Killer Cell Chemotaxis (GO:2000501) *4.21e-09
Antigen Processing And Presentation Of Peptide Antigen Via MHC Class II (GO:0002495) *5.31e-09
−log₁₀(p-value)
G
Innateness Receptors
PTGDR
KLRD1
FGR
TGFBR3
NMUR1
CD160
IL2RB
S1PR5
ADGRG5
FCGR3A
ADRB2
-log p-value
Innateness (Beta)
H
IL2RB (Gutierrez-Arcelus et al. 2019)
P = 8.5e-22, Beta = 0.55
Log2 (TPM+1)
CD4+T
CD8+T
MAIT
iNKT
Vδ1
Vδ2
NK

fig. S9

**Fig. S9. DR regulates IL15-IL2RB and innateness signatures** *(related to Fig. 4).*
(**A**) Relative IL2RB levels by ELISA in engineered VAT-NKs (eNKs) post oxidative stress tests in indicated days (hydrogen peroxide, 250 μM).
(**B**) Relative IL2RB levels by ELISA in VAT-NKs from KO mice models and control models post HFD (6 weeks) and subjected to CR (2 weeks).
(**C**) Umap scRNAseq plots of adipose tissue stromal vascular fraction from the tabula muris repository. Lower panel, expression of key genes on associated clusters.
(**D**) Gene regulatory networks (repository from Greene et al (*62*)) associated with IL2RB andIL15 across metabolic tissues, and functional annotation (shaded red box).
(**E**) Innateness gradients across lymphocyte human population from scRNA-seq (data from Gutierrez-Arcelus et al. (*63*)). Association beta score was downloaded and plotted, showing top innateness associated gene signatures (in red).
**(F**) Functional annotation of biological process enrichments using enrichR for the innateness gene signature.
(**G**) Receptors linked to innateness (associated signature as in E).
(**H**) Il2rb expression across lymphocytes and innateness beta score (data from Gutierrez-Arcelus et al. (*63*)). Animal experiments were done with n=4-6 mice per group. Data show mean values per group and SEM. Cell extraction experiments were done in each mice per group and mean per groups were compared. Unpaired, two-tailed student's t-test was used when two groups were compared, and ANOVA followed by fisher's least significant difference (LSD) test for post hoc comparisons for multiple groups. * p-value <0.05.

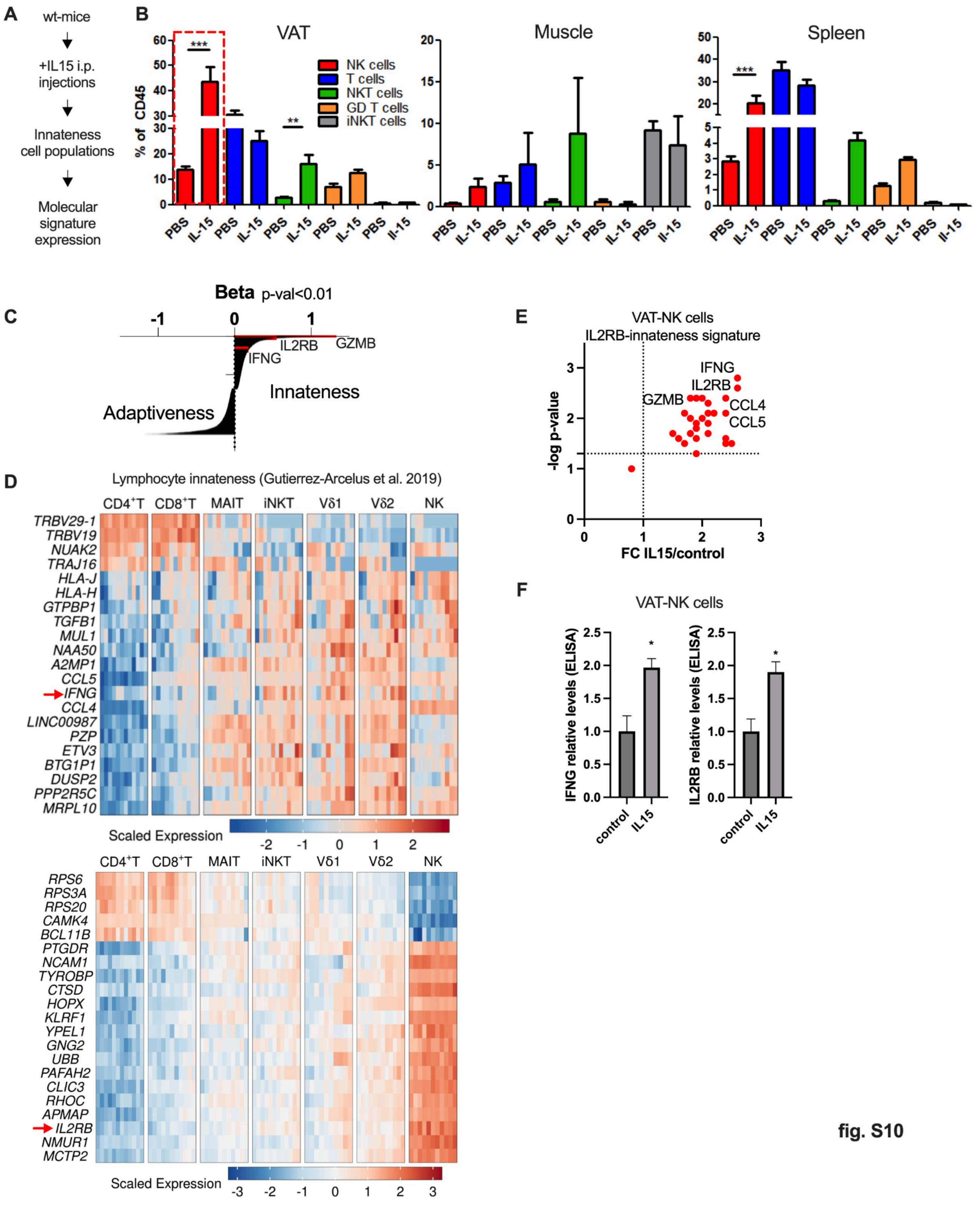
A
wt-mice
+IL15 i.p. injections
Innateness cell populations
Molecular signature expression
B
VAT
Muscle
Spleen
% of CD45
NK cells
T cells
NKT cells
GD T cells
iNKT cells
PBS
IL-15
C
Beta p-val<0.01
GZMB
IL2RB
IFNG
Innateness
Adaptiveness
D
Lymphocyte innateness (Gutierrez-Arcelus et al. 2019)
CD4+T CD8+T MAIT iNKT Vδ1 Vδ2 NK
TRBV29-1
TRBV19
NUAK2
TRAJ16
HLA-J
HLA-H
GTPBP1
TGFB1
MUL1
NAA50
A2MP1
CCL5
IFNG
CCL4
LINC00987
PZP
ETV3
BTG1P1
DUSP2
PPP2R5C
MRPL10
Scaled Expression
RPS6
RPS3A
RPS20
CAMK4
BCL11B
PTGDR
NCAM1
TYROBP
CTSD
HOPX
KLRF1
YPEL1
GNG2
UBB
PAFAH2
CLIC3
RHOC
APMAP
IL2RB
NMUR1
MCTP2
Scaled Expression
E
VAT-NK cells
IL2RB-innateness signature
-log p-value
FC IL15/control
IFNG
IL2RB
GZMB
CCL4
CCL5
F
VAT-NK cells
IFNG relative levels (ELISA)
IL2RB relative levels (ELISA)
control
IL15

fig. S10

**Fig. S10. DR regulates IL15-IL2RB and innateness signatures** *(related to Fig. 4).*
(**A**) Illustration of workflow overview.
(**B**) Percentage of lymphocytes in different tissues comparing control PBS and IL15 injections (intraperitoneal) at day 1 and day 4, followed by termination at day 7 (n=4-6).
(**C**) Top associated genes with innateness across lymphocyte human populations (scRNA-seq data and analysis from Gutierrez-Arcelus et al. (*63*)).
(**D**) As in C, scaled expression of innateness associated genes.
(**E**) From mice as in B, volcano plot expression of innateness genes.
**(F**) Relative IFNG and IL2RB levels by ELISA in VAT-NKs from mice as in B.
Animal experiments were done with n=4-6 mice per group. Data show mean values per group and SEM. Cell extraction experiments were done in each mice per group and mean per groups were compared. Unpaired, two-tailed student's t-test was used when two groups were compared, and ANOVA followed by fisher's least significant difference (LSD) test for post hoc comparisons for multiple groups. * p-value <0.05.

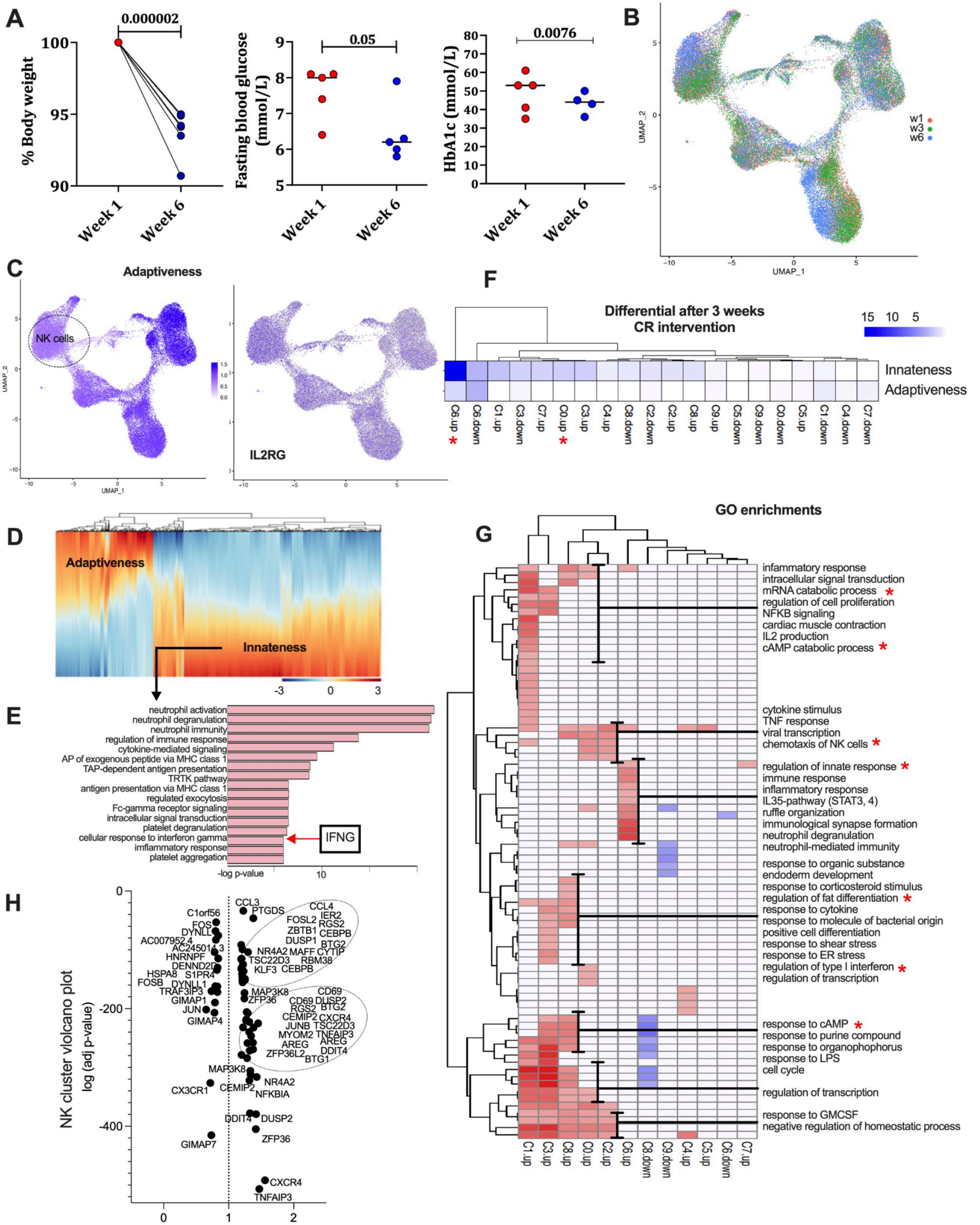
A
0.000002
% Body weight
Week 1
Week 6
0.05
Fasting blood glucose (mmol/L)
0.0076
HbA1c (mmol/L)
B
UMAP_1
UMAP_2
w1
w3
w6
C
Adaptiveness
NK cells
IL2RG
F
Differential after 3 weeks
CR intervention
Innateness
Adaptiveness
C6.up
C6.down
C1.up
C3.down
C7.up
C0.up
C3.up
C4.up
C8.down
C2.down
C2.up
C8.up
C9.up
C5.down
C9.down
C0.down
C5.up
C1.down
C4.down
C7.down
D
Adaptiveness
Innateness
E
neutrophil activation
neutrophil degranulation
neutrophil immunity
regulation of immune response
cytokine-mediated signaling
AP of exogenous peptide via MHC class 1
TAP-dependent antigen presentation
TRTK pathway
antigen presentation via MHC class 1
regulated exocytosis
Fc-gamma receptor signaling
intracellular signal transduction
platelet degranulation
cellular response to interferon gamma
imflammatory response
platelet aggregation
-log p-value
IFNG
GO enrichments
G
infammatory response
intracellular signal transduction
mRNA catabolic process *
regulation of cell proliferation
NFKB signaling
cardiac muscle contraction
IL2 production
cAMP catabolic process *
cytokine stimulus
TNF response
viral transcription
chemotaxis of NK cells *
regulation of innate response *
immune response
inflammatory response
IL35-pathway (STAT3, 4)
ruffle organization
immunological synapse formation
neutrophil degranulation
neutrophil-mediated immunity
response to organic substance
endoderm development
response to corticosteroid stimulus
regulation of fat differentiation *
response to cytokine
response to molecule of bacterial origin
positive cell differentiation
response to shear stress
response to ER stress
regulation of type I interferon *
regulation of transcription
response to cAMP *
response to purine compound
response to organophophorus
response to LPS
cell cycle
regulation of transcription
response to GMCSF
negative regulation of homeostatic process
C1.up
C3.up
C8.up
C0.up
C2.up
C6.up
C8.down
C9.down
C4.up
C5.up
C6.down
C7.up
H
NK cluster violcano plot
log (adj p-value)
Fold-change (week6/week0-3)
CCL3
PTGDS
CCL4
IER2
FOSL2
RGS2
ZBTB1
CEBPB
DUSP1
BTG2
NR4A2
MAFF
CYTIP
TSC22D3
RBM38
KLF3
CEBPB
C1orf56
FOS
DYNLL
HNRNPF
HSPA8
S1PR4
FOSB
DYNLL1
TRAF3IP3
GIMAP1
JUN
GIMAP4
MAP3K8
ZFP36
CD69
DUSP2
CEMIP2
CXCR4
JUNB
TSC22D3
MYOM2
TNFAIP3
AREG
DDIT4
ZFP36L2
BTG1
NR4A2
CX3CR1
NFKBIA
DUSP2
GIMAP7
TNFAIP3

**fig. S11**

**Fig. S11. DR regulates IL15-IL2RB and innateness signatures** *(related to Fig. 4).*
(**A**) Left panel, percentage of body weight in obese subjects at week 1 and 6 of a dietary weight loss intervention. Middle panel, fasting blood glucose. Right panel, glycosylated hemoglobin levels.
(**B**) Umap clustering of scRNA-seq data from lymphocyte cells from obese subjects under dietary intervention at week 0, 3, and 6.
(**C**) Left panel, enrichment of adaptiveness signatures (data from Gutierrez-Arcelus et al. (*63*)) on scRNA clusters from obese subjects (as in B). Right panel, Il2rg gene expression in scRNA umap clusters.
(**D**) Hierarchical clustering of innateness and adaptive signatures across lymphocyte scRNAseq data from obese subjects.
(**E**) Functional annotation of innateness signature genes from D, using the enrichR tool.
**(F**) Enrichment of innateness and adaptiveness signatures in differentially expressed genes across lymphocytes from obese subjects.
(**G**) Hierarchical clustering of gene-ontology enrichment in differentially expressed genes across lymphocytes.
(**H**) Volcano plot of differentially expressed genes after dietary restriction intervention in NK cells.
Data show mean values per group and SEM. Unpaired, two-tailed student's t-test was used when two groups were compared, and ANOVA followed by fisher's least significant difference (LSD) test for post hoc comparisons for multiple groups. * p-value <0.05.

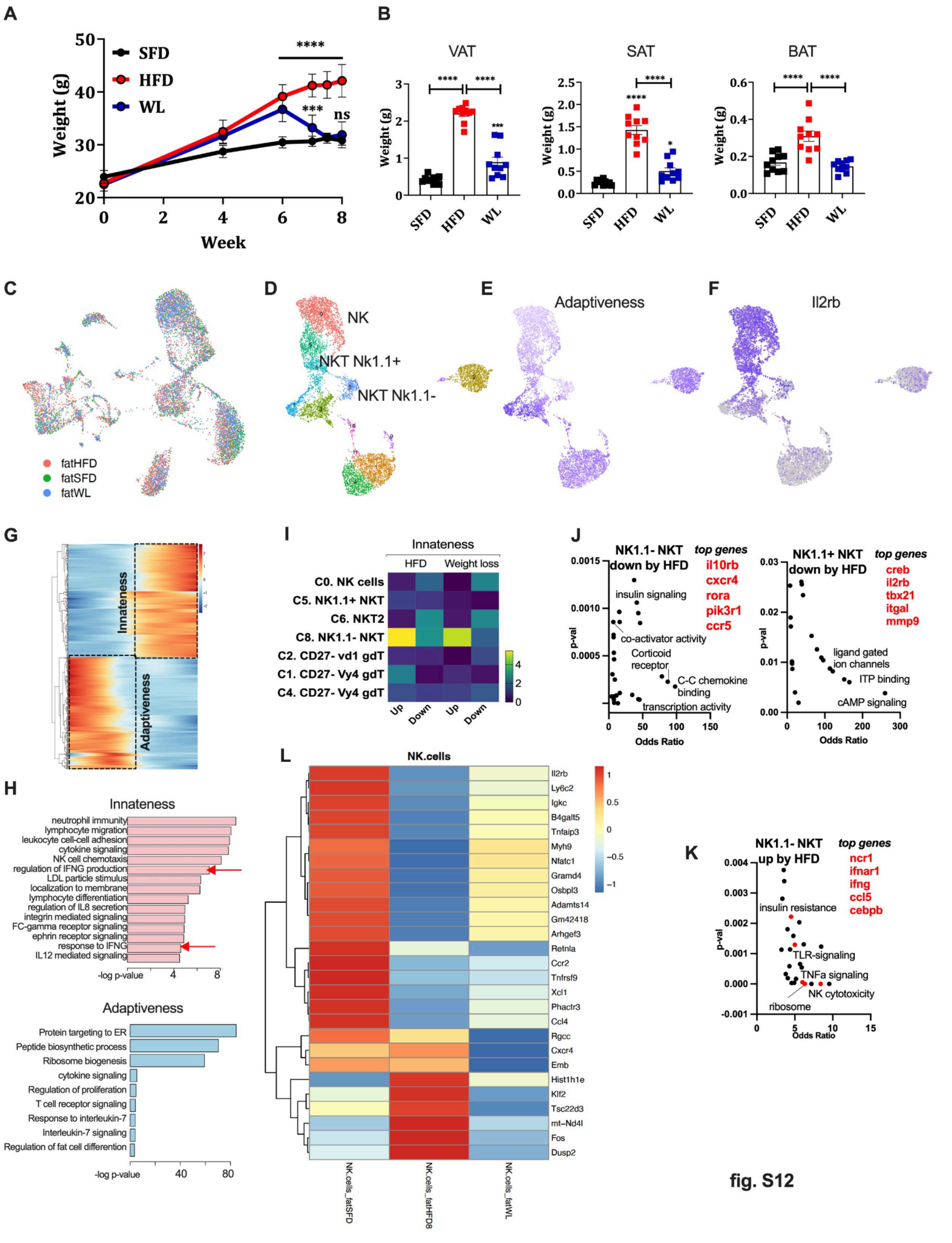

A
SFD
HFD
WL
Weight (g)
Week
****
***
ns
B
VAT
SAT
BAT
Weight (g)
SFD
HFD
WL
C
fatHFD
fatSFD
fatWL
D
NK
NKT Nk1.1+
NKT Nk1.1-
E
Adaptiveness
F
Il2rb
G
Innateness
Adaptiveness
H
Innateness
neutrophil immunity
lymphocyte migration
leukocyte cell-cell adhesion
cytokine signaling
NK cell chemotaxis
regulation of IFNG production
LDL particle stimulus
localization to membrane
lymphocyte differentiation
regulation of IL8 secretion
integrin mediated signaling
FC-gamma receptor signaling
ephrin receptor signaling
response to IFNG
IL12 mediated signaling
-log p-value
Adaptiveness
Protein targeting to ER
Peptide biosynthetic process
Ribosome biogenesis
cytokine signaling
Regulation of proliferation
T cell receptor signaling
Response to interleukin-7
Interleukin-7 signaling
Regulation of fat cell differention
-log p-value
I
Innateness
HFD
Weight loss
C0. NK cells
C5. NK1.1+ NKT
C6. NKT2
C8. NK1.1- NKT
C2. CD27- vd1 gdT
C1. CD27- Vy4 gdT
C4. CD27- Vy4 gdT
Up
Down
Up
Down
J
NK1.1- NKT down by HFD
top genes
il10rb
cxcr4
rora
pik3r1
ccr5
insulin signaling
co-activator activity
Corticoid receptor
C-C chemokine binding
transcription activity
p-val
Odds Ratio
NK1.1+ NKT down by HFD
top genes
creb
il2rb
tbx21
itgal
mmp9
ligand gated ion channels
ITP binding
cAMP signaling
p-val
Odds Ratio
K
NK1.1- NKT up by HFD
top genes
ncr1
ifnar1
ifng
ccl5
cebpb
insulin resistance
TLR-signaling
TNFa signaling
NK cytotoxicity
ribosome
p-val
Odds Ratio
L
NK.cells
Il2rb
Ly6c2
Igkc
B4galt5
Tnfaip3
Myh9
Nfatc1
Gramd4
Osbpl3
Adamts14
Gm42418
Arhgef3
Retnla
Ccr2
Tnfrsf9
Xcl1
Phactr3
Ccl4
Rgcc
Cxcr4
Emb
Hist1h1e
Klf2
Tsc22d3
mt−Nd4l
Fos
Dusp2
NK.cells_fatSFD
NK.cells_fatHFD8
NK.cells_fatWL


fig. S12

**Fig. S12. DR regulates IL15-IL2RB and innateness signatures** *(related to Fig. 4)*.
(**A**) Body weight (grams; g) evolution.
(**B**) Adipose depots mass (g) normalized by body weight.
(**C**) Umap clustering of scRNA-seq data from mice models under dietary intervention.
(**D**) Umap reclustering of scRNA-seq data for lymphocyte cells.
(**E**) Enrichment of adaptiveness signatures (data from Gutierrez-Arcelus et al. (*63*)) on scRNA clusters from obese subjects (as in B).
(**F**) Il2rb gene expression in scRNA umap lymphocyte clusters.
(**G**) Hierarchical clustering of innateness and adaptive signatures across lymphocyte scRNAseq murine data.
(**H**) Top panel, functional annotation of innateness signature genes from D, using the enrichR tool. Lower panel, functional annotation of adaptiveness signature genes.
(**I**) Enrichment of innateness and adaptiveness signatures in differentially expressed genes across lymphocytes.
(**J**) Functional biological enrichments using enrichR tools from differentially expressed genes across distinct lymphocyte populations.
(**K**) Functional biological enrichments using enrichR tools from differentially expressed genes across distinct lymphocyte populations.
(**L**) Hierarchical clustering of differentially expressed genes in NK cells
Animal experiments were done with n=4-6 mice per group. Data show mean values per group and SEM. Cell extraction experiments were done in each mice per group and mean per groups were compared. Unpaired, two-tailed student's t-test was used when two groups were compared, and ANOVA followed by fisher's least significant difference (LSD) test for post hoc comparisons for multiple groups. * p-value <0.05.

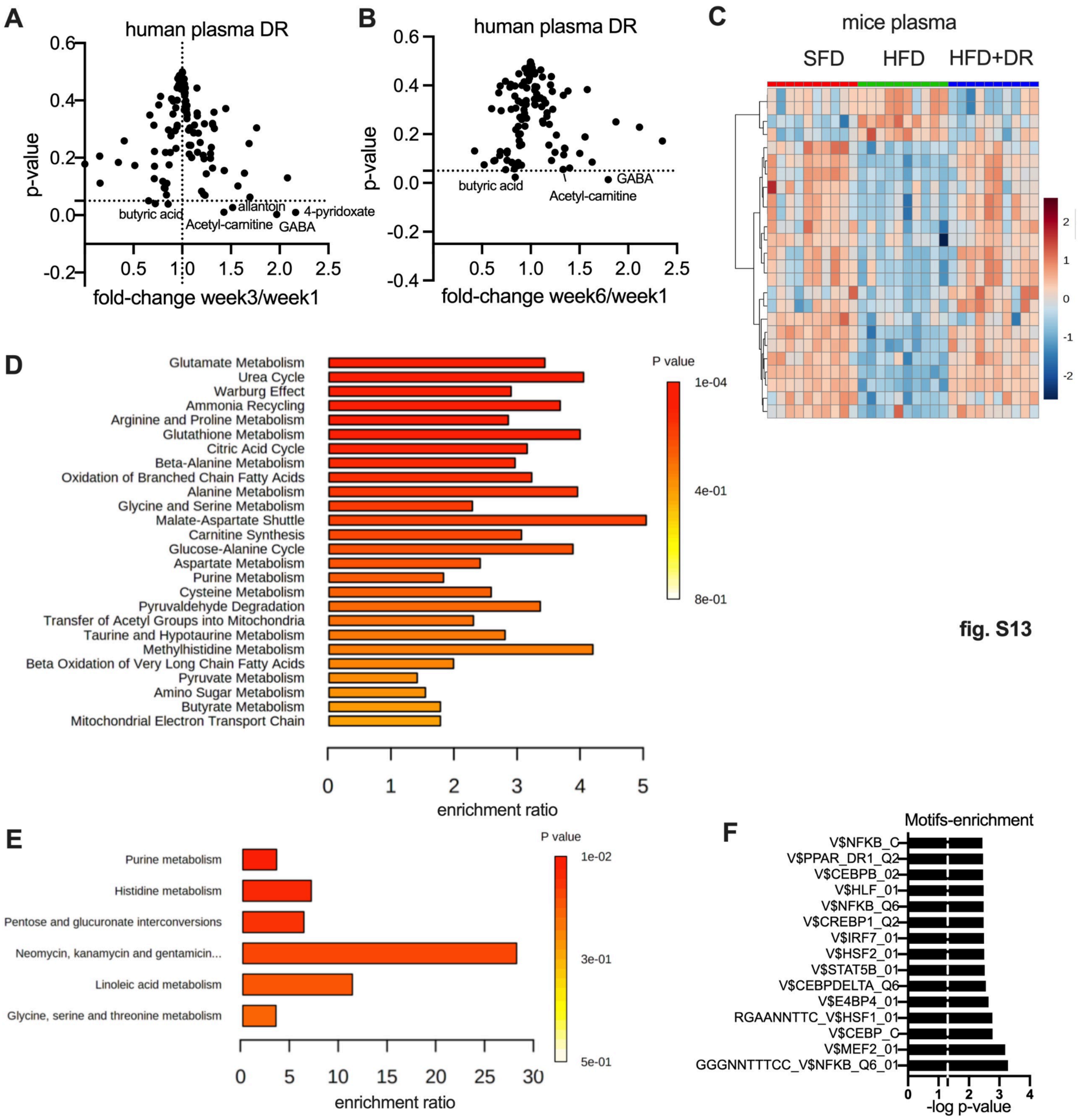


**Fig. S13. DR regulates IL15-IL2RB and innateness signatures** *(related to Fig. 5).*
(**A**) Volcano plots for human plasma metabolite variation after 3 weeks of dietary restriction.
(**B**) Volcano plots for human plasma metabolite variation after 6 weeks of dietary restriction.
(**C**) Hierarchical clustering for murine plasma metabolite variation comparing standard fat diet, HFD (6 weeks), and HFD followed by dietary restriction (2 weeks).
(**D**) Metabolic pathway enrichment analysis for plasma metabolites increasing after dietary restriction.
(**E**) Metabolic pathway enrichment analysis for VAT metabolites increasing after dietary restriction in mice.
(**F**) Motif enrichment analysis from multi-omic enriched pathways and associated genes (metaboanalyst tool).

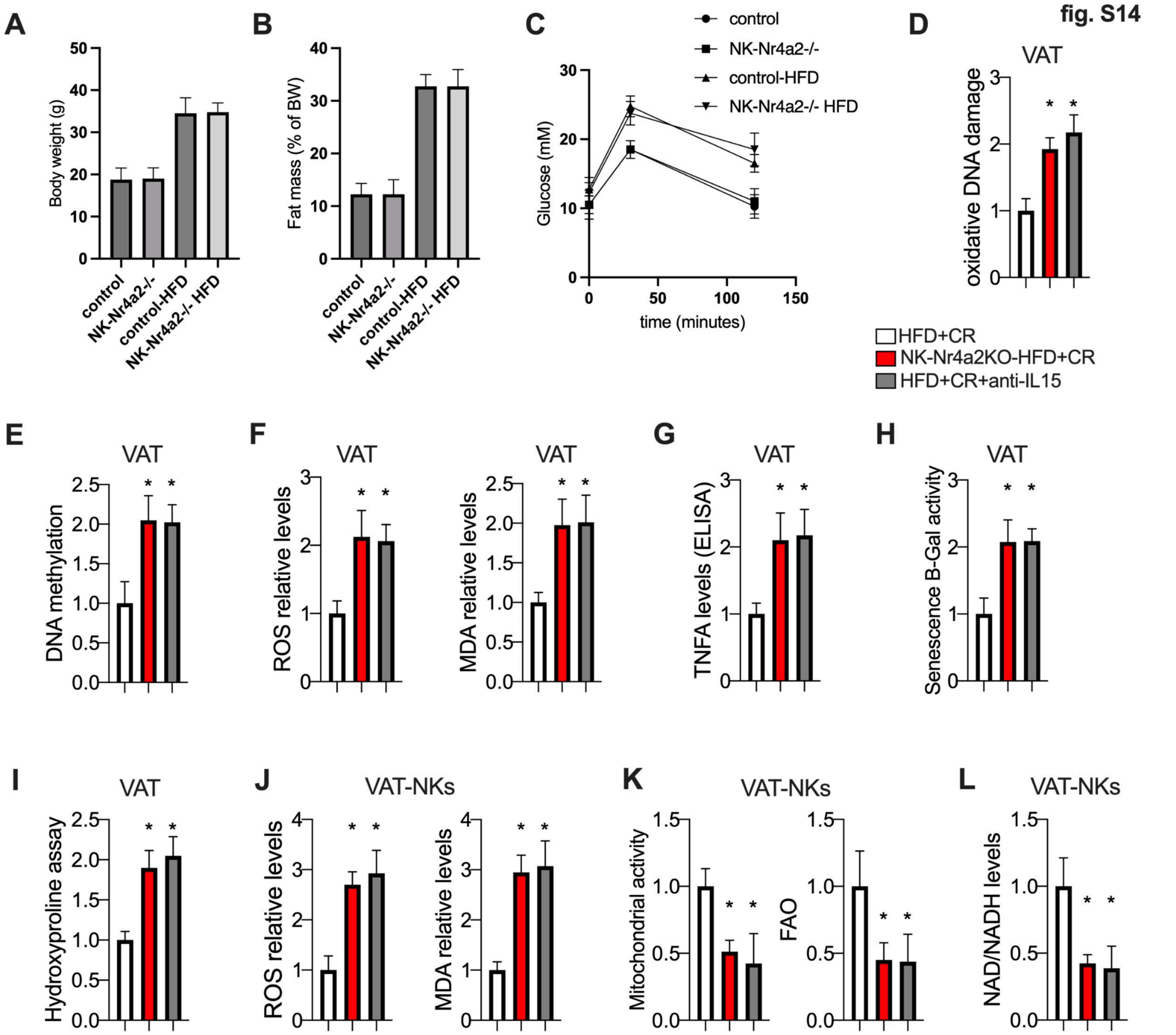


**Fig. S14. DR regulates IL15-IL2RB and innateness signatures** *(related to Fig. 5).*

(**A**) Body weight (grams; g) from indicated models.

(**B**) Fat mass percentage.

(**C**) Glucose tolerance test.

(**D**) Relative oxidative DNA damage in VAT.

(**E**) Relative DNA methylation in VAT.

(**F**) Left panel, reactive oxygen species (ROS) assay levels in VAT. Right panel, MDA relative levels (lipid peroxidation) in VAT.

(**G**) Relative TNFA levels by ELISA in VAT.

(**H**) Relative senescence B-gal assay activity in VAT.

(**I**) Relative hydroxyproline assay levels in VAT.

(**J**) Left panel, ROS assay activity in VAT-NK cells. Right panel, MDA relative levels in VAT-NK cells.

(**K**) Left panel, basal mitochondrial activity in VAT-NK cells. Right panel, FAO in VAT-NK cells (treated with FCCP 1 μM).

(**L**) Relative NAD+ levels in VAT-NK cells.

Animal experiments were done with n=4-6 mice per group. Data show mean values per group and SEM. Cell extraction experiments were done in each mice per group and mean per groups were compared when indicated. Unpaired, two-tailed student's t-test was used when two groups were compared, and ANOVA followed by fisher's least significant difference (LSD) test for post hoc comparisons for multiple groups. Two-way ANOVA was used to estimate significance between groups constrained by time measurements, and Tukey test for multiple comparisons * p-value <0.05.

fig. S15

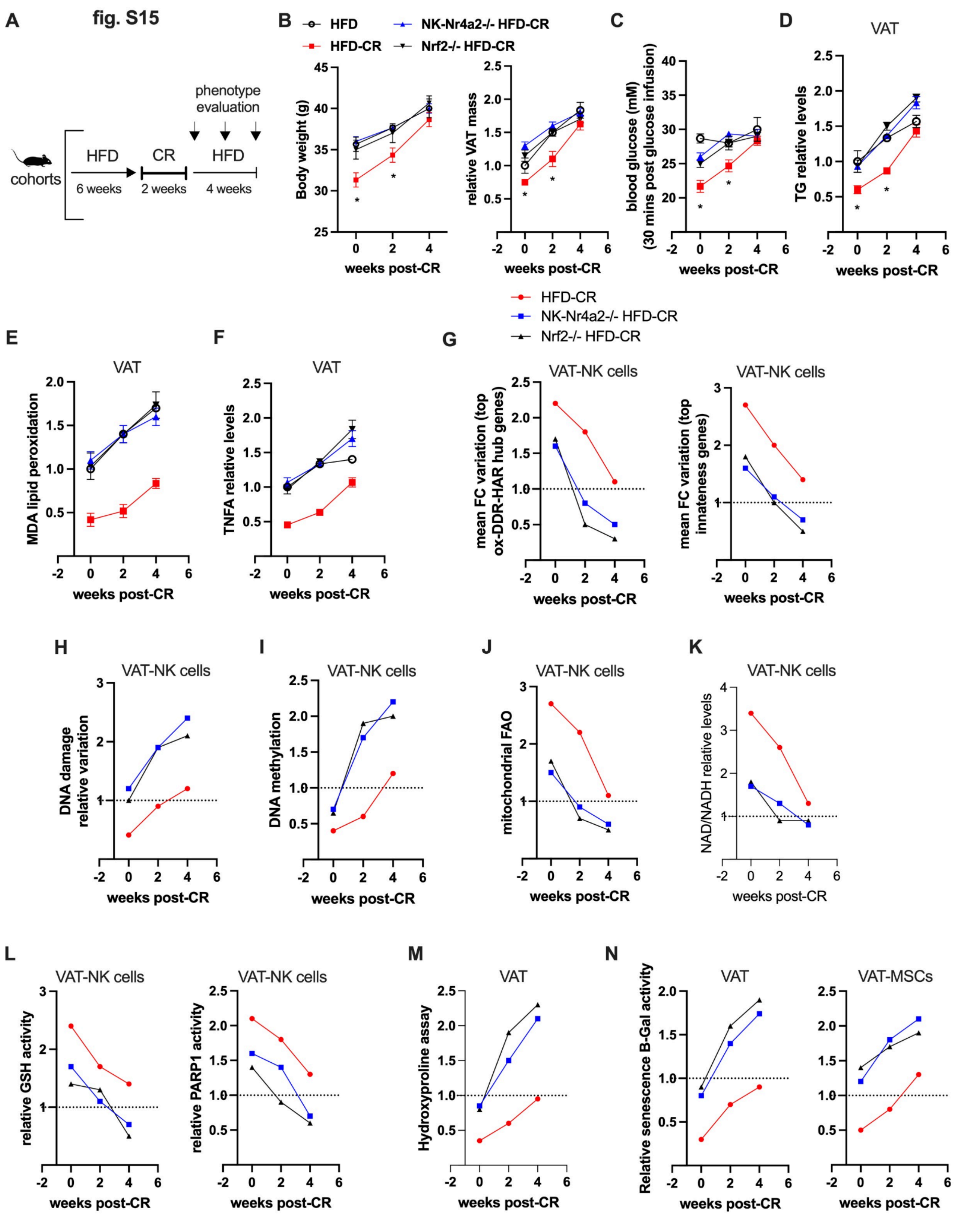
A
phenotype evaluation
HFD
CR
HFD
cohorts
6 weeks
2 weeks
4 weeks
B
HFD
HFD-CR
NK-Nr4a2-/- HFD-CR
Nrf2-/- HFD-CR
Body weight (g)
relative VAT mass
weeks post-CR
C
blood glucose (mM) (30 mins post glucose infusion)
D
VAT
TG relative levels
HFD-CR
NK-Nr4a2-/- HFD-CR
Nrf2-/- HFD-CR
E
VAT
MDA lipid peroxidation
F
VAT
TNFA relative levels
G
VAT-NK cells
mean FC variation (top ox-DDR-HAR hub genes)
mean FC variation (top innateness genes)
H
VAT-NK cells
DNA damage relative variation
I
DNA methylation
J
mitochondrial FAO
K
NAD/NADH relative levels
L
relative GSH activity
relative PARP1 activity
M
VAT
Hydroxyproline assay
N
VAT-MSCs
Relative senescence B-Gal activity

**Fig. S15. Dietary and evolutionary cofactors regulate epigenetic memory of resilience-enhancing NK cells**. *(related to Fig. 6).*
(**A**) Illustration of workflow overview.
(**B**) Body weight (grams; g) from indicated models.
(**B**) VAT mass relative to body weight.
(**C**) Blood glucose levels 30 mins post glucose infusion.
(**D**) Relative triglyceride (TGs) levels in VAT.
(**E**) Relative MDA levels (lipid peroxidation) in VAT.
(**F**) Relative TNFA levels in VAT.
(**G**) Left panel, module evaluation of fold-change (FC) variation across ox-DDR-HAR hub genes (n=10 genes) in VAT-NK cells from mice as in A. Right panel, module evaluation of fold-change (FC) variation across innateness genes (n=10 genes).
(**H**) Relative DNA damage in VAT-NK cells.
(**I**) Relative DNA methylation in VAT-NK cells.
(**J**) Relative mitochondrial FAO in VAT-NK cells.
(**K**) Relative NAD+ levels in VAT-NK cells.
(**L**) Left panel, relative glutathione activity. Right panel, relative PARP1 activity.
(**M**) Relative hydroxyproline assay activity in VAT.
(**N**) Relative senescence B-gal assay activity in VAT (left) and VAT-MSCs (right).
Animal experiments were done with n=4-6 mice per group. Data show mean values per group and SEM. Cell extraction experiments were done in each mice per group and mean per groups were compared when indicated. Unpaired, two-tailed student's t-test was used when two groups were compared, and ANOVA followed by fisher's least significant difference (LSD) test for post hoc comparisons for multiple groups. Two-way ANOVA was used to estimate significance between groups constrained by time measurements, and Tukey test for multiple comparisons * p-value $<0.05$.

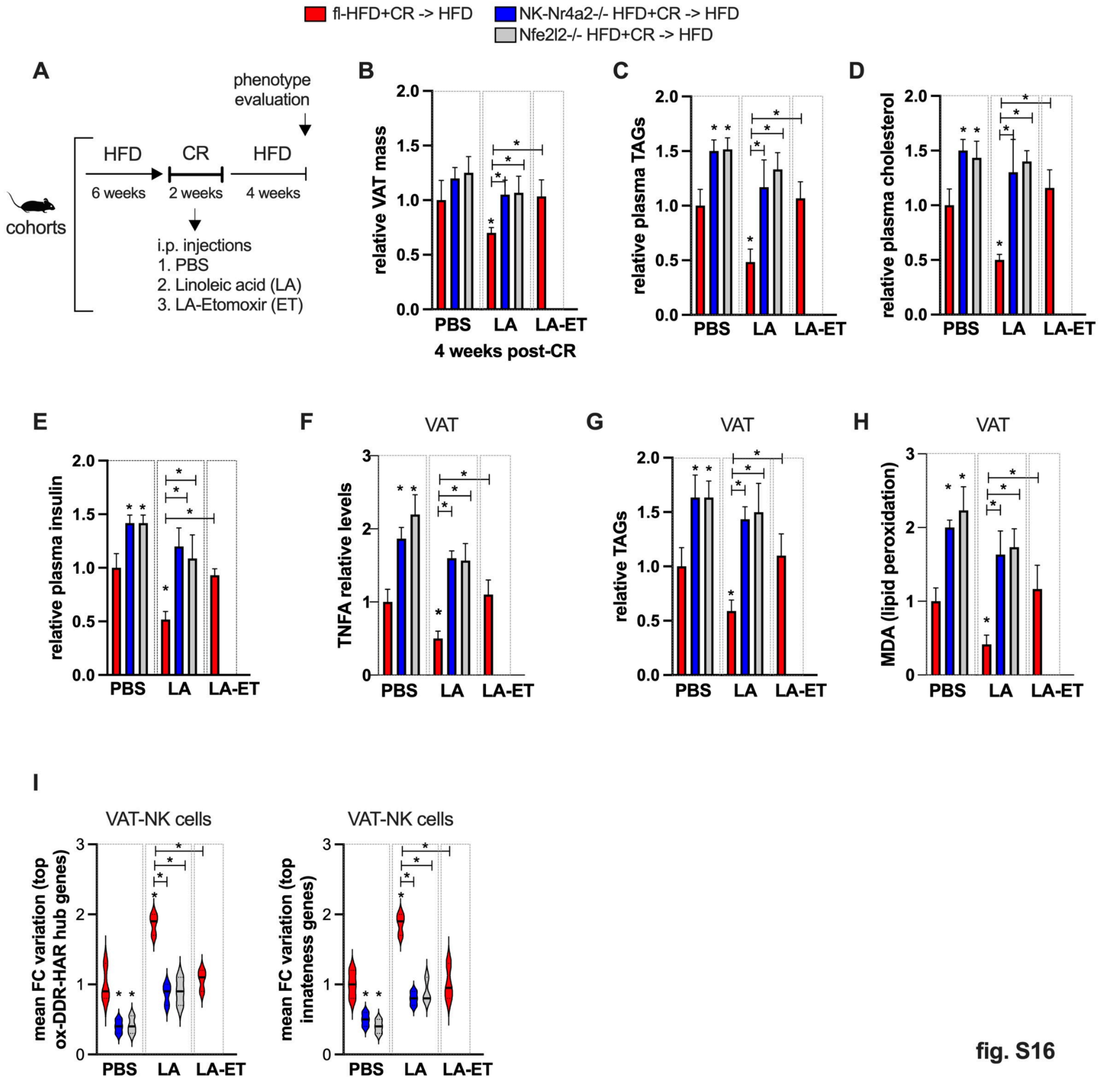


**Fig. S16. Dietary and evolutionary cofactors regulate epigenetic memory of resilience-enhancing NK cells**. *(related to Fig. 6).*
(**A**) Illustration of workflow overview.
(**B**) VAT mass relative to body weight.
(**C**) Relative triglyceride (TAGs) levels in plasma.
(**D**) Relative cholesterol levels in plasma.
(**E**) Relative insulin levels in plasma.
(**F**) Relative TNFA levels by ELISA in VAT.
(**G**) Relative TAGs levels in VAT.
(**H**) Relative MDA levels (lipid peroxidation) in VAT.
(**I**) Left panel, module evaluation of fold-change (FC) variation across ox-DDR-HAR hub genes (n=15 genes) in VAT-NK cells from mice as in A. Right panel, module evaluation of fold-change (FC) variation across innateness genes (n=15 genes).

Animal experiments were done with n=4-6 mice per group. Data show mean values per group and SEM. Cell extraction experiments were done in each mice per group and mean per groups were compared when indicated. Unpaired, two-tailed student's t-test was used when two groups were compared, and ANOVA followed by fisher's least significant difference (LSD) test for post hoc comparisons for multiple groups. Two-way ANOVA was used to estimate significance between groups constrained by time measurements, and Tukey test for multiple comparisons * p-value $<0.05$.

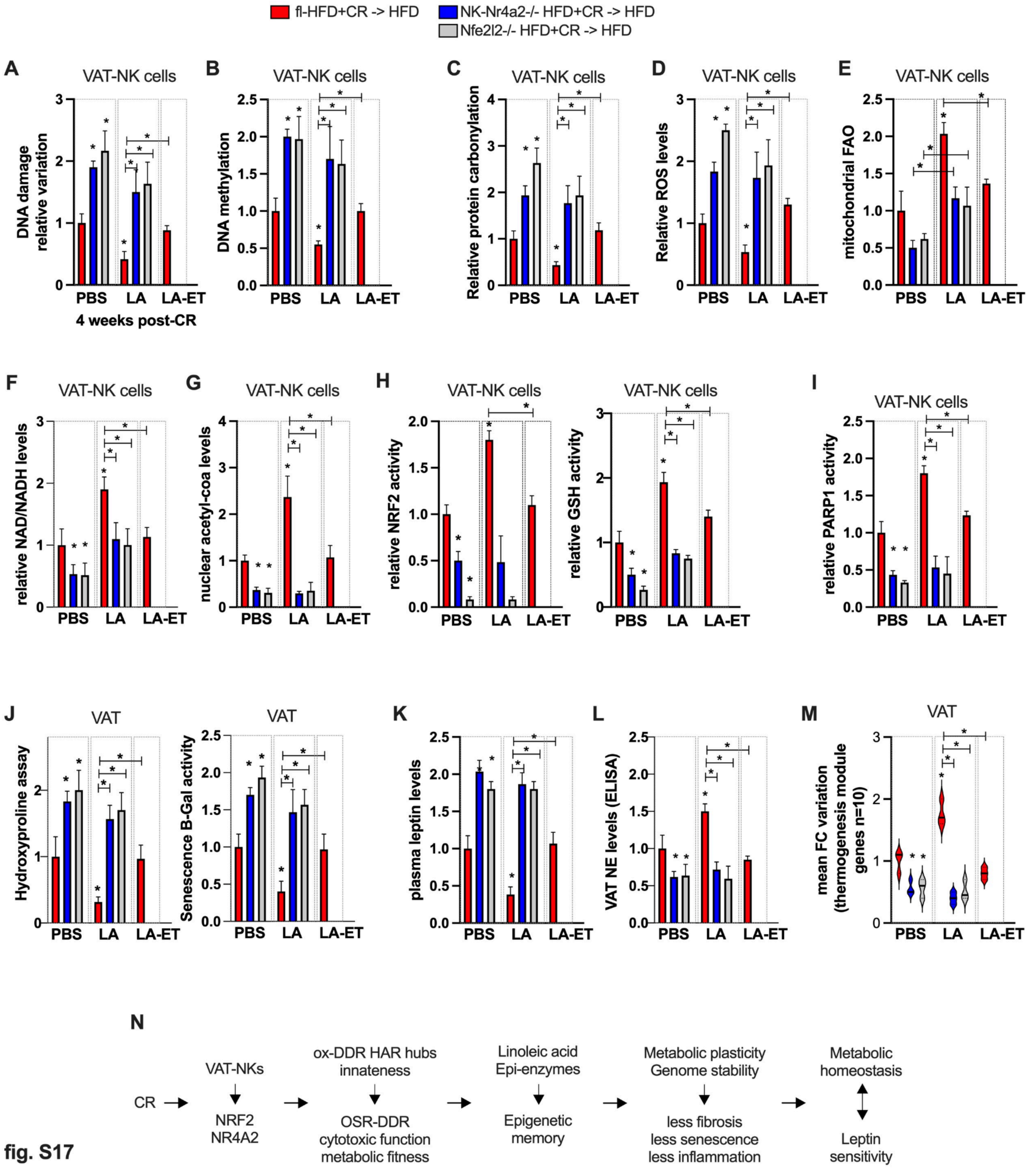


fig. S17

**Fig. S17. Dietary and evolutionary cofactors regulate epigenetic memory of resilience-enhancing NK cells** *(related to Fig. 6).*

(**A**) Relative DNA damage in VAT-NK cells from indicated mice models 4 weeks post-CR and reintroduced to a HFD.

(**B**) Relative DNA methylation in VAT-NK cells.

(**C**) Relative protein carbonylation levels in VAT-NK cells.

(**D**) Relative reactive oxygen species (ROS) assay levels in VAT-NK cells.
(**E**) Mitochondrial FAO in VAT-NK cells.
(**F**) Relative NAD+ levels in VAT-NK cells.
(**G**) Relative acetyl-coa levels in nuclear extracts from VAT-NK cells.
(**H**) Left panel, relative NRF2 activity in VAT-NK cells. Right panel, relative glutathione activity in VAT-NK cells.
(**I**) Relative PARP1 activity in VAT-NK cells.
(**J**) Left panel, relative hydroxyproline assay activity in VAT-NK cells. Right panel, Relative senescence B-gal assay activity.
(**K**) Relative plasma leptin levels.
(**L**) Relative norepinephrine (NE) levels in VAT.
(**M**) Module evaluation of fold-change (FC) variation across thermogenesis-associated genes (n=10 genes) in VAT.
(**N**) Summary illustration.
Animal experiments were done with n=4-6 mice per group. Data show mean values per group and SEM. Cell extraction experiments were done in each mice per group and mean per groups were compared. Unpaired, two-tailed student's t-test was used when two groups were compared, and ANOVA followed by fisher's least significant difference (LSD) test for post hoc comparisons for multiple groups. * p-value <0.05.

fig. S18

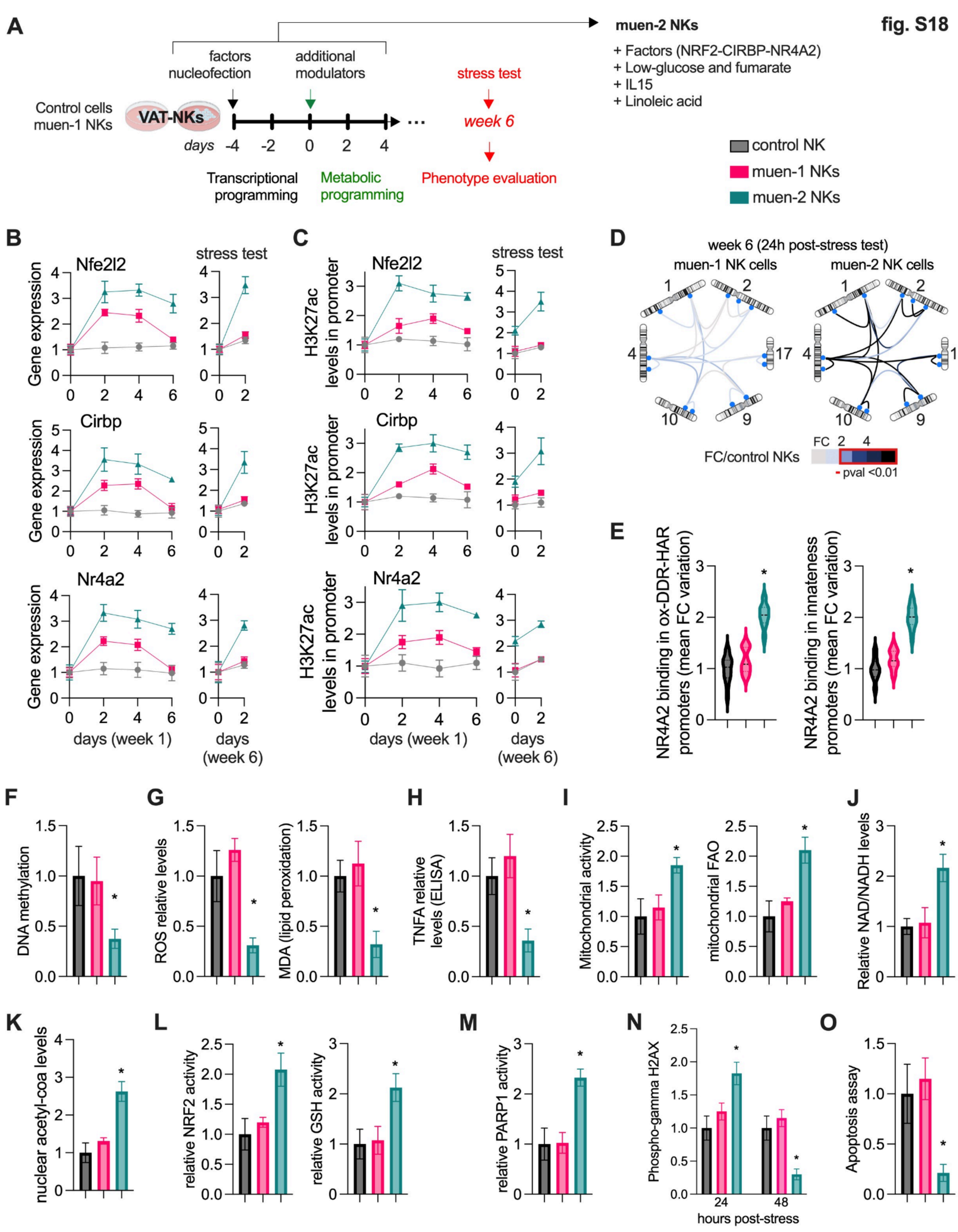

A
factors nucleofection
additional modulators
muen-2 NKs
+ Factors (NRF2-CIRBP-NR4A2)
+ Low-glucose and fumarate
+ IL15
+ Linoleic acid
Control cells muen-1 NKs
VAT-NKs
stress test
week 6
days -4 -2 0 2 4
Transcriptional programming
Metabolic programming
Phenotype evaluation
control NK
muen-1 NKs
muen-2 NKs
B
Nfe2l2
Cirbp
Nr4a2
stress test
Gene expression
days (week 1)
days (week 6)
C
Nfe2l2
Cirbp
Nr4a2
stress test
H3K27ac levels in promoter
days (week 1)
days (week 6)
D
week 6 (24h post-stress test)
muen-1 NK cells
muen-2 NK cells
FC/control NKs
FC 2 4
- pval <0.01
E
NR4A2 binding in ox-DDR-HAR promoters (mean FC variation)
NR4A2 binding in innateness promoters (mean FC variation)
F
DNA methylation
G
ROS relative levels
MDA (lipid peroxidation)
H
TNFA relative levels (ELISA)
I
Mitochondrial activity
mitochondrial FAO
J
Relative NAD/NADH levels
K
nuclear acetyl-coa levels
L
relative NRF2 activity
relative GSH activity
M
relative PARP1 activity
N
Phospho-gamma H2AX
24 48
hours post-stress
O
Apoptosis assay

**Fig. S18. Programming of resilience-promoting NK cells.** *(related to Fig. 7).*
(**A**) Illustration of workflow overview.
(**B**) Temporal gene expression in VAT-NK cells treated as indicated, and after an oxidative stress test at week 6 (hydrogen peroxide, 250 μM).
(**C**) Temporal H3K27ac levels (by ChIP) in promoters from cells as in B.
(**D**) *in situ* ChIP-loop chromosome conformation assays for chromatin contacts between top active ox-DDR-HAR genomic hubs in VAT-NK cells as in B. Targeted contact variations are represented as fold-change over the control group. Contacts with FC > 2 have p-value <0.01.
(**E**) Relative DNA methylation.
(**F**) Left panel, relative reactive oxygen species (ROS) assay levels. Right panel, relative MDA levels (lipid peroxidation).
(**G**) Relative TNFA levels by ELISA.
(**H**) Relative mitochondrial activity.
(**I**) Relative FAO.
(**J**) Relative NAD+ levels
(**K**) Relative acetyl-coa levels in nuclear extracts.
(**L**) Left panel, relative NRF2 activity. Right panel, relative glutathione activity.
(**M**) Relative PARP1 activity
(**N**) Relative H2AX phosphorylation assay by ELISA.
(**O**) Relative apoptosis assay.
Primary cell experiments were done with 3 independent replicates. Data show mean values per group and SEM. Cell extraction experiments were done in each mice per group and mean per groups were compared when indicated. Unpaired, two-tailed student's t-test was used when two groups were compared, and ANOVA followed by fisher's least significant difference (LSD) test for post hoc comparisons for multiple groups. * p-value <0.05.

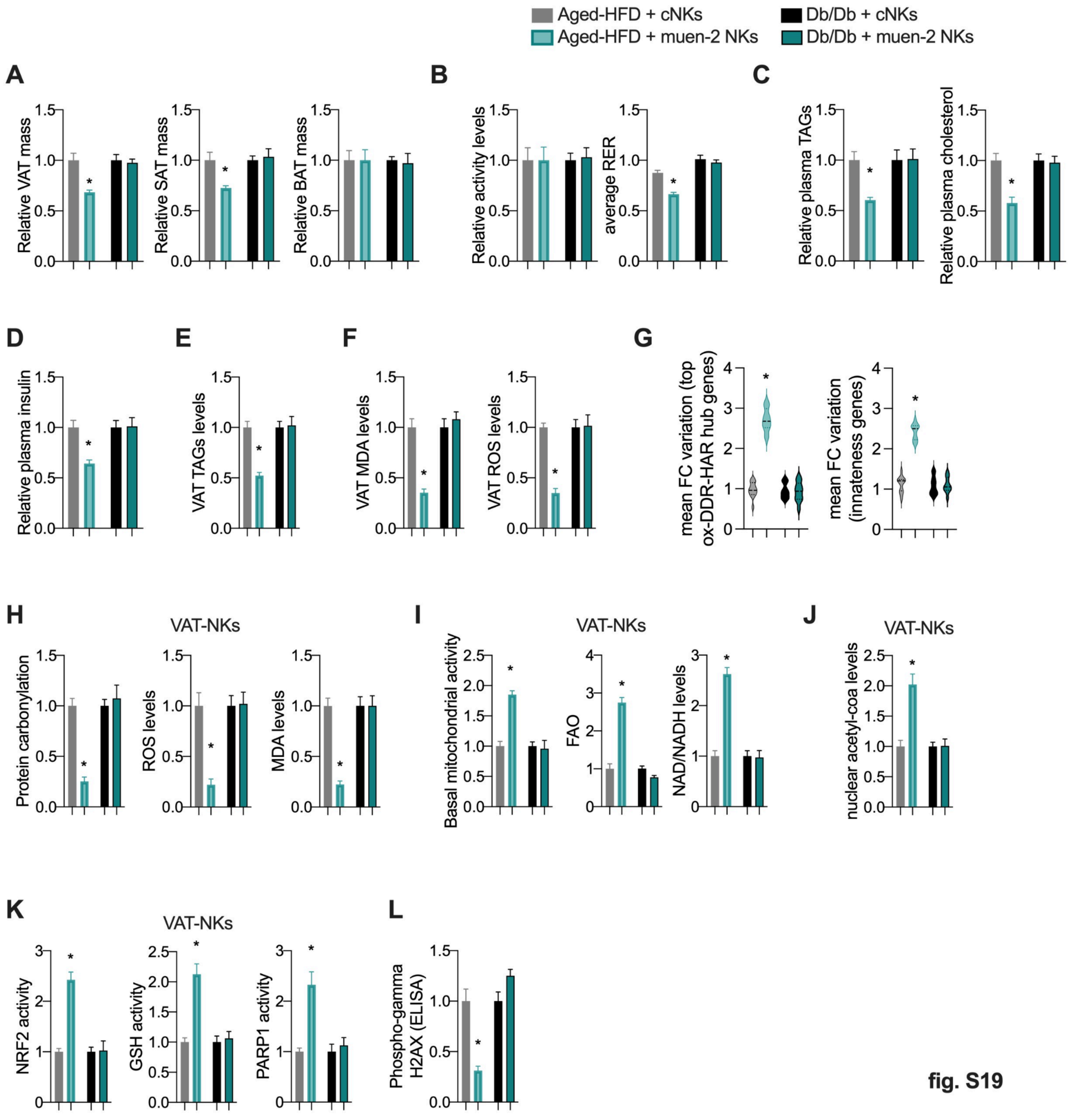


fig. S19

**Fig. S19. Resilience-promoting NK cells** *(related to Fig. 7).*
(**A**) Adipose depots mass relative to body weight in indicated mice models.
(**B**) Left panel, relative activity levels from 48 hours measurements in metabolic cages.. Right panel, average RER levels (night).
(**C**) Left panel, relative plasma triglyceride (TAGs) levels. Right panel, relative cholesterol levels.
(**D**) Relative insulin levels in plasma.
(**E**) Relative TAGs levels in plasma.
(**F**) Left panel, relative MDA levels (lipid peroxidation) in VAT. Right panel, relative reactive oxygen species (ROS) assay levels.

(**G**) Left panel, module evaluation of fold-change (FC) variation across ox-DDR-HAR hub genes (n=15 genes) in VAT-NK cells from mice as in A. Right panel, module evaluation of fold-change (FC) variation across innateness genes (n=15 genes).
(**H**) Left panel, relative protein carbonylation levels in VAT-NK cells from mice as in A. Middle panel, ROS relative levels. Left panel, MDA levels.
(**I**) Left panel, relative mitochondrial activity. Middle panel, FAO. Right panel, relative NAD+ levels.
(**J**) Relative nuclear acetyl-coa levels.
(**K**) Left panel, relative NRF2 activity. Middle panel, relative PARP1 activity. Right panel, relative glutathione activity
(**L**) Relative H2AX phosphorylation assay by ELISA
Bars show mean values and error bars indicate SEM. Unpaired, two-tailed student's t-test was used when two groups were compared, and ANOVA followed by fisher's least significant difference (LSD) test for post hoc comparisons for multiple groups.
Animal experiments were done with n=4-6 mice per group. Data show mean values per group and SEM. Cell extraction experiments were done in each mice per group and mean per groups were compared. Unpaired, two-tailed student's t-test was used when two groups were compared, and ANOVA followed by fisher's least significant difference (LSD) test for post hoc comparisons for multiple groups. * p-value <0.05.

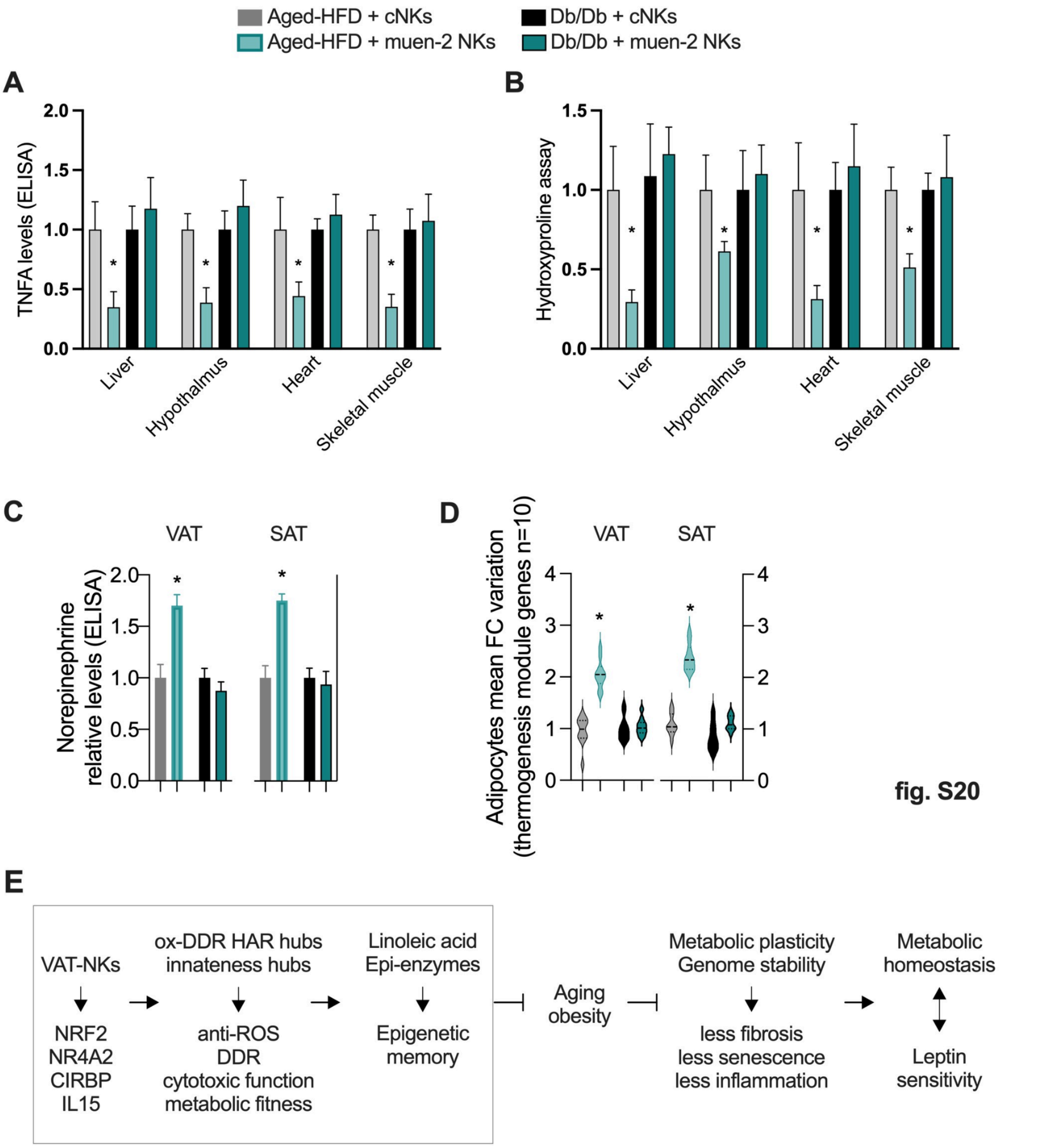


**Fig. S20. Resilience-promoting NK cells** *(related to Fig. 7).*

(**A**) Relative TNFA levels by ELISA across several tissues from indicated models.

(**B**) Relative hydroxyproline assay activity as in A.

(**C**) Relative norepinephrine (NE) levels in white adipose depots.

(**D**) Module evaluation of fold-change (FC) variation across thermogenesis-associated genes (n=10 genes) in white adipose depots.

(**E**) Summary illustration.

Animal experiments were done with n=4-6 mice per group. Data show mean values per group and SEM. Cell extraction experiments were done in each mice per group and mean per groups were compared. Unpaired, two-tailed student's t-test was used when two groups were compared, and ANOVA followed by fisher's least significant difference (LSD) test for post hoc comparisons for multiple groups. * p-value <0.05.

### 1. Supplementary tables

#### 1.1. Table S1. Metabolic phenotype comparison and human specific molecular signatures

1.1.A. Metabolic parameters and fasting endurance score in nonprimate mammals
1.1.B. Metabolic parameters and fasting endurance score in primate mammals
1.1.C. Metabolic parameters and fasting endurance score in hominoids
1.1.D. Tissue mass and metabolic parameters across mammals
1.1.E. GREAT analysis from ATACseq comparison between human-primate in fat: human increase
1.1.F. GREAT analysis from ATACseq comparison between human-primate in fat: human decrease
1.1.G. Fat branch-specific acceleration and positive selection from ATACseq comparison between humans and primates
1.1.H. Accelerated evolution signature genes in brain
1.1.I. List of human accelerated regions coordinates
1.1.J. HAR-associated genes discovery from Capra et al. 2013
1.1.K. Full list of HAR-genes compiled for this study
1.1.L. Functional annotation of HAR genes in brain-specific networks
1.1.M. Functional annotation of HAR genes in fat-specific networks
1.1.N. Data

#### 1.2. Table S2. Transcriptional regulatory networks of HAR-genes

1.2.A. ChIP-seq enrichment ChIP-atlas of HAR-genes
1.2.B. Adipocytes ChIP-seq enrichment ChIP-atlas of HAR-genes
1.2.C. Neural ChIP-seq enrichment ChIP-atlas of HAR-genes
1.2.D. Stem cells ChIP-seq enrichment ChIP-atlas of HAR-genes
1.2.E. Blood ChIP-seq enrichment ChIP-atlas of HAR-genes
1.2.F. Muscle ChIP-seq enrichment ChIP-atlas of HAR-genes
1.2.G. Liver ChIP-seq enrichment ChIP-atlas of HAR-genes
1.2.H. TFBS enrichment from MSDB
1.2.I. TFBS enrichment form genome browser
1.2.J. ENCODE and ChEA consensus enrichment
1.2.K. Data

#### 1.3. Table S3. Transcriptional regulatory networks, and PPI reconstitution

1.3.A. Node + 1 network reconstruction of PPIs for HAR-genes regulators
1.3.B. Network topological prioritization of network in A
1.3.C. Full PGC1A-network reconstruction of protein-protein interactions
1.3.D. Network topological prioritization of full PGC1A-network
1.3.E. Reactome functional annotation of HAR cooperative regulators
1.3.F. Gene-ontology functional annotation of HAR cooperative regulators
1.3.G. Fat and brain tissue-specific PPIs for PGC1A-network
1.3.H. Data

#### 1.4. Table S4. Transcriptional regulation of HAR-genes and metabolic-HAR hubs

1.4.A. Functionally-associated and interacting transcriptional regulators: co-regulated genes by promoter ChIP-seq enrichment
1.4.B. Co-bound transcriptional regulators: co-regulated HAR-genes (metabolic-HAR genes)
1.4.C. Positional gene enrichments of metabolic-HAR genes
1.4.D. Cytogenetic band enrichments of metabolic-HAR genes
1.4.E. Functional annotation of metabolic-HAR PGE-domains
1.4.F. Data

### Materials and Methods

#### Animal experiments

All animal experiments were conducted in accordance with internationally accepted guidelines for the use of laboratory animals and were approved by multiple institutional and regional committees. These include the Institutional Animal Care and Use Committees (IACUC) of Novo Nordisk Research Center Seattle, Novo Nordisk Research Center Lexington, Massachusetts; the Novo Nordisk Ethical Review Committees in China; MIT committee of animal care; the Institutional Animal Care and Use Committee guidelines of The Dana-Farber Cancer Institute and Harvard Medical School; the Trinity College Dublin University Ethics Committee and the Health Products Regulatory Authority in Ireland; the regional ethics committee of Alicante and Universidad Miguel Hernández-CSIC, Spain; and the Institutional Animal Care and Use Committee of Brigham and Women's Hospital. C57BL/6J mice aged 6-12 weeks or 18 months (Jackson Laboratories, Stock # 000664). High fat diet: At 6 weeks of age wild-type mice or 18 months mice were placed on a 60% high fat diet (High-Fat Diet #D12492 (20.0% Protein, 60% Fat, 20% Carbohydrate) during indicated times in weeks. Caloric restriction: Indicated mouse models followed a reduction in calorie intake to 60% of the ad-libitum control food intake, resulting in approximately 40% fewer calories. This was carried out for 2 weeks. Dietary restriction: indicated mouse models after a HFD intervention (6 weeks) are reintroduced to standard calorie intake diet for 2 weeks. HFD-reintroduction: following a CR intervention, mouse models when indicated are reintroduced to HFD for a maximum of 4 weeks. We used Nrf2 KO mice (Jax, stock 017009). Cirbp deficient mice generated by homologous recombination using the targeting vector pKO-Cirbp and ES cells, followed by obtaining chimeric mice. These mice were mated with C57BL/6 mice to obtain cirbp+/- mice and backcrossed until obtaining KO mice (*89*). Ncr1-cre mice were generated as previously demonstrated (*90*), In brief, cre was inserted into a bacterial artificial chromosome (BAC, RP23-267N11, Children's hospital oakland research institute, California), with the Ncr1 gene. Recombined BAC, containing the cre recombinase, were used to generate chimeric mice, followed by mating and backcrossing with C57BL/6 mice. Floxed Nr4a2 mice were generated by homologous recombination as previously described (*91*). In brief, mouse embryonic cells were electroporated with the targeting vector containing the Nr4a2 gene flanked by loxp site in intron 2 (*91*), followed by chimeric mice generation and backcrossing with C57BL/6 mice. These mice were mated with the Ncr1 cre line to generate NK cell deletion of the Nr4a2 gene. We also used Db/Db leptin receptor deficient mice (Jax, stock 000697). Mice protocols and treatments: intraperitoneal daily injections of linoleic acid in solution (10 mg/kg) and in combination with etomoxir (10 mg/kg). Anti-IL15 antibodies (eBioscience) were used as previously reported (*92*). Briefly, mouse models were treated with neutralizing anti-IL15 antibodies (5ug ip daily in 100 ul of sterile saline) for the duration of the CR intervention. Mouse models were also given IP injection of recombinant IL15 during a week (daily, 2,5 ug, Peprotech).

#### Mammalian cell isolation and culture

Human peripheral blood mononuclear cells (PBMCs) were isolated from obese subjects volunteer donors undergoing dietary restriction intervention by ficoll density gradient separation using ficoll-paque plus (GE Healthcare), and residual RBCs were removed by adding red cell lysis solution (ACK Lysing Buffer, VWR). Human PBMCs were used for downstream single-cell sequencing (see protocol). Adipose tissue was excised, finely minced using a razor, and digested in 1 mg/ml Collagenase Type II (Worthington) in RPMI medium while shaking for 25 minutes at 37°C. The digested cell suspension was then filtered through a 70 µm nylon mesh and centrifuged at 15,000 rpm for 7 minutes to pellet the stromal vascular fraction (SVF) for further processing.

Adipocyte Progenitors and Mesenchymal Stem Cells (MSCs) from VAT: To isolate adipose progenitors from adipose tissues, the Adipose Tissue Dissociation Kit (Miltenyi Biotec) and the Adipose Tissue Progenitor Isolation Kit (Miltenyi Biotec) were used following the manufacturer's instructions.

Adipose Tissue Mesenchymal Stem Cells (AT-MSCs): To isolate AT-MSCs, the CD271 MicroBead Kit (Miltenyi Biotec) was used following manufacturer's instructions.

Isolation of Adipocytes was performed during the stromal vascular fraction (SVF) extraction with modifications during the pelleting steps. After incubating the tissue in a vigorously shaking water bath for 20-30 minutes at 37°C, cells disintegrate into individual components. Centrifugation was performed at 500g (2200 RPM) for 10 minutes to separate the floating adipocytes from the pelleted SVF. The floating adipocyte layer was carefully aspirated using a cut 1000 µL pipette tip and transferred to a fresh tube containing a warm cell buffer (for further experiments) or SDS-lysis buffer (for analysis).

Primary NK cells were purified by positive selection using the NK cell purification kit (Miltenyi Biotec) from visceral adipose tissue of indicated mouse models, and using the magnetic-activated cell sorting (MACS). NK cells were selected based on positive expression of NK1.1. NK cells were then cultured with RPMI 1640 (Life Technologies) supplemented with 10% heat-inactivated fetal bovine serum (Life Technologies), 2 mM L-glutamine (Gibco™), 50 µM β-mercaptoethanol (Sigma-Aldrich), 100 U/ml penicillin, and 100 µg/ml streptomycin (Gibco™), and recombinant mouse IL-2 (PeproTech) at

1000 U/ml. Maintenance: Media were refreshed every 2-3 days, with IL-2 supplementation, and cells were passaged at 80% confluence by diluting them in fresh medium.

Cd11b+ monocyte/macrophage cells, Cd11c+ dendritic cells, and pan-T cells from VAT: After tissue dissociation, samples are processed as described below following magnetic-activated cell sorting strategies. Monocytes/macrophages are isolated by the digestion with pronase and collagenase digestion followed by discontinuous gradient ultracentrifugation. After the first centrifugation step at 600 rpm, the supernatant containing non-parenchymal cells (which include immune cells) is carefully collected. This supernatant is further centrifuged at 1500 rpm for 5 minutes to obtain a pellet enriched in immune cells, including monocytes and macrophages. The cell pellet is washed with DMEM medium and centrifuged again. After washing, the pellet is resuspended in 90 µl of MACS buffer (1x PBS, 0.5% BSA, 2 mM EDTA) with 0.6% citrate-dextrose solution (Sigma-Aldrich) to prevent cell aggregation. CD11b Magnetic Labeling: Add 10 µl of CD11b MicroBeads (Miltenyi Biotec, #130-049-601) to the cell suspension and incubate for 20 minutes at 4°C. Ensure the cells are mixed gently during the incubation to allow for efficient labeling. Magnetic-Activated Cell Sorting (MACS): Prepare a MACS LS column (Miltenyi Biotec, #130-042-401) by placing it on a MACS MultiStand and washing it with 3 ml of PBS followed by 3 ml of MACS buffer. After incubation, pass the cell suspension through the MACS column to remove CD11b-negative cells. Wash the column with 5 ml of MACS buffer to collect the CD11b-positive monocytes/macrophages. The CD11b+ cells are now isolated and ready for further analysis or experimentation. For isolation of DCs, we use pan-DC microbeads consisting of CD11c and Anti-mPDCA-1 MicroBeads (Miltenyi Biotec, #130-092-465) following manufacturer's instructions. For isolation of pan T cells, we use pan-T microbeads (Miltenyi Biotec, #130-095-130) following manufacturer's instructions.

NK cell treatments: Indicated cell cultures were kept in different condition media as described: Dulbecco's media with low glucose (2mM) when indicated. For RNAi screens cells were transfected using lipofectamine 3000 (Invitrogen, Thermo fisher technologies) with siRNA for transcriptional regulators, nfe2l2, cirbp, tead4, foxm1, and klf7, along with scramble control (Applied biosystems). When indicated, cells were treated with palmitate, (50 µM, Tocris), fumarate (50 µM, Tocris), etomoxir (50µM, Sigma Aldrich), hydrogen peroxide (250 µM, Tocris), IL15 (20 ng/ml, Peprotech), linoleic acid (100 µM, Sigma).

Plasmids and transfection: Plasmids were transfected into NK cells using the Amaxa Nucleofector and Nucleofector Kits (Lonza) for primary cell line transfection, following the manufacturer's protocols. Cells were transfected when they reached 70-80% confluence after 1-3 passages. For co-transfection of multiple plasmids, plasmid DNAs were pre-mixed in equal concentrations prior to transfection. siRNA molecules were transfected using Lipofectamine 3000 (Invitrogen, Thermo Fisher Scientific) and removed 24 hours post-transfection, in accordance with the manufacturer's protocols. After transfection, cells were subjected to protein identification and other functional assays as indicated. Expression vector for Nfe2l2 (NM_010902) mouse Tagged Clone
(Origene, CAT#: MR226717), Cirbp (NM_007705) Mouse Tagged Clone (Origene, CAT#: MR201471), and Nr4a2 (NM_001139509) Mouse Tagged Clone (Origene, CAT#: MR226282).

NK cell engineering: VAT-NK cells from mice models were isolated and maintained in vitro for 3 days in 12 well plates to cell density of 2x10e5 cell/well. On day 3 cells were dissociated with Trypsin (Sigma aldrich) and resuspended in solution. Cells were transfected in suspension with the Amaxa-nucleofector system (Lonza) and recommended primary cell nucleofector kits (Lonza). When indicated, cells were transfected with expressing plasmid vectors for NRF2, CIRBP, NR4A2 individually or in combination. When indicated, media was changed to low glucose media (2mM). Protein variation and transfection efficiency were tested using proximity ligation assays as indicated (see PLA assays) and gene expression. When indicated, cells were maintained 4-12 passages, with passaging (sub-culturing) days every 3rd day, where the cells were resuspended, counted and plated. Control cells were maintained as indicated and were transfected with empty expression vectors. Molecular and functional assays were performed in modified and control cells.

**Analysis of Gene Expression**

Total RNA was isolated from cells or tissues using Isol-RNA Lysis Reagent (5 PRIME) according to the manufacturer's instructions. To remove any contaminating DNA, Amplification Grade DNase I (Life Technologies) was used to treat 1 µg of RNA, after which 500 ng of the treated RNA was used for cDNA preparation following the Applied Biosystem Reverse Transcription Kit (Life Technologies) protocol.

Quantitative Real-Time PCR (qPCR) was performed using a ViiA 7 and QuantStudio Real-Time PCR system thermal cycler with SYBR Green PCR Master Mix (both Applied Biosystems). Gene expression analysis was conducted using the ΔΔCt method, with relative expression levels normalized to hypoxanthine phosphoribosyltransferase (HPRT) mRNA as a reference gene. When necessary, gene expression data were presented as mRNA levels relative to control samples. To assess the transcriptional activity of genes located within genomic hubs (such as OSR-DDR-HAR hub genes) or functionally similar molecular signatures (e.g., pro-fibrotic genes, SASP genes, thermogenesis genes, leptin-induced genes), the mean fold-change variation over control for each gene was calculated. These results were then compared across different biological conditions and displayed as a module to visualize the overall effects. Primer sequences are available upon reasonable request.

**Chromatin immunoprecipitation**

Cells, ranging from 2 to 10 million, were plated and pooled before homogenization, while tissue samples, approximately 50 mg, were homogenized in PBS. Both cells and tissues were crosslinked with 1% formaldehyde for 10 minutes to stabilize protein-DNA interactions. The reaction was quenched by adding 125 mM glycine, followed by centrifugation at 4000 rpm for 5 minutes. After removing the supernatant, the pellet was washed with 1 ml of cold PBS and centrifuged again at 4000 rpm for 5 minutes. This washing step was repeated twice: first using buffer 1 (containing 0.25% Triton X-100, 10 mM EDTA, 0.5 mM EGTA, and 10 mM HEPES, pH 6.5), and then buffer 2 (composed of 200 mM NaCl, 1 mM EDTA, 0.5 mM EGTA, and 10 mM HEPES, pH 6.5). Following the washes, the pellet was resuspended in 300 µl of lysis buffer (1% SDS, 10 mM EDTA, 50 mM Tris-HCl, pH 8) containing protease inhibitors. Samples were sonicated on ice using a Bioruptor (Diagenode) for 30 cycles of 30 seconds ON and OFF intervals, shearing the DNA into fragments approximately 300 bp in length. After sonication, specific antibodies were added to the lysates, with 10-20 µl of each antibody used for immunoprecipitation. Negative controls without antibodies were included in each ChIP assay, and no precipitation was detected by quantitative Real-Time PCR (qPCR) analysis. Input samples were processed in parallel to the immunoprecipitated samples. To isolate antibody/protein/DNA complexes, the samples were incubated with either salmon sperm DNA/protein A agarose slurry or Protein A/G PLUS agarose beads (Santa Cruz sc-2003), followed by several washes to ensure purity. The immune complexes were then eluted using a solution of 1% SDS and 0.1 M NaHCO3. The samples were treated with proteinase K for 1 hour to degrade proteins, and the DNA was purified by phenol/chloroform extraction, followed by ethanol precipitation. Finally, the DNA was resuspended in 20 µl of H2O. Both input DNA and immunoprecipitated DNA were analyzed by quantitative PCR (qPCR). Primer sequences used to detect the targeted DNA regions are available upon request. To evaluate epigenetic activity and H3K27ac levels in transcriptional compartment hub genes (such as OSR-DDR-HAR hub genes), fold-change variations over control were calculated for each promoter region and presented as a module to compare the overall effects under different biological conditions.

**3C Chromosome conformation assays**

Between 15,000 and 2 million cells were used for in situ chromosome conformation capture (3C) assays, following previously established methods (*93*) with some modifications. Briefly, cells were crosslinked with 1% formaldehyde for 10-15 minutes at room temperature to preserve chromatin interactions. Cells were then lysed using a buffer containing 60 mM Tris (pH 7.5), 0.5% Igepal, 0.25% sodium deoxycholate, 0.1% SDS, 150 mM NaCl, and protease inhibitors. The pelleted nuclei were resuspended in 0.4% SDS and incubated at 60°C for 30 minutes to permeabilize the chromatin. After quenching with 1% Triton-X for 12 minutes at 37°C, nuclei were digested with 5 U/µl DpnII in 1x DpnII buffer overnight at 37°C. Following digestion, the cells were pelleted (2500g for 10 minutes), and the restriction enzyme was inactivated at 65°C for 20 minutes. The ends of the DNA fragments were filled in with biotinylated dATP during a 1.5-hour incubation at 37°C, and ligation was carried out at 25°C for 4 hours with rotation. The ligated nuclei were then pelleted and sonicated in a buffer containing 10 mM Tris (pH 7.5), 1 mM EDTA, and 0.25% SDS for 16 minutes, using a sonicator set to 100 intensity, 200 cycles per burst, and a maximum temperature of 5°C. The DNA was reverse cross-linked overnight at 65°C in the presence of proteinase K and RNase A. After reverse cross-linking, the ligated DNA was purified and sheared to approximately 250-500 base pairs. Biotinylated ligation junctions were pulled down using streptavidin beads and prepared for targeted quantitative measurement of chromatin interactions using qPCR assays to compare different biological samples. Results were represented as fold-change of interaction over control conditions, with statistical significance and fold-variations shown as indicated in figure legends. The top active genes in each hub were used for this analysis, followed by targeted 3C evaluation of promoter-promoter interactions, guided by GeneHancer or NCBI promoter coordinates. Primer sequences are available upon reasonable request.

**Proximity ligation assays**

Cell and tissue lysates were prepared using RIPA buffer (50 mM Tris pH 7.5, 150 mM NaCl, 1 mM EDTA, 1% (w/v) Triton-X-100, 0.5% (w/v) sodium deoxycholate, 0.1% (w/v) sodium dodecyl sulfate, 20 mM glycerol-2-phosphate, 5 mM sodium pyrophosphate) freshly supplemented with 1 mM DTT and 0.5 mM PMSF. Antibodies were conjugated with oligonucleotides using the Oligonucleotide Conjugation Kit for Proximity Ligation Assays (PLA, ab218260). The protein concentration was determined using the Bradford Protein Assay following the manufacturer's instructions (BioRad). Protein extracts and conjugated antibodies were used in PLA assays, where qPCR on conjugated products was performed according to the manufacturer's instructions. For the homogeneous PLA assay (individual proteins), antibodies specific to tag proteins (such as Myc tags) and proteins of interest (NRF2 and CIRBP) were conjugated with 3' or 5' oligonucleotide probes, respectively.

This was followed by immuno-qPCR in accordance with the manufacturer's protocol for detecting protein-protein interactions.

**Extracellular Flux Analysis (Seahorse) Assays**

Cells were maintained and treated as described above. When indicated, mitochondrial oxidative phosphorylation from cells was analyzed using extracellular flux analysis (XF24; Seahorse Biosciences) in DMEM buffer (pH 7.4; Sigma Aldrich). Baseline oxygen consumption rates (OCR) were measured every 7 min for control and cells in different biological conditions as previously described. Following baseline measurements, oligomycin (1 μM), Carbonyl cyanide-4-(trifluoromethoxy)phenylhydrazone (FCCP) (1 μM), and antimycin A (2 μM) were sequentially injected to measure OCR.

**Colony-forming fibroblast assays**

Adipose tissue mesenchymal stem cells (MSCs) were expanded in culture until reaching 70%–80% confluence, harvested using trypsin-EDTA (GIBCO, Invitrogen, Germany), and counted. The cells were then diluted in culture media and plated in 96-well plates at a seeding density of 500 cells per well. Each biological condition was plated in separate 96-well plates. After incubation for 3, 7, and 14 days at 37°C in 5% humidified $CO_2$, the cells were washed with PBS and stained with 0.5% Crystal Violet (Sigma-Aldrich) for 15 minutes at room temperature. Following staining, the cells were washed with PBS, and the number of wells containing visible colonies was counted from three plates (triplicates) for each condition. The data were represented for each time point and biological condition as the average percentage of wells (out of 96) with visible colonies, with results reported in triplicate.

**Intraperitoneal glucose tolerance test (IP-GTT)**

Intraperitoneal GTT was performed on overnight fasted mice. Blood glucose levels were measured at basal state (0 min) and then at 30, and 120 min after i.p. injection of glucose (1.5 mg/kg body weight). Blood glucose concentrations were measured using the Accu-Chek Aviva monitoring system (Roche, Basel, Switzerland).

**Insulin tolerance test (ITT)**

ITT was performed on fed mice. Blood glucose levels were measured at basal state (0 min) and then at 30, and 120 min after i.p. injection of human insulin (1.0 mU/g body weight; Actrapid Penfill, Novo Nordisk, Denmark). Blood glucose concentrations were measured using the Accu-Chek Aviva monitoring system (Roche, Basel, Switzerland).

**Comparison of metabolic phenotypes in mammals**

Data for multi-species comparisons of metabolic phenotype were obtained as follows: total energy expenditure (TEE) from, including hominidae fat and lean mass, and (21, 23), experimental fat mass measurement across mammals (24) (table S1).

**Fasting endurance score (theoretical survival time)**

To calculate energy endurance score, we followed previous work by Lindstedt et al. (22), where they generated a fasting endurance score (theoretical survival time in nutritional deficit) as a function of body size in mammals. We used a similar approach including allometric (10% of body weight) and experimentally measured fat mass. Briefly, the theoretical survival time or endurance score is based on the relationship between fat mass energy content (in kilojoules) and the amount of energy consumed per day in the form of total-energy-expenditure (TEE) collected from previous reports and shown in table S2. This measure represents resource availability, assuming energy intake is low. This gives a representation of the number of days a given species can last solely on the energy stored in fat tissue. To calculate the theoretical survival time for each species, we first calculate the energy content in the fat mass (assuming fat mass is 10% of body mass) and then divide that by the TEE (kcal/d) for each species.

Energy content in fat = Fat mass (kg) * 7700 kcal/kg (1 g of fat contains approximately 7.7 kcal)

Theoretical survival time (days) = Energy content in fat (kcal) / TEE (kcal/d)

After obtaining the theoretical survival time or endurance score for each species (wherever data was available), we plot this as a XY graph together with fat mass. Fat mass is derived from both experimentally measured and allometric fat mass. We then performed linear regression to compare both experimental and allometric models, followed by F-test model comparison.

**Somatic mutation and DNA methylation rates**

To calculate energy endurance score, we followed previous work by Lindstedt et al. (22), where they generated a fasting endurance score (theoretical survival time in nutritional deficit) as a function of body size in mammals. We used a similar

approach including allometric (10% of body weight) and experimentally measured fat mass. Briefly, the theoretical survival time or endurance score is based on the relationship between fat mass energy content (in kilojoules) and the amount of energy consumed per day in the form of total-energy-expenditure (TEE) collected from previous reports and shown in table S2. This measure represents resource availability, assuming energy intake is low. This gives a representation of the number of days a given species can last solely on the energy stored in fat tissue. To calculate the theoretical survival time for each species, we first calculate the energy content in the fat mass (assuming fat mass is 10% of body mass) and then divide that by the TEE (kcal/d) for each species.
Energy content in fat = Fat mass (kg) * 7700 kcal/kg (1 g of fat contains approximately 7.7 kcal)
Theoretical survival time (days) = Energy content in fat (kcal) / TEE (kcal/d)
After obtaining the theoretical survival time or endurance score for each species (wherever data was available), we plot this as a XY graph together with fat mass. Fat mass is derived from both experimentally measured and allometric fat mass. We then performed linear regression to compare both experimental and allometric models, followed by F-test model comparison.

**Human accelerated regions (HAR) associated genes**
Human accelerated regions and HAR associated genes were obtained from previous work (*5*) (table S3). Full HAR-associated genes were completed with subsequent studies (*6*). Enhancer coordinates and HAR-genes were used for downstream analysis. HAR-genes were used as input for network-based module discovery in specific tissues, followed by interrogation of overlapping molecular signatures with results from primate-human high-throughput comparisons.

**Identification of genomic hubs by positional gene enrichments (PGE)**
To identify genomic hubs or regions with positional gene enrichments (PGEs) from the location of genes of interest, we utilized previously established methods (*45*). PGEs exploit the topological positioning of genes to determine genomic regions that are enriched in specific gene sets. This approach calculates the hypergeometric distribution along genomic distances to assess the likelihood of gene set enrichment within a particular region. For a given genomic region, the probability of observing a specific set of genes is determined. To assess statistical significance, we used cumulative p-value distributions derived from random simulations or applied a false discovery rate (FDR) to large gene sets, as described in previous studies (*45*). The probability of achieving p-value enrichments that are better than chance estimates are used to define genomic hub enrichments. Additional criteria for defining a genomic region as displaying positional gene enrichment include: having >4 genes;No smaller region with fewer genes was found through random permutations and cumulative p-value distributions; no larger regions with more target genes were found. This algorithm defines genomic regions based on the distance between queried genes, followed by p-value distribution estimations to compare with random expectations. Genomic ranges for positional enrichments were limited to 10 million base pairs, and larger PGEs were filtered out. To obtain qualitative PGEs we used regions that were enriched within cytogenetic bands (*71*). Additionally, we performed random permutations with gene sets containing the same number of input genes, followed by statistical comparisons using a Chi-square test and p-values to confirm the robustness of the positional enrichments.

**Kits and antibodies**
Triglyceride (Abcam ab65336), free fatty acid (ab65341), ROS detection (ab139476), fatty acid oxidation assay (ab211154), hydroxyproline assays (ab222941), DNA damage quantification colorimetric Kit (ab284577), norepinephrine ELISA Kit (ab287789), mouse IL2RB / CD122 (ELISA Kit, LS-F36999, LS-bio)
Nrf2 transcription factor assay kit (ab207223), 8-hydroxy 2 deoxyguanosine ELISA kit (abcam ab201734), global DNA methylation assay kit (5 methylcytosine, ab233486), phospho-Histone H2AX (S139) ELISA kit (Cat #: KCB2288, biotechne), senescence b-galactosidase activity assay (#23833 cell signaling technology), mouse IFN-gamma quantikine ELISA kit (Catalog #: MIF00, biotechne), mouse leptin ELISA Kit (ab199082), plasma insulin levels were quantified using the Crystal Chem Ultra Sensitive Mouse Insulin ELISA kit (Crystal Chem Ultra Sensitive Mouse Insulin Elisa Kit, 90080). Plasma total cholesterol (thermo fisher scientific, infinity total cholesterol reagent, TR13421), triglycerides (thermo fisher scientific, infinity total triglycerides reagent, TT22421), PARP1 enzyme activity assay (Sigma, 17-10149), mouse glutathione (GSH) ELISA kit (mybiosource, MBS267424), annexin V-alexa fluor 488/PI apoptosis detection kit (mybiosource), TNF alpha ELISA kit (legend max mouse kit biolegend, Cat# 430907), Acetyl-coenzyme A assay kit (sigma, MAK039), malondialdehyde lipid peroxidation assay (sigma MAK085), oligonucleotide conjugation kit for proximity ligation assays (ab218260) and qPCR on conjugated products following manufacturer's instructions, NAD/NADH assay kit (ab65348). Antibodies: Anti-NRF2 polyclonal antibody (invitrogen, cat# PA5-27882), CIRBP polyclonal antibody (proteintech, cat# 10209-2-AP), H3K27Ac (abcam, Anti-histone H3 (acetyl K27) antibody, Cat# ab4729), PARP1 polyclonal antibody (invitrogen, cat# PA5-34803), Nurr1/NR4A2 polyclonal antibody (Proteintech, Cat#10975-2-AP), anti-mouse CD4 (RM4-5, thermofisher, cat#48-0042-82), anti-mouse CD45

(thermofisher, cat#25-0453-82), anti-CD44 (BD pharmigen, IM7), anti-mouse/human CD11b (M1/70) (biolegend, cat#101242), anti-CD27 (ebioscience, LG.7F9), anti-Vγ2 (ebioscience, UC3-10A6), anti-TCRδ (biolegend, GL3; 1:200), anti-Vγ1 (biolegend, 2.11), anti-TCRβ (biolegend, H57-597), anti-mouse Foxp3 (thermofisher, cat#11-5773-82), anti-mouse CD8a (thermofisher, cat#11-0081-82), anti-mouse CD3e (thermofisher, cat#HM3420), RORγt (ebioscience, B2D), anti-mouse CD122 (IL-2Rβ) antibody (biolegend, cat#105905), anti-mouse CD19 (thermofisher, cat#47-0193-82), anti-mouse CD127 (IL-7Rα) antibody (biolegend, cat#135002), anti-mouse NK1.1 (thermofisher, cat#25-5941-8), anti-mouse CD19 (thermofisher, cat#47-0193-82), anti-F4/80 (thermofisher, BM8).

**Protein extracts**

Cell cultures were homogenized in lysis buffer (50 mM Tris-HCl pH 7.4, 180 mM NaCl, 1% Triton X-100, 15% glycerol, and 1 mM EDTA) supplemented with 1 mM DTT, 0.5 mM PMSF and phosphatase inhibitor cocktail (Sigma-Aldrich). Protein concentration was determined by Bradford Protein Assay following manufacturer's instructions (BioRad).

**Nuclear extraction**

Cells were collected (5 x 106) in PBS by centrifugation (non-adherent) or by scraping from culture flasks (adherent). Cells were washed twice with cold PBS, and the pellet was collected. Cells were transferred into a prechilled microcentrifuge tube, and resuspended in 500 μL 1X Hypotonic Buffer (20 mM Tris-HCl, pH 7.4; 10 mM NaCl; 3 mM MgCl2) by pipetting up and down several times. Cells were incubated on ice for 15 minutes, followed by adding 25 μL detergent (10% NP-40) and vortexing for 10 seconds at highest setting. The homogenate was centrifuged for 10 minutes at 3,000 rpm at 4°C, and the supernatant was transferred (contains the cytoplasmic fraction). The pellet, containing the nuclear fraction, was resuspended in 50 μL complete cell extraction buffer (10 mM Tris pH 7.4, 2 mM Na3VO4, 100 mM NaCl, 1% Triton X-100, 1 mM EDTA, 10% glycerol, 1 mM EGTA, 0.1% SDS, 1 mM NaF, 0.5% deoxycholate, 20 mM Na4P2O7, 1mM PMSF, and protease inhibitor cocktail) for 30 minutes on ice with vortexing at 10 minute internals. Cells were centrifuged for 30 minutes at 14,000 x g at 4°C and the supernatant was transferred (nuclear fraction) to a clean microcentrifuge tube. Aliquots were made and stored at –80°C for downstream analysis.

**Cytotoxicity assay**

NK cells were co-cultured with AT-MSCs extracted from the aged-HFD group at different ratios (1:1 or 10:1) for 4 hours at 37°C. NK cell cytotoxicity was assessed using the CytoTox-Fluor Cytotoxicity Assay (Promega) according to the manufacturer's instructions. For the assay, target cells were labeled with either 1 μM CellTrace Violet (Life Technologies) or 2 μM CellVue Maroon (Ebioscience) and mixed with NK cells at various NK
ratios in NK cell culture medium in a 96-well plate. The cells were centrifuged to synchronize NK conjugate formation and incubated at 37°C for 1 to 2 hours. After incubation, the cells were washed with cold PBS, and the percentage of dead target cells was determined by propidium iodide (PI) staining (Sigma-Aldrich) using flow cytometry.

**Adoptive cell transfer**

Adipose tissue NK cells from 6-12 weeks wild type mice were extracted and maintained as indicated (see mammalian cell culture). These cells are used for ex vivo engineering and subsequently transfer to host mice. A total of 2x10e6 cells/mice were resuspended in 200 ul of PBS and were injected by intraperitoneal injections to the gonadal area of host mice. Recipient mice were as indicated throughout the study. One week before cell transfer, host mice received one dose of 100 mg/kg of cyclophosphamide to facilitate engrafting as previously described (*94*). Each group of host mice received either control or engineered cells (muen-1 or muen-2 NK cells), followed by interventions and observation as indicated throughout the study. 2, 4, or 12 weeks later, mice were subjected to metabolic and molecular analysis as previously described.

**Flow cytometry**

Excluding dead cells was performed with the fixable viability stain or Zombie Aqua (BioLegend). Cells were fixed for 20 min at room temperature and incubated with antibodies for 30-45 min. iNKT cells were identified as live, single lymphocytes binding to anti-TCRβ or anti-CD3 antibodies and αGalCer analog PBS57-loaded CD1d tetramer (NIH Tetramer Core Facility/Emory Vaccine Center). For analysis of iNKT cells and NK cells, a "dump" channel with antibodies against CD19 and F4/80 was used to eliminate nonspecific staining. Lymphocytes were subsetted based on CD56 and CD3 expression, identifying CD56+ CD3- NK cells and CD56+/-, CD3+ T cells. iNKT were identified using CD3+ and CD1d tetramer loaded with a glycolipid (α-GalCer). γδ T cell subsets were identified based on the expression of TCR Vδ1, with CD4+ and CD8+ T cells were gated accordingly. Flow cytometry analysis was performed with a FACS Fortessa, LSRII or Canto II using FACS Software (BD Biosciences) and data analyzed using FlowJo software (BD Biosciences).

**Metabolic Parameters**
Mice were acclimated during 24 h in single cages. After this period mice were monitored for 2-7 days (when indicated) using CLAMS (Comprehensive Laboratory Animal Monitoring System; Columbus Instruments and promethion metabolic cage system) with access to food ad libitum. The following parameters were monitored continuously: Food intake, drinking volume, O2 consumption (VO2), CO2 production (VCO2), heat and locomotion.

**Brain Microdissection**
Extracted brains were rapidly removed, freshly frozen, and immediately stored at −80°C until use. Brains were sectioned into 500 μm slices using a cryostat and dissected according to previously established protocols (*95*, *96*). Microdissection of specific brain regions such as the hypothalamus, was performed using punchers of various sizes (0.75–2 mm, Harris UniCore, Ted Pella, Inc., Redding, California). Anatomical regions were identified based on the brain atlas by Paxinos and Watson (*95*).

**Weight loss intervention in humans**
All patient samples used in this study were collected from St Columchille's Hospital, Loughlinstown in collaboration with Dr. Johan Meurling and Dr. Donal O'Shea. Obese patients were self-admitted to St. Columchille's Hospital, where they underwent a 6 week long dietary restriction program (1100 calories daily). Patients are not allowed to leave the hospital grounds for 6 weeks while participating in the program. They get six servings of full fat milk daily in addition to Multibionta (multi vitamins, minerals and biotic culture) and are encouraged to drink at least 2 L of water each day. It is unusual for the patients to feel hungry or weak. Patients see a physiotherapist once a week and are encouraged to attend an exercise class on Fridays. Regarding their daily activity, patients aim to reach 8000 steps a day but this depends on the patient's own ability. Blood was taken upon admission (week 0), after 3 weeks and after 6 weeks. PBMCs were isolated for functional analysis of cytotoxic and innate lymphocytes and single cell RNA sequencing. Blood plasma was collected and frozen in liquid nitrogen to analyze plasma metabolite profiles and levels.

**Single cell RNA sequencing data analysis**
Reads were aligned to the mouse reference genome version GRCm38 or human reference genome version GRCh38 and read counts were estimated using Cellranger 3.0.1 (10x Genomics, Pleasanton CA). The Seurat R package v.3.2.0 was used to analyze the generated cell-by-gene unique molecular identifier (UMI) count matrix. We only kept the cells expressing at least 500 genes and genes with expression in at least 50 cells. The cells were also filtered by the maximum of 6000 detected RNA and of 10% mitochondrial genes. DoubleFinder was used to identify the potential doublets in the dataset with the parameter of 7.5% doublet formation rate based on the recommendation of 10x Genomics. The detected potential doublets were removed. The counts were then normalized for each cell by the total number of counts, multiplied by 10000, and log-transformed. We used Seurat's default method to identify top 2000 highly variable genes and scale data for regressing out variation from UMI. The scaled data with variable genes was used to perform principal component analysis (PCA). The top 20 principal components were used for clustering to identify cell populations. UMAPs were calculated in the Seurat package using top 20 PCs and min_dist=0.5. Cell type annotation and marker genes identification:
We used SingleR to estimate the cell type for each cell based on the embedded markerset in the package and determined the cell type for each cluster by evaluating the dominating cell type. We then checked the canonical markers (B cells: MS4A1, CD79A; NK cells: GNLY, NKG7; Naïve CD4+ T cells: IL7R, CCR7; Memory CD4+ T cells: S100A4; T cells: CD3E; Monocytes: CD14, LYZ, FCGR3A, MS4A7; Dendritic cells: FCER1A, CST3; Platelet: PPBP; CD8 T cells: CD8A) in each cluster to confirm the cell type annotation. To better characterize the cell types of NK, NKT and all possible subtypes of T cells, we performed clustering analysis on the cells that might belong to these cell types and annotated the cell types based on the highly expressed genes in each cluster using FinderMarker function in Seurat package, as well as markers identified in previous studies (*97*–*99*). Innateness and adaptiveness analysis: The gene sets for innateness and adaptiveness were identified from Gutierrez-Arcelus et al. (*63*). The AddModuleScore function in Seurat was used to add the innateness and adaptiveness score for each cell. According to the score, we identified adaptiveness/innateness associated genes using a linear model, smoothed the gene expression along the score by fitting a smoothing spline in R and visualized the gene expression patterns with hierarchical clustering. Differential gene expression analysis: The differentially expressed genes were detected using Wilcoxon rank sum test (the default parameters in Seurat) by comparing the cell types between conditions. Functional enrichment analysis: Enrichr in R was used to perform functional enrichment analysis based on the database: Gene Ontology and FDR <0.01 was used as a threshold to select the significant enrichment. The tabula muris project (*61*) data was used to interrogate expression levels of key genes in adipose tissue cells.

**Targeted metabolomics**

Metabolites were analyzed using a liquid chromatography-mass spectrometry (LC-MS) system consisting of a Nexera X2 U-HPLC (Shimadzu Scientific Instruments) coupled with a Q Exactive-HF-X Orbitrap mass spectrometer (Thermo Fisher Scientific). Samples were promptly isolated and homogenized in an extraction solution containing 80% methanol with the addition of internal standards (inosine-15N4, thymine-d4, and glycocholate-d4, Cambridge Isotope Laboratories). A 30 µl aliquot of the homogenate was then diluted with 120 µl of the same extraction solution. The samples were centrifuged at 9,000g for 10 minutes at 4°C, and the resulting supernatants were injected directly onto a 150 × 2.0 mm Luna NH2 column (Phenomenex). Elution was carried out at a flow rate of 400 µl/min, starting with 10% mobile phase A (20 mM ammonium acetate and 20 mM ammonium hydroxide in water) and 90% mobile phase B (10 mM ammonium hydroxide in 75:25 v/v acetonitrile/methanol), followed by a 10-minute linear gradient to 100% mobile phase A. Mass spectrometry analyses were performed using electrospray ionization in negative ion mode, scanning over an m/z range of 60–750 at a resolution of 70,000 and a data acquisition rate of 3 Hz. The additional parameters used for mass spectrometry were as follows: ion spray voltage at −3.0 kV, capillary temperature at 350°C, probe heater temperature at 325°C, sheath gas at 55, auxiliary gas at 10, and S-lens RF level at 40. Raw data were processed using Progenesis QI software (NonLinear Dynamics) for feature alignment, non-targeted signal detection, and signal integration. Targeted analysis of known metabolites and isotopologues was conducted using TraceFinder software (Thermo Fisher Scientific), and compound identities were confirmed by comparison with reference standards. Once samples were collected and the metabolites were identified, traditional univariate methods were used for statistical comparison between groups, including normalization of intensities by control groups, assessment of fold-change variation in other groups, followed by t-test analysis, and volcano plot generation.

**Transcriptomic and metabolomics**

We analyzed scRNA-transcriptomics (differentially expressed genes) and metabolomics data (differentially changed metabolites) from our cohorts using the metaboanalyst package (found at https://www.metaboanalyst.ca/). This package is a platform for metabolomics analysis, including joint pathway analysis using transcriptional profiles. It can be used online or through API access using the R package MetaboAnalystR. We used it to evaluate metabolite set enrichment analysis (MSEA), pathway set enrichment analysis (PSEA), joint PSEA, and network analysis using KEGG ortholog data from metagenomics, and downstream TFBS inference from enriched ontologies.

**Network-based module discovery in pre-built tissue-specific networks**

To assess functional gene dependencies in tissue-specific networks, we performed a targeted approach of overlapping network representation using data from tissue-integrated genome-scale analysis (*62*) (archived in http://giant.princeton.edu and https://hb.flatironinstitute.org/). This was followed by evaluating functional enrichments of associated networks.

**Data, Code Availability**

Analyses were performed in R (3.5 and 3.6) and python (2.7, 3.5 and 3.7). Libraries for data analysis include numpy, pandas, scikit-learn, scanpy and scipy. The datasets reported in this paper are available in supplementary materials. Below is detailed data and source to different datasets used in this paper. Tissue-specific networks for functional annotation of topological dependencies can be found at http://giant.princeton.edu and https://hb.flatironinstitute.org/. Positional gene enrichment analysis tool can be found at https://github.com/leandroagudelo189/Metabolic-resilience-topo-genomics/Positional-gene-enrichment. Functional annotation tool from a large library of biological repositories can be found at https://maayanlab.cloud/Enrichr/. Prediction of TFBS from DNA sequences can be found at http://alggen.lsi.upc.es/. Prediction of conserved TFBS in promoter regions of genes of interest can be found at http://www.gsea-msigdb.org/gsea/msigdb/index.jsp. PPIs reconstruction can be found at http://www.interactome-atlas.org/ and at https://thebiogrid.org/. Cis-regulatory elements relationships were obtained from GeneHancer, and can be found at http://www.genecards.org/. A curated database of cell-specific active regulatory regions can be found at http://www.enhanceratlas.org/index.php. For topological comparative genomics and to evaluate chromosomal synteny, tools can be found at the synteny portal, archived in http://bioinfo.konkuk.ac.kr/synteny_portal/.

**Statistical Analysis**

Statistical analyses related to experimental procedures were performed using GraphPad Prism Software (San Diego, CA) and all parameters are indicated in the corresponding figure legend. Quantitative data are presented as the mean ± SEM and n is indicated for each experiment. All experiments were carried out with 3 biological replicates and 3 independent times. Animal experiments were performed in more than 3 animals per group, both female and males were used when indicated. Unpaired Student's t test was used to determine statistical significance when two groups were compared. One-way ANOVA followed by

Fisher's least significant difference (LSD) test for post hoc comparisons was used to determine statistical significance when multiple groups were compared. Two-way ANOVA was used to estimate significance between groups constrained by time-based or concentration-based measurements, followed by Tukey test for post-hoc multiple comparisons. Hypergeometric test enrichments and Chi-square tests were used to determine the statistical significance between expected and observed frequencies. Correlations were calculated by Spearman rank and Pearson correlation with a two-tailed test. Statistical significance was defined as p value < 0.05 by either test and is denoted with an asterisk as follows: * p-value 0.05-0.01, ** p-value 0.01-0.001 and *** < 0.001 when indicated.

**Disclosure on the use of generative AI:** During the preparation of this work, the authors used the assistance of LLMs such as GPT5, Claude, Gemini pro to edit and improve readability and grammar. All suggestions of text edits were edited, reviewed, validated, and manually corrected by the authors prior to submission of this work. The authors take full responsibility for the originality and content of this work. In addition, during preparation of specific parts, the authors used the same models for code assistance. The authors reviewed and edited the content as needed and take responsibility for the content of this work.
